# CTCF binding landscape is shaped by the epigenetic state of the N-terminal nucleosome in relation to CTCF motif orientation

**DOI:** 10.1101/2024.09.25.614770

**Authors:** Md Tajmul, Dharmendra Nath Bhatt, Luminita Ruje, Emma Price, Yon Ji, Dmitri Loukinov, Vladimir B. Teif, Victor V. Lobanenkov, Elena M. Pugacheva

## Abstract

CTCF binding sites serve as anchors for the 3D chromatin architecture in vertebrates. The functionality of these anchors is influenced by the residence time of CTCF on chromatin, which is determined by its binding affinity and its interactions with nucleosomes and other chromatin-associated factors. In this study, we demonstrate that CTCF occupancy is driven by CTCF motifs, strategically positioned at the entry sides of a well-positioned nucleosome, such that, upon binding, the N-terminus of CTCF is oriented towards the nucleosome. We refer to this nucleosome as the CTCF priming nucleosome (CPN). CTCF recognizes its binding sites if they are not methylated. It can then displace the CPN, provided the nucleosome is not marked by CpG methylation or repressive histone modifications. Under these permissive conditions, the N-terminus of CTCF recruits SMARCA5 to reposition the CPN downstream, thereby creating nucleosome-free regions that enhance CTCF occupancy and cohesin stalling. In contrast, when CPNs carry repressive epigenetic marks, CTCF binding is transient, without nucleosome displacement or chromatin opening. In such cases, cohesin is not effectively retained at CTCF binding sites. We propose that the epigenetic status of CPNs shapes cell-specific CTCF binding patterns, ensuring the maintenance of chromatin architecture throughout the cell cycle.

**Graphical Abstract:** 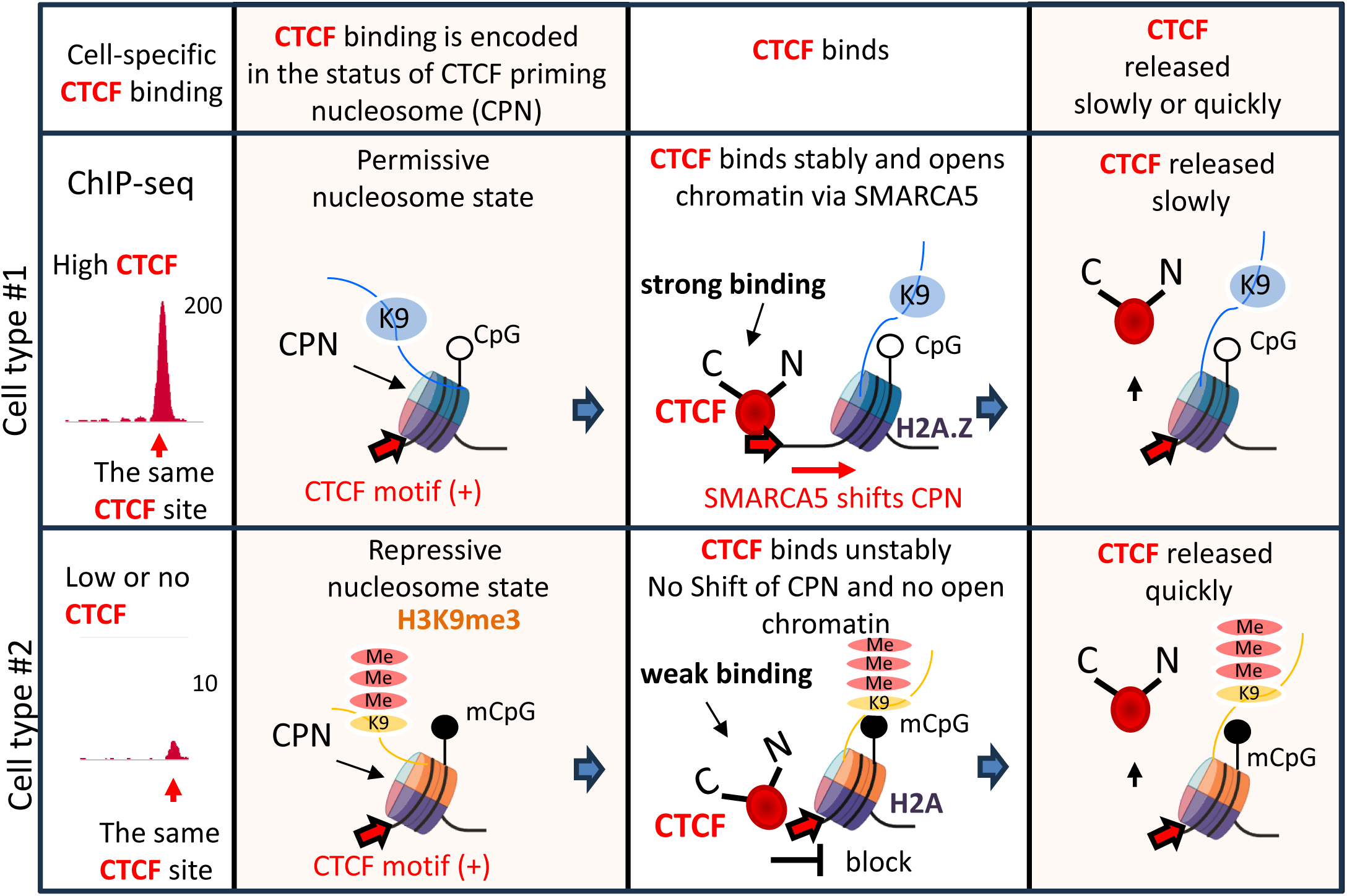

## Introduction

3D genome organization in vertebrates is largely shaped by CTCF, which stops cohesin-guided extrusion of chromatin at its binding sites in the convergent orientation (1,2). CTCF is a ubiquitously expressed, highly conserved 11 Zinc Finger (ZF)-DNA binding factor (3). As part of the ENCODE project, CTCF binding sites were among the first to be mapped by ChIP-seq across multiple tissues and species (4). It is estimated that there are approximately 326,000 CTCF binding sites in the human genome, with an average of 66,000 sites occupied by CTCF in a single cell type (5). CTCF binding sites serve as anchors for 3D genome organization (2), activators or repressors of transcription (6), insulators (7), imprinting control regions (8), DNA repair sites (9), and more (10). Dysfunction of CTCF binding sites has been associated with various pathological conditions, including myotonic dystrophy (11), aberrant X-chromosome inactivation (12), neurodevelopmental disorders (13), and cancer (14,15).

Although the overall CTCF binding pattern remains largely consistent across different cell types, a small but significant proportion (∼10-30%) of CTCF binding sites are cell type-specific (16). These cell-specific sites play a critical role in defining cell identity by contributing to cell-specific 3D genome organization and gene transcription (17). For instance, during ES cell differentiation into distinct lineages, changes in CTCF occupancy—marked by the loss and gain of cell-specific sites—are closely linked to cell specialization (18). Similarly, unique cancer-specific CTCF binding patterns have been identified across various human malignancies (14), and anti-cancer drug treatments have been shown to alter CTCF occupancy (19,20). Therefore, CTCF binding patterns appear to be fundamental to cell fate and specialization. The dynamic reorganization of CTCF occupancy may represent a crucial cellular mechanism for adapting to new conditions or environmental challenges. Nevertheless, key questions remain underexplored: What determines the specificity of CTCF binding in different cell types, and how are changes in CTCF occupancy achieved during processes such as development, differentiation, or stress responses?

It has been established that CTCF occupancy is regulated by CpG methylation (21–23), cell-specific transcription factors (TFs) (17,24), kinases that phosphorylate ZFs to displace CTCF during mitosis (25) or in hormone response (26), chromatin remodeling factors that open chromatin for CTCF occupancy (27), histone marks (28) and more (29). CTCF binds a great assortment of DNA sequences using combinatorial clustering of its 11 ZFs (30,31). CTCF ZFs 3-7 are responsible for recognition and binding the 15-bp CTCF core consensus (30), while ZFs 1-2 and ZFs 9-11 bind downstream and upstream from the core consensus sequences, respectively (32). CTCF bound sites are located in nucleosome-depleted regions (NDRs) flanked by several well-positioned nucleosomes around CTCF binding sites (33). As cohesin translocates along chromatin, it encounters nucleosomes before reaching CTCF bound at convergent motifs, suggesting that nucleosomes may act as “regulatory barriers,” potentially modulating cohesin’s translocation speed and adding an additional layer of regulation to cohesin dynamics and loop extrusion.

Single-molecule level analysis of chromatin structure has demonstrated that only a small fraction of individual fibers is bound by CTCF at any given time in a cell population (34). Similar conclusions about the transience of CTCF binding were drawn from single-cell nucleosome positioning (35) and single-cell DNA-footprint (36). CTCF binding is dynamic, with CTCF protein binding on and off chromatin with an average residence time of 1-2 minutes (37,38), while the residence time of CTCF-anchored loops is approximately 10 minutes (39). Such dynamic binding is essential for 3D genome organization and necessary for chromatin extrusion by cohesin (16). When CTCF is off chromatin, its place is occupied by a nucleosome (36,40,41). To re-bind to chromatin, CTCF must first reposition this nucleosome. This process involves chromatin-remodeling factors such as SNF2H (SMARCA5), CDH4, and BRG1, which can slide, evict, or remove nucleosomes around CTCF binding sites (27,42,43). Consequently, nucleosomes play a crucial role in regulating CTCF occupancy. Evidence supporting this includes the significant loss of CTCF occupancy observed after the loss of SMARCA5, which disrupts CTCF’s ability to reposition nucleosomes and re-bind to its target sites (27,44,45).

The CTCF-nucleosome regulation model suggests that CTCF can recognize and bind to its sites even when they are occupied by nucleosomes (36). Moreover, CTCF restores the exact same binding pattern in daughter cells after mitosis, during which most CTCF is released from chromatin due to phosphorylation of CTCF zinc fingers (25,46). Additionally, when CTCF is degraded using auxin-degron systems, its occupancy is erased, but upon auxin removal, CTCF occupancy is restored exactly as it was before (47). These observations suggest that the CTCF binding pattern for each cell type is encoded in the DNA and chromatin status, thereby maintaining cell identity.

In this study, we sought to identify how chromatin status is involved in a cell-specific CTCF occupancy. By comparing ChIP-seq and Affinity-seq data from the same cell type, we analyzed CTCF binding sites both *in vivo* and *in vitro*. Based on our findings, we propose a model for the regulation of CTCF occupancy that is driven by the epigenetic status of the nucleosome that is well-positioned relative to CTCF binding sites. To confirm this proposed model, we used wild type (wt) and mutant (mut) CH12 cells. Compared to wtCH12 cells, a homozygous deletion of CTCF ZF 9-11 in mutCH12 cells resulted in a loss of CTCF occupancy at five thousand (5K) binding sites (48,49). Ectopic expression of wild type CTCF in the mutCH12 cells not only recapitulated CTCF occupancy like in wild type cells but, most importantly, restored CTCF occupancy at the same 5K lost CTCF sites (48), suggesting that these sites are somehow pre-marked to be bound by CTCF in the CH12 cells. In this study, we demonstrate that cell-specific CTCF occupancy is preprogrammed not only by the status of CpG methylation in the CTCF site itself, but also by the status of CpG methylation and histone modifications of the nucleosome asymmetrically positioned relative to CTCF binding site.

## Materials and methods

### Cell culture, plasmids

CH12 wild type (wt) and mutant (mut) cells were described previously in (48,49) and obtained as a gift from Dr. Casellas’s lab. The cell lines were maintained in RPMI 1640 supplemented with 10% FBS, 1% penicillin/streptomycin and 55 μM 2-β mercaptoethanol. All ectopically expressed CTCF and BORIS constructs described in this study were designed on a backbone of pMy-MouseCTCFbiotag-T2A-mOrange plasmid (Addgene plasmid #50564) and described in (48). NIH3T3, stably expressing either EV (Empty Vector) or BORIS (clone#2), were maintained in Dulbecco’s modified Eagle’s medium (DMEM) supplemented with 10% of fetal calf serum and penicillin–streptomycin, described in (24).

### ChIP-seq

For ChIP-seq, 2×10^6^ asynchronously growing cells were crosslinked with 1% formaldehyde for 10 minutes at room temperature, followed by quenching with 125 mM glycine for 10 minutes. Cells were then washed twice with 1× phosphate-buffered saline (PBS) and resuspended in chromatin immunoprecipitation (ChIP) lysis buffer (150 mM NaCl, 1% Triton X-100, 0.1% SDS, 20 mM Tris-HCl pH 8.0, 2 mM EDTA). Chromatin was sheared to 200–600 bp and immunoprecipitated using 5 μg of respective antibodies. The immunoprecipitated DNA was purified using Zymo-spin^TM^ columns (ZymoResearch), and DNA concentration was measured with a Qubit 4 Fluorometer (ThermoFisher). For sequencing, 5–10 ng of DNA was used to prepare libraries with the NEBNext Ultra II DNA Library Prep Kit for Illumina (New England, Catalog E7645S). Paired-end sequencing was performed on a NovaSeq 6000 system. All ChIP-seq experiments were conducted with at least two biological and/or technical replicates. Antibodies used in ChIP-seq: SMARCA5 (Snf2h) (Abcam, #ab3749), H3K9me3 (Abcam, ab#8898), RAD21 (Abcam, ab217678), SMC3 (Abcam, ab#9263), SMC1 (Bethyl Laboratories, #A300-055A).

### ATAC-seq

Freshly growing CH12 cells (5 × 10^4^) were pelleted by centrifugation and resuspended in 50 µL of cold lysis buffer (10 mM Tris-HCl, pH 7.4, 10 mM NaCl, 3 mM MgCl_2_, 0.1% IGEPAL CA-630), gently pipetting up and down ten times. Nuclei were pelleted by centrifugation at 2500g for 10 min at 4°C and resuspended in 25 µL of 2x TD Buffer (Illumina Cat #FC-121-1030), containing 8 µL of Tn5 Transposes (Illumina Cat #FC-121-1030) and 17 µL of nuclease-free water. The reaction was incubated for 30 min at 37°C. Tagmented DNA was purified by using a Zymo Research DNA Clean & Concentrator™-5 - Capped Columns. The libraries were amplified using NEBNext High-Fidelity 2× PCR Master Mix (New England Biolabs, M0541). The libraries were size-selected by adding 1.8X volume Agencourt AMPure XP beads (Beckman, 63881). Library concentration was measured by DNA High Sensitivity Kit (Invitrogen) on a Qubit fluorometer (Invitrogen). Library quality and fragment sizes were assessed on a Bioanalyzer (Agilent). ATAC-Seq libraries from at least two biological replicates for each condition were paired-end-sequenced on an Illumina NextSeq550 platform. ATAC-seq libraries were generated as previously described (50).

### MNase-Seq Assisted H3 ChIP-seq

MNase-ChIP-seq was performed on wtCH12 or mutCH12 fixed cells using the SimpleChIP® Enzymatic Chromatin IP Kit (Magnetic Beads) (Cell Signaling Technology, Cat #9003) following the manufacturer’s protocol. Briefly, after obtaining MNase-digested lysates as described in the MNase-seq method, chromatin lysates were incubated with 5 μg of Histone H3 (Cell Signaling, #D2B12) XP® Rabbit mAb and rotated overnight at 4°C. Following this incubation, 30 µL of Protein G Magnetic Beads were added to the mixture and rotated for 2 hours at 4°C. The Protein G magnetic beads were then pelleted using a magnetic separation rack for 1–2 minutes. The supernatant was carefully removed, and the beads were washed three times with 1 mL of low salt wash buffer, incubating each wash for 5 minutes at 4°C with rotation. After the final low salt wash, 1 mL of high salt wash buffer was added, and the beads were incubated again for 5 minutes at 4°C with rotation. The beads were pelleted, and the supernatant was removed. Next, 150 µL of 1X ChIP Elution Buffer was added, and the chromatin was eluted from the antibody/Protein G magnetic beads by placing the tubes on a thermomixer at 65°C for 30 minutes with gentle shaking (1,000 rpm). The eluted chromatin supernatant was transferred to a new tube, and the immunoprecipitated DNA was purified according to the kit protocol. Sequencing libraries were prepared using the NEBNext® Ultra™ II DNA Library Prep Kit (New England, Catalog E7645S).

### MeDIP Library Preparation

Genomic DNA (gDNA) was extracted from freshly growing CH12 cells and sonicated using a Covaris ultrasonicator to obtain a mean fragment size of 200 bp. For library preparation, 5 μg of sonicated gDNA was end-repaired and ligated with adaptors using the NEBNext Ultra DNA Library Prep Kit for Illumina (New England, Catalog E7645S). The reaction mixture was then purified with AMPure XP beads (Beckman-Coulter, CA, USA, Catalog #A63881). For precise size selection, half of the sample was run on a 1.5% agarose gel, and DNA fragments of 150-250 bp were extracted from the gel. Both the size-selected and non-size-selected libraries were denatured at 95°C for 10 minutes, followed by incubation on ice for 10 minutes. The denatured DNA was subjected to immunoprecipitation with 5-Methylcytosine antibody (Active Motif, Catalog #61479) by incubating the reaction mixture at 4°C for 15 hours. The immunoprecipitated DNA was size-selected again using AMPure XP beads and amplified using Q5 High-Fidelity DNA Polymerase (New England Biolabs, Catalog #M0491). The amplification products were purified with AMPure XP beads. Finally, the purity and fragment length distribution were assessed using a Bioanalyzer 2100 (Agilent Technologies), and the libraries were subjected to deep sequencing on the Illumina HiSeq 2000 platform.

### Published Next-Generation Sequencing (NGS) datasets used in this study

For the Affinity-seq mouse spleen data, we used the dataset GSE111772, generated in (51). For CTCF ChIP-seq, we used the following datasets: mouse spleen GSM918745, GSM918763 (52); CH12 cells (GSE137216), NIH3T3 cells (GSE207058) from (48), mouse ES cells (SRR5085156, SRR5085157) (37), MCF7 cells (GSE70764), K562 cells (GSE137216) from (24). For RAD21 ChIP-seq, we used the following datasets: CH12 cells (GSE137216) (48), NIH3T3 (current study), mES cells (SRR5085154, SRR5085155) (53). To generate Figure 1D-E and Supplementary Figures S3A, S4C, S6A we used: MNase-H3 ChIP-seq (current study), H2A.Z ChIP-seq (GSE207058) (24), H3K9me3 ChIP-seq (GSM946548, SRR507879) from (52), and 5mC MeDIP-seq (GSM1368909) from (54). To generate Supplementary Figure S2, we used the following NGS data for mES cells: H3K9me3 (SRR21782395), MNase-H3 (GSE146082) (55), 5mC MeDIP-seq (SRR6321573) (56). To generate Figure 2, we used CTCF ChIP-seq, MNase-seq, and H3K9me3 ChIP-seq data from parental and Setdb1 knockout mouse ES cells, as published in (28) and available under GSE184471. To generate Supplementary Figure S3C,D, we used data from (57). To generate Supplementary Figure S5A,B, we used the following NGS data: for MCF7 cells: MNase-seq (SRR3142031, SRR3142029); for K562 cells: MNase-seq (SRR3211681, SRR3211680), 5mC MeDIP-seq (SRR1237957). To generate Supplementary Figure S5D, we used H3K9me3 ChIP-seq for mouse spleen (SRR30337758) from (58). For SMARCA5 depletion data in Figure 5 and Supplementary Figure S8, we used datasets from (27,44,45). For ATAC-seq data in Figure 6 and Supplementary Figure S10, we used the following NGS data: mouse ES cells (SRR14701291) (59), NIH3T3 (GSE207058) (24), CH12 cells (current study). For wtCH12 and mutCH12 cells, we utilized ChIP-seq dataset GSE137216 (for CTCF, BORIS, FL-CTCF, deltaC-CTCF, and deltaN-CTCF), which were generated in our previous study (48). For NIH3T3 cells, we used ChIP-seq dataset GSE207058 that we previously generated by our lab (24). The genomic coordinates (mm9, hg19) are provided in Table S1.

**Figure 1.**
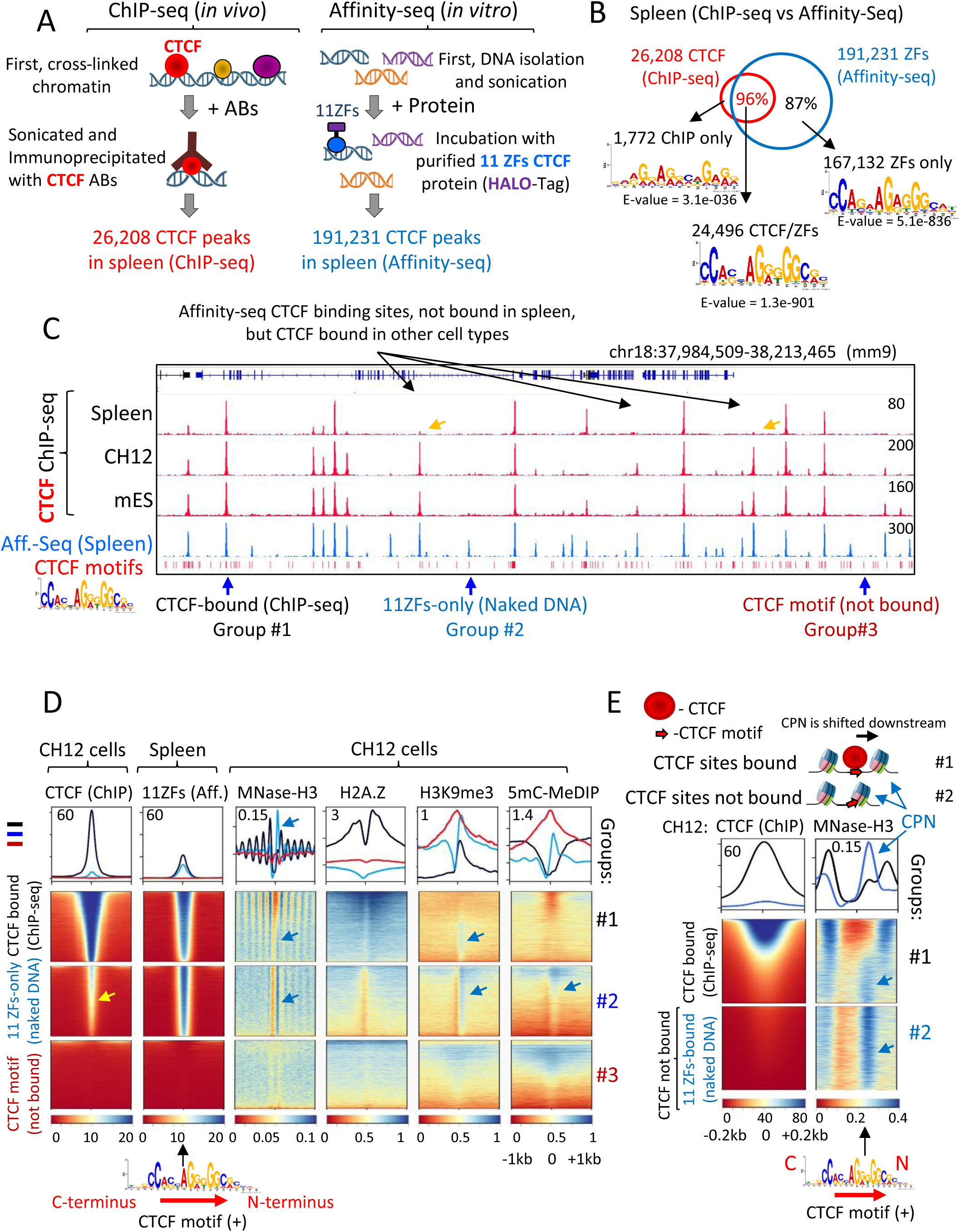
CTCF binding sites are strategically positioned at the entry side of well-positioned nucleosome. **(A)** Schematic comparison of ChIP-seq and Affinity-seq approaches used to map CTCF binding sites in mouse spleen. **(B)** Venn diagram illustrating the overlap between CTCF binding sites identified by ChIP-seq and Affinity-seq. MEME motifs enriched in each group of CTCF sites are indicated by black arrows. **(C)** Representative genome browser view displaying CTCF ChIP-seq data (red) compared to CTCF Affinity-seq data (Aff.-seq, blue). The red bars at the bottom show the 14bp-CTCF motif sequences present in the mouse genome (mm9). The yellow arrows indicate CTCF sites that were not detected by ChIP-seq in the spleen but were identified as CTCF-bound in Affinity-seq data from the spleen. Blue arrows highlight examples of CTCF binding sites categorized into three groups, as analyzed in panel **(D). (D)** Heatmaps centered on the CTCF motif in the plus (sense) orientation, showing (from left to right): CTCF ChIP-seq (CH12 cells), CTCF Affinity-seq (spleen), MNase-H3 nucleosome positioning (CH12), H3K9me3 ChIP-seq (CH12), and 5mC MeDIP-seq data (CH12 cells). These data are aligned across three groups of CTCF binding sites: (#1) CTCF-bound sites in CH12 cells (black), (#2) Affinity-seq recovered CTCF sites that were not identified as significant peaks by MACS in CTCF ChIP-seq of CH12 cells (blue), and (#3) CTCF motifs neither bound by CTCF nor recovered by Affinity-seq (red). The CTCF priming nucleosome (CPN) is indicated by a blue arrow. The low CTCF ChIP-seq signal at the 11 ZFs-only group (#2) is shown by a yellow arrow. **(E)** A zoomed-in view of data from the panel **(D)**, focusing on a 400 bp window centered on the plus CTCF motif. This panel combines CTCF ChIP-seq data with MNase-H3 data from CH12 cells to illustrate the downstream shift of well-positioned nucleosomes from the CTCF-bound motifs. At CTCF motifs recovered by Affinity-seq only, this nucleosome is positioned at the entry side of the plus CTCF motif (indicated by a blue arrow). A schematic representation of bound (#1) and unbound (#2) CTCF sites is provided at the top.

**Figure 2.**
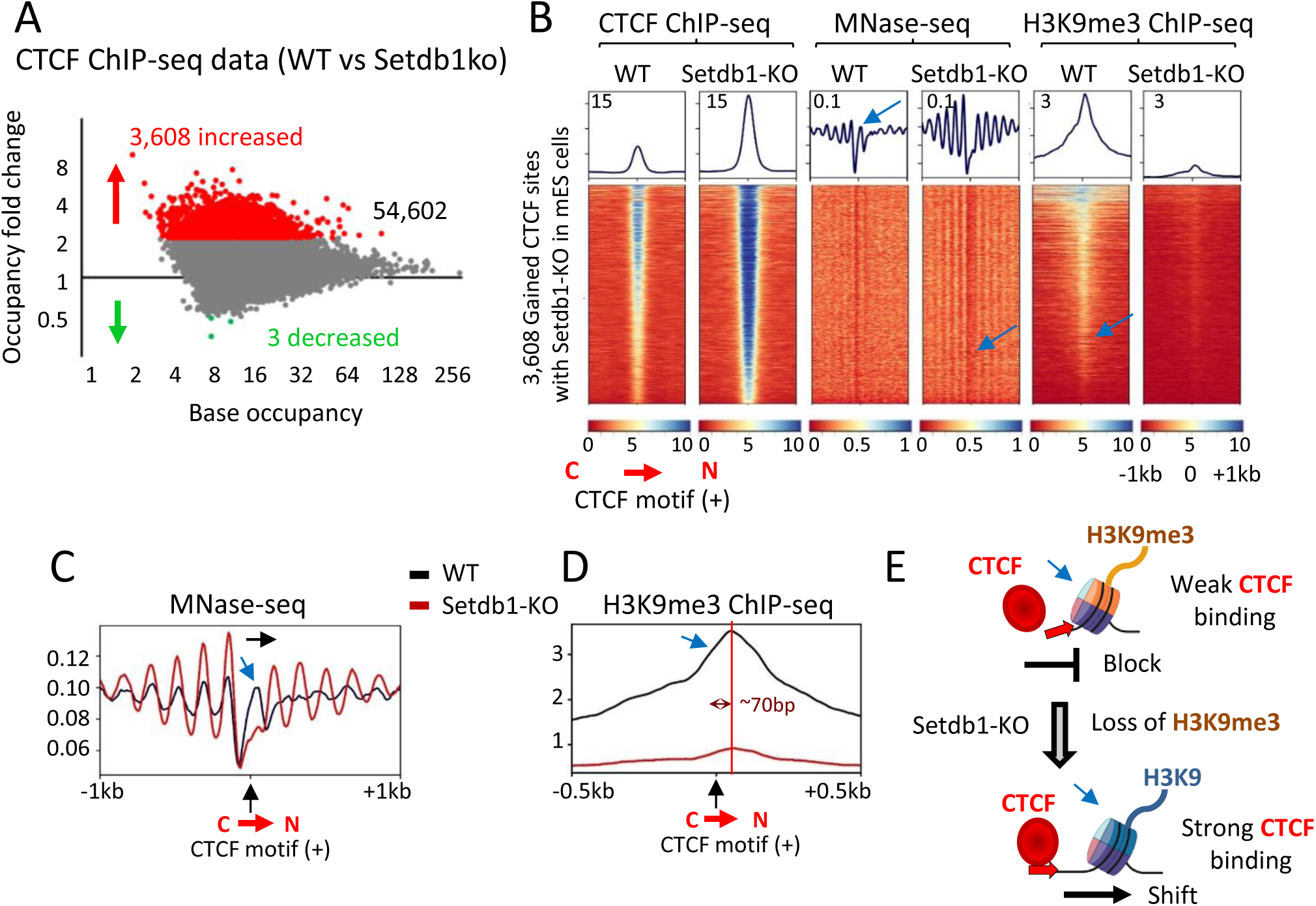
The loss of H3K9me3 at the CTCF priming nucleosome facilitates stable CTCF occupancy by removing repressive chromatin barriers, allowing for stronger binding. **(A**) MA plot showing CTCF occupancy (tag density) in wild-type (WT) versus Setdb1 knockout (KO) mES cells. CTCF peaks are categorized, based on tag enrichment, as unchanged (grey), upregulated (red), or downregulated (green) in Setdb1-KO cells, compared to WT. Axes are displayed in log₂ scale. **(B)** Heatmap centered on the CTCF motif in the plus orientation for 3,608 gained CTCF sites identified in panel **(A).** The heatmap presents a comparative analysis of CTCF ChIP-seq, MNase-seq, and H3K9me3 ChIP-seq data at CTCF sites that gained occupancy in Setdb1-KO cells compared to WT cells. **(C, D)** Profiles derived from panel **(B)** comparing nucleosome occupancy and histone modification enrichment at the 3,608 gained CTCF sites in WT versus Setdb1-KO cells: **(C)** MNase-seq profile showing nucleosome positioning around CTCF motif in plus orientation. The potential shift of the CPN upon Setdb1-KO is indicated by a black arrow. **(D)** H3K9me3 ChIP-seq profile showing enrichment of H3K9me3 at the CPN. The position of the CPN relative to the beginning of the 14 bp CTCF motif in the plus orientation is indicated by a double-ended arrow and red line. **(E)** Schematic representation summarizing the conclusions: the loss of H3K9me3 at the CPN enhances CTCF binding stability. **(B–E)** The CTCF priming nucleosome (CPN) is indicated by a blue arrow.

### Bioinformatic analysis of ChIP-seq, ATAC-seq, MNase-seq, meDIP-seq data

Sequences generated by the Illumina or NovaSeq 6000 were aligned against either the human (build hg19) or mouse (build mm9) genome using the Bowtie2 program with options “–very-sensitive -X 2000” (60). Bowtie output files (SAM) were converted into BAM files using the “samtools view” option in SAMtools (61), and these BAM files were then used for generating heatmaps with deepTools (62). To call ChIP-seq or meDIP-seq peaks we applied MACS2 peak calling algorithm with defaults parameters (63). The ChIP-seq data were visualized using the Integrative Genomics Viewer (IGV) (64). The peak overlaps between ChIP-seq data sets were determined with the BEDTools Suite, with at least 1 bp overlap (65). For ATAC-seq, the open chromatin regions were called using MACS2 with options: –nomodel – shift -100 – extsize 200 - f BAMPE. For MNase-Seq Assisted H3 ChIP-seq, we produced multiple replicates that exhibited a strong Pearson correlation, typically ranging from 0.75 to 0.95, ensuring robust reproducibility. The best replicates were combined into one set for heatmap visualization using deepTools. The overlapping of genomic coordinates to generate Venn diagrams was performed using Intervene (66).

### Affinity-Seq data

CTCF Affinity-seq data for 3-month-old mouse spleen were generated in the Zuo et al. study (51). In this study, the authors isolated genomic DNA from mouse spleen using the phenol-chloroform extraction method and randomly sheared it to ∼200 bp fragments through sonication (51). This extraction method preserves the DNA CpG methylation status, allowing mapping of CTCF binding only to unmethylated DNA sequences. The genomic DNA was incubated overnight with purified 6HisHALO-tagged CTCF 11-Zinc fingers protein (51). In the current study, we downloaded the FASTQ files for CTCF Affinity-seq data from mouse spleen (GSM3039567) and mouse control DNA (GSM3039568) from the series GSE111772. The FASTQ files were aligned to the mouse genome (mm9 build) using Bowtie2 with the options “– very-sensitive -X 2000.” The resulting Bowtie output files (SAM) were converted into BAM files, which were subsequently used for generating heatmaps with deepTools. To call peaks for Affinity-seq, we applied the MACS2 peak-calling algorithm with default parameters, using mouse control DNA as the input control. The list of Affinity-seq CTCF peaks is provided in Supplementary Table 1.

### Heatmaps for NGS data visualization

The heatmaps of ChIP-Seq, ATAC-seq, MeDIP-seq, MNase-seq-H3 ChIP tag densities for different clusters were generated using deepTools (62). For heatmap visualization of ATAC-seq data, we sorted BAM files of paired-end mapped reads into two categories based on fragment size, using the bamCoverage option from deepTools. For CTCF footprints, we applied the arguments “--Mnase --minFragmentLength 40 --maxFragmentLength 130”, and for nucleosome positioning, we used “--Mnase --minFragmentLength 160 --maxFragmentLength 200”. For heatmap visualization of MNase-assisted H3 ChIP-seq data, we similarly used the bamCoverage option with the --Mnase argument. For ChIP-seq and meDIP-seq, we used just bamCoverage option without an additional argument. All BAM files in this study were normalized using the deepTools option “--normalizeUsing RPKM”. Heatmaps were generated using the computeMatrix option to simultaneously analyze and visualize multiple NGS datasets at the same genomic coordinates. Since certain datasets, such as ChIP-seq, exhibited significantly higher enrichment values compared to MNase-H3 ChIP-seq or MeDIP-seq, we adjusted the heatmap intensities using different --zMax values in plotHeatmap. The heatmaps, each with different zMax values as reflected by the numbers on the average plots and scale bars, were then combined into a single figure. The heatmaps and average plots were centered on either a strand-oriented CTCF consensus or the summits of CTCF ChIP-seq peaks, depending on the analysis. For analyses involving MNase-seq H3 data, we specifically used a strand-oriented CTCF consensus because nucleosome positioning is sensitive to the orientation of the consensus motif. Conversely, for ChIP-seq (Figure 3, 5F, 6C,D), ATAC-seq (Figure 3), Supplementary Figures S1, S6B, S10, we centered the data on the summits of CTCF ChIP-seq peaks. This approach ensured that all CTCF binding events, regardless of motif orientation, were included. It is worth noting that only approximately 70–80% of CTCF ChIP-seq peaks contain a canonical CTCF consensus motif with a p-value <1e-5. By centering analyses on the summits of CTCF ChIP-seq peaks, we were able to account for all CTCF binding events comprehensively.

**Figure 3.**
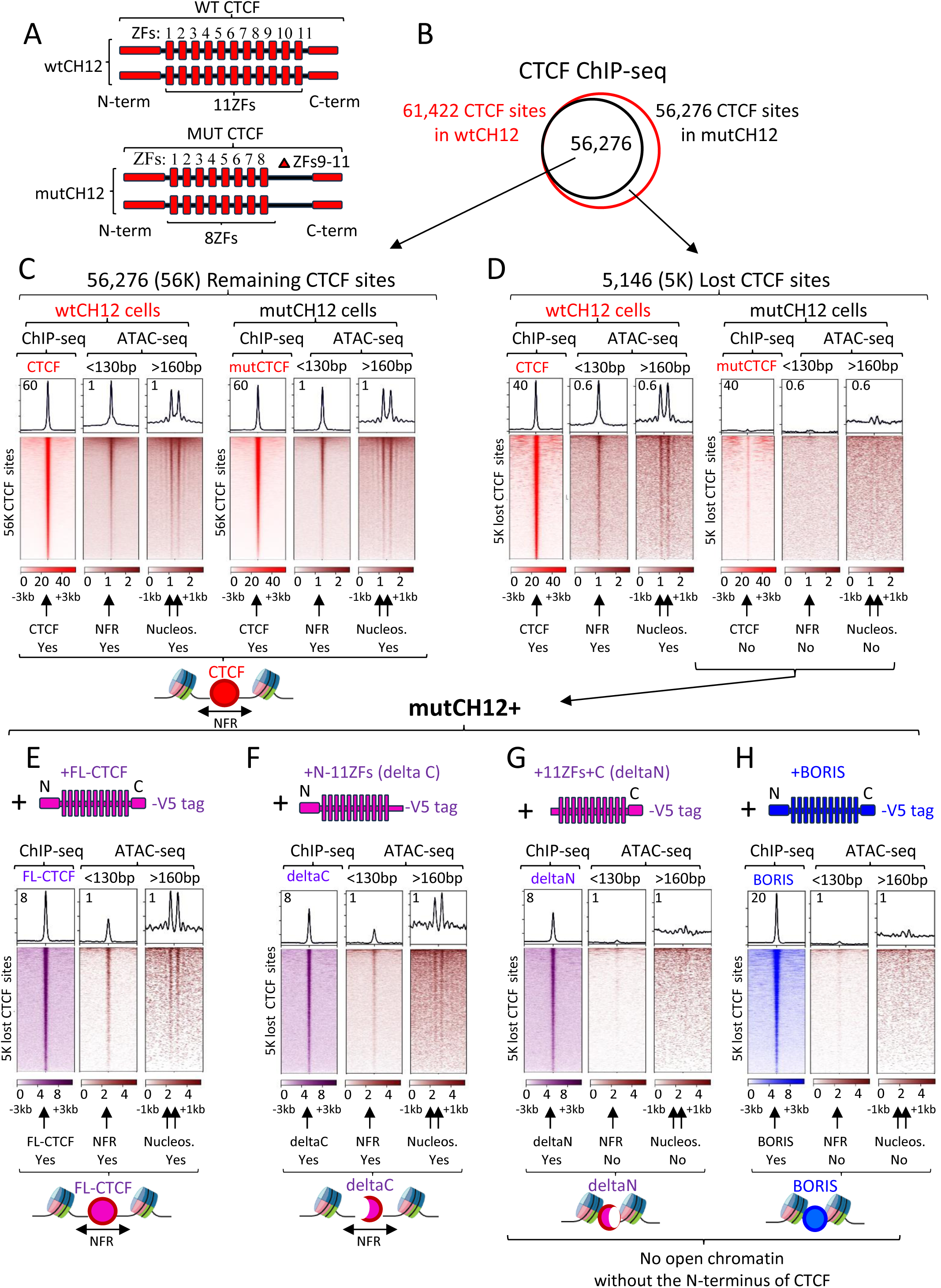
The loss of CTCF occupancy leads to reduced chromatin accessibility, a process dependent on the N-terminus of CTCF. **(A)** Schematic representations of wild-type and mutant CTCF proteins in wtCH12 and mutCH12 cells, respectively. The mutant CTCF has a homozygous deletion of zinc fingers 9-11 (ZFs 9-11). **(B)** Venn diagram illustrating the overlap of CTCF binding sites mapped by ChIP-seq in wtCH12 versus mutCH12 cells. **(C,D)** Heatmaps showing CTCF occupancy in relation to chromatin accessibility (ATAC-seq) in wtCH12 and mutCH12 cells. The analysis focuses on sites where CTCF binding is either remained **(C)** or lost **(D**) in mutCH12 cells. Paired-end ATAC-seq reads were classified into short fragments (<130 bp), representing CTCF footprints, and nucleosome-sized fragments (>160 bp) to assess nucleosome positioning. **(E-H)** Chromatin accessibility analysis in mutCH12 cells ectopically expressing different CTCF constructs: Full-length CTCF (FL-CTCF**) (E)**, CTCF truncated at the C-terminus (deltaC**) (F),** CTCF truncated at the N-terminus (deltaN) **(G)**, human BORIS **(H)**. At the top of each panel, a schematic representation of the ectopically expressed protein is displayed, mapped by ChIP-seq, with occupancy at the 5K lost CTCF sites visualized in heatmaps labeled “ChIP-seq.“ At the bottom of each heatmap **(C-H):** The presence of Nucleosome-Free Regions (NFR) and the two flanking nucleosomes (Nucleos.) around the summit of CTCF ChIP-seq data is indicated with black arrows. Labels “Yes” or “No” denote the presence or absence of these features. Below each panel is a schematic illustrating the proposed binding of the ectopically expressed protein in relation to nucleosome positioning and open chromatin states. The heatmaps **(C-H)** are centered at the summit of CTCF ChIP-seq peaks, spanning a 6 kb window for ChIP-seq data and a 2 kb window for ATAC-seq data.

### Identification of CTCF binding motifs/consensuses

For CTCF motif analysis, we used the summit CTCF ChIP-seq or CTCF Affinity-Seq BED files generated by MACS2. The sequences around the summits of ChIP-seq or Affinity-Seq peaks were extended by 100 bp upstream and downstream for CTCF motif discovery. FASTA files were generated using BEDTools (getfasta option) based on either the human (hg19) or mouse (mm9) genome. These FASTA files were then used to recover position weight matrices using Multiple EM for Motif Elicitation (MEME) software (67). We ran MEME with the parameters (-nmotifs 1 -mod oops -revcomp -w 15, or -w20, or -w40) to identify the motif for CTCF ChIP-seq or CTCF Affinity-seq peaks. To determine the orientation of CTCF motifs for each CTCF ChIP-seq peak, we used the FIMO tool from the MEME suite with default settings (p-value < 0.0001). FIMO allowed us to identify the genomic coordinates and strand-specific CTCF motif orientation under each ChIP-seq peak called by MACS2. Based on the strand-specific orientation, CTCF motifs were classified as either plus or minus motifs for further analysis. To generate heatmaps in deepTools that account for strand-specific information, we included strand data directly in the BED file with genomic coordinates (6^th^ column). This enables deepTools’ computeMatrix program to handle strand orientation correctly, aligning data in the direction of CTCF binding site orientation. In cases where the 100 bp window around the summit of a CTCF peak contained more than one CTCF motif, we selected the motif with the most significant p-value. If several motifs within a single CTCF ChIP-seq peak had identical p-values, those sites were excluded from the analysis. The list of plus and minus CTCF motifs used in this study is provided in Table S1. Notably, not all CTCF ChIP-seq peaks were associated with a MEME PWM (Position Weight Matrix) p-value < 0.0001, as some contained variations of the CTCF motif or motifs with lower statistical significance. These CTCF ChIP-seq peaks were excluded from the deepTools heatmap analysis, along with peaks containing multiple CTCF consensus sequences. Consequently, the total number of mapped CTCF ChIP-seq peaks exceeds the number of CTCF motifs presented in Table S1, which were used for heatmap generation. To calculate the total number of CTCF motifs in the mouse genome, we used the position weight matrices from CTCF ChIP-seq data in CH12 cells to scan the mouse genome (mm9) using the FIMO tool with default settings (p-value < 0.0001). To identify motif enrichment around CTCF ChIP-seq peaks, we used the MEME AME (Analysis of Motif Enrichment) tool, analyzing 400-bp sequences centered on the CTCF summits against the JASPAR CORE and UniPROBE Mouse motif databases.

### AlphaFold-Multimer prediction

We used the AlphaFold Server (https://alphafoldserver.com/) to predict potential interactions between CTCF, BORIS, and SMARCA5. Interactive Predicted Aligned Error (PAE) viewer plots were generated by the server using the amino acid sequences for CTCF (Sequence ID: NP_006556.1), BORIS (Sequence ID: NP_001255969.1), and SMARCA5 (Sequence ID: NP_003592.3). The accuracy of the predicted multimeric structures was evaluated using AlphaFold-Multimer (68), which employs Interface Predicted Template Modelling (ipTM) to assess prediction quality.

## Results

### About sevenfold discrepancy between in vivo bound and potentially available CTCF binding sites

We initiated this study by asking how does the status of chromatin influence cell-specific CTCF occupancy. One of the most straightforward ways to address this question is to compare CTCF binding to chromatin with CTCF binding to naked DNA isolated from the same type of cells. This comparison provides information on how many sites are available for CTCF binding on DNA and how many are indeed bound in chromatin.

To this end, we used previously published studies (51,52) to compare CTCF binding sites in mouse spleen identified by either ChIP-seq (chromatin) or Affinity-seq (naked genomic DNA), using either CTCF antibodies or the 11 Zinc Finger DNA-binding domain of CTCF, respectively (Figure 1A). Using these two genome-wide approaches, we mapped 26,208 CTCF binding sites in chromatin and 191,231 potential CTCF binding sites in naked genomic DNA of mouse spleen, respectively. (Figure 1A-B). The majority of CTCF sites (96%) bound *in vivo* (chromatin) were also identified by the 11 ZFs binding to naked genomic DNA (*in vitro*), confirming that the CTCF binding domain is a main factor of CTCF occupancy (Figure 1B).

However, 167,132 potential CTCF binding sites were not occupied by CTCF in vivo, despite being accessible to the 11-zinc-finger (ZF) domain binding as identified by Affinity-seq. Since Affinity-seq preserves genomic CpG methylation, which is known to negatively affect CTCF binding (21), DNA methylation alone cannot fully explain the absence of CTCF occupancy at these sites. Given that these sites were free of CpG methylation and available for 11 ZF binding, we hypothesized that other regulatory factors may influence CTCF occupancy *in vivo*. Notably, a comparison of ChIP-seq and Affinity-seq data revealed that CTCF binding affinity is more uniform when the 11-ZF domains bind directly to genomic DNA, as observed in Affinity-seq. This contrasts with the variable binding strength observed in ChIP-seq, which is modulated by the chromatin environment (Supplementary Figure S1A).

Motif analysis of CTCF binding sites detected by ChIP-seq and Affinity-seq identified the classic 14-bp CTCF core consensus motif at the summit of the recovered peaks (Figure 1B). However, this motif alone is not sufficient to predict CTCF occupancy. When the mouse genome (mm9) was scanned for the CTCF consensus, over 900,000 potential CTCF binding sites were detected, most of which were never bound by CTCF in the spleen or any other mouse cell types (Figure 1C and Supplementary Figure S1B). Interestingly, there were 1,772 CTCF ChIP-seq peaks in the mouse spleen that were not recovered by Affinity-seq. These CTCF peaks exhibited a degenerate variant of CTCF consensus motif (Figure 1B), leading to significantly reduced CTCF binding affinity compared to peaks with a canonical CTCF motif (Supplementary Figure S1A). Given that CTCF occupancy can be modulated by other transcription factors (17), a motif analysis conducted within a 400 bp window around the summit of the 1,772 CTCF ChIP-seq peaks revealed significant enrichment of binding sites for various B-cell specific transcription factors, including members of the IKAROS family (Supplementary Figure S1C). These transcription factors may contribute to stabilizing weak CTCF binding events at these regions. Thus, this analysis indicates that neither the presence of the CTCF consensus motif nor the absence of CpG methylation is fully sufficient to explain the cell-specific occupancy of CTCF *in vivo*.

### CTCF motifs are strategically positioned at the entry sides of well-positioned nucleosomes, such that upon binding, the N-terminus of CTCF is oriented towards the nucleosome

Based on the comparison of ChIP-seq and Affinity-seq data (Figure 1B), it appears there are potentially seven times more available CTCF binding sites than those actually bound by CTCF in mouse spleen chromatin. Importantly, CTCF ChIP-seq data from various mouse tissues (ENCODE data) show that these potential binding sites are occupied by CTCF in other cell types (Figure 1C and Supplementary Figure S1B,D). This suggests that these sites are valid CTCF binding sites, likely free from CpG methylation (due to their accessibility to the 11 ZFs of CTCF), but exhibit tissue-specific chromatin properties that restrict CTCF binding in the spleen. We aim to investigate which DNA and chromatin features that render these potential CTCF sites unavailable for binding in the spleen while allowing CTCF binding in other cell types.

In Figure 1C, a genome browser visualization demonstrates that the potential CTCF binding sites recovered by Affinity-seq, but not by ChIP-seq in mouse spleen, are bound by CTCF in mouse B-cell lymphoma (CH12) and mouse embryonic stem (mES) cells (shown by arrows). There are also Affinity-seq recovered CTCF sites that are not bound by CTCF in CH12 and mES cells but bound by CTCF in other cell lines (Supplementary Figure S1B, D). Based on these observations, for future analysis, we categorized CTCF binding sites into three groups: (Group #1) sites bound by CTCF in the chromatin of the analyzed cells (CTCF-bound, ChIP-seq), (Group #2) Affinity-seq recovered sites not bound by CTCF in the chromatin of the analyzed cells (11 ZFs-only, naked DNA), and (Group #3) CTCF consensus sequences that are not bound by CTCF *in vivo* and not recovered by Affinity-seq (CTCF motif, not bound by CTCF) (Figure 1C). For each group, we randomly selected approximately 32,000 CTCF binding motifs and analyzed them in the plus (sense) orientations, reversing the signal from the minus orientation to match the plus strand (Figure 1D and Supplementary Figure S2).

Previous studies have shown that CTCF competes with nucleosomes for DNA binding *in vivo* (36,40,41). Once bound to chromatin, CTCF can reposition nucleosomes on either side of its binding site (33). To investigate nucleosome positioning at the three selected groups of CTCF binding sites, we analyzed MNase-assisted H3 ChIP-seq data for both CH12 and mES cells (Figure 1D and Supplementary Figure S2). Our analysis first focused on plus-oriented CTCF motifs bound by CTCF in CH12 cells (Group #1) (Figure 1D, E). The combined analysis of CTCF occupancy via ChIP-seq and Affinity-seq, along with nucleosome positioning, revealed distinct patterns. High-occupancy CTCF ChIP-seq peaks were primarily found in nucleosome-free regions flanked by well-positioned nucleosomes, while low-occupancy CTCF sites exhibited less stable binding and showed a nucleosome approaching from the right side of plus-oriented CTCF sites (shown by blue arrow in Figure 1D). Clustering of CTCF-bound sites (Group #1) into three clusters based on occupancy levels revealed that low-occupancy CTCF sites, unlike high-occupancy sites, exhibit a narrower DNA footprint influenced by the nucleosome positioned on the right of the plus CTCF motif (Supplementary Figure S3A, blue arrow). Interestingly, the same nucleosome on the right is well-positioned at Affinity-seq derived CTCF binding sites that are not bound by CTCF in CH12 cells (Group #2) (Figure 1D, E, blue arrow). This nucleosome is positioned such that, once CTCF binds, its N-terminus is oriented towards the nucleosome (32), with the CTCF consensus sequence located at the entry side of the nucleosome. We refer to this nucleosome as the CTCF Priming Nucleosome (CPN). The same analysis of CTCF sites bound or not bound by CTCF in mES cells showed consistent results with those observed in CH12 cells (Supplementary Figure S2). Additionally, re-analysis of CTCF depletion in mES cells using the auxin-degron system (57) revealed that upon CTCF loss, nucleosomes, including the CPN, shifted inward toward the CTCF site but did not fully occupy it (Supplementary Figure S3C,D), suggesting that the positioning of the CPN and CTCF occupancy are interrelated across cell types.

Based on these observations, we propose that CTCF binding sites are strategically positioned at the entry side of the CPN. At sites where CTCF strongly binds, it displaces the CPN for stable DNA binding, with its N-terminus oriented toward the nucleosome, shifting the nucleosome downstream from the binding site (Figure 1E, schematic, black arrow). In contrast, at sites identified only by Affinity-seq, CTCF is unable to displace the CPN for a strong DNA binding (Figure 1D, E and Supplementary Figures S2, S3A). Therefore, we suggest that specific epigenetic features of the CPN may regulate CTCF’s ability to bind in chromatin.

### CTCF cannot successfully displace the priming nucleosome if the nucleosome carries the repressive histone mark H3K9me3 or is wrapped in a CpG-methylated sequence

Our next objective was to identify the epigenetic features that influence CTCF’s ability to displace nucleosomes at CTCF-bound sites (identified by ChIP-seq) compared to sites identified by Affinity-seq and not bound by CTCF in vivo. To achieve this, we compared CTCF ChIP-seq data from CH12 cells with Affinity-seq data, classifying CTCF binding sites into two categories: sites bound by CTCF in CH12 cells (58,010 sites) and sites recovered by Affinity-seq but not bound by CTCF in CH12 cells (133,108 sites) (Supplementary Figure S4A). Next, we overlapped these two groups of CTCF sites with histone modification and variant data mapped by ENCODE in CH12 cells. This analysis revealed that the histone variant H2A.Z is significantly enriched at CTCF-bound sites, whereas the repressive histone modification H3K9me3 is predominantly enriched at sites not bound by CTCF (Supplementary Figure S4B).

To further investigate, we analyzed ChIP-seq data for H2A.Z and H3K9me3 at three groups of CTCF sites classified in Figure 1D. High-occupancy CTCF-bound sites (Group #1) were flanked by nucleosomes carrying the active histone variant H2A.Z and were completely depleted of the repressive histone mark H3K9me3. In contrast, not bound by CTCF sites (11ZFs-only, Group #2) and low-occupancy CTCF sites within the CTCF-bound group (Group #1) were enriched with H3K9me3 at the CPN (Figure 1D). These findings suggest that the CPN, marked by H3K9me3, may inhibit CTCF’s ability to displace nucleosomes for strong DNA binding. Conversely, H2A.Z may facilitate nucleosome displacement, consistent with previous studies implicating SMARCA5 in this process (69). Clustering of CTCF-bound sites (Group #1) and analysis of mES cell data further confirmed that H3K9me3 enrichment at the CPN restricts CTCF binding within the chromatin environment (Supplementary Figures S2, S3A).

Since H3K9me3 histone modification is often associated with CpG methylation (70), we extended our analysis to include the analysis of CpG methylation in CH12 and mES cells at the three groups of selected CTCF sites (Figure 1D, Supplementary Figures S2, S3A). This analysis showed that CTCF-bound sites (Group #1) were associated not only with the absence of CpG methylation at the CTCF consensus itself but also at the DNA sequence wrapped around the CPN (Figure 1D and Supplementary Figures S2, S3A). Thus, the methylation status of DNA wrapped around the CPN is also important for CTCF occupancy. In contrast, CTCF sites recovered by Affinity-seq only (Group #2) but not bound *in vivo* showed enrichment of CpG methylation precisely at the CPN, thus preventing a strong CTCF occupancy *in vivo* (Figure 1D). Group #3, consisting of CTCF sites never bound by CTCF, showed more widespread enrichment of both H3K9me3 and CpG methylation outside of the CPN (Figure 1D). Similar patterns were observed in mES cells, leading to the same conclusions (Supplementary Figure S2).

Additionally, the analysis of CTCF binding sites specific to either CH12 or mES cells revealed that the presence of H3K9me3 and CpG methylation at the CPN inhibits or reduces CTCF binding in a cell-specific manner (Supplementary Figure S4C). For example, CTCF sites that are bound only in mES cells are enriched with both H3K9me3 and CpG methylation in CH12 cells, suggesting a repressive environment that blocks CTCF binding. Conversely, CTCF sites bound only in CH12 cells are depleted of these repressive marks, allowing for strong CTCF binding in these regions (Supplementary Figure S4C). Similar patterns were observed in the human cell lines K562 and MCF7, further supporting the idea that the cell-specific occupancy of CTCF is influenced by the epigenetic landscape surrounding the CPN (Supplementary Figure S5A-C).

Based on these observations, we propose that cell-specific CTCF occupancy is determined by CTCF’s ability to reposition the CpN for effective DNA binding. When the CpN is enriched with CpG methylation and/or H3K9me3 histone modifications, CTCF is less capable of repositioning the nucleosome, resulting in reduced or absent binding. Conversely, when the CPN incorporates an active histone variant such as H2A.Z, CTCF can effectively displace the nucleosome, facilitating stronger binding. Thus, the methylation status and histone modification state of the CpN are critical factors in determining CTCF occupancy.

### The loss of H3K9me3 at the CTCF priming nucleosome opens CTCF sites for a stable CTCF occupancy

As we describe the interrelation between high CTCF occupancy and the absence of the H3K9me3 mark on the CPN, we tested this hypothesis by examining whether CTCF sites identified by Affinity-seq in spleen cells but not bound by CTCF in vivo could be attributed to the enrichment of H3K9me3 at the CPN. Analysis of H3K9me3 ChIP-seq data in mouse spleen confirmed that the CPN of CTCF sites not occupied in vivo showed clear enrichment of repressive histone marks, whereas these marks were absent from the CPN of CTCF-bound sites (Supplementary Figure S5D). This finding corroborates observations in CH12 and mES cells (Figure 1D and Supplementary Figures S2, S3A), reinforcing that the presence of H3K9me3 at the CPN is a key factor preventing strong CTCF binding in vivo.

To further support this, we reanalyzed published data reporting that the knockout (KO) of Setdb1, a histone methyltransferase, in mES cells increased CTCF occupancy as H3K9me3 levels decreased (28). Comparing CTCF ChIP-seq data from wild-type and Setdb1 knockout (KO) cells, we identified at least 3,608 CTCF binding sites where CTCF occupancy increased at least twofold (p-value < 0.01) upon Setdb1 KO (Figure 2A). The combination of CTCF ChIP-seq, MNase-seq, and H3K9me3 ChIP-seq at these 3,608 sites showed that increased CTCF occupancy was accompanied by a downstream shift of the CPN, moving away from the N-terminus of CTCF as H3K9me3 was lost specifically at the priming nucleosome (Figure 2B,C). In parental mES cells, the plus-oriented consensus for the 3,608 CTCF sites was located at the entry side of the CPN marked by H3K9me3, and this repressive modification inhibited CTCF binding. Upon loss of H3K9me3, CTCF occupancy dramatically increased. Although the H3K9me3 heatmap data at these sites appeared to position the repressive histone modification directly at the CTCF consensus (Figure 2B), more precise analysis revealed that the average summit of H3K9me3 ChIP-seq peaks, which were lost upon Setdb1 KO, is located at least 70 bp downstream from the start of the 14-bp CTCF core consensus, downstream from the N-terminus of CTCF (Figure 2D). These results confirm our hypothesis that the repressive epigenetic status of the CPN hinders strong CTCF binding. However, upon removal of H3K9me3, the CPN shifts downstream, allowing strong CTCF binding (Figure 2E).

### High- and low-occupancy CTCF sites are associated with specific grammars of the CTCF core consensus sequence

Upon examining the genome browser views of CTCF ChIP-seq data across multiple cell lines (Figure 1C, Supplementary Figure S1D) and the heatmaps generated in this study, we observed that low-occupancy, small CTCF ChIP-seq peaks, those not classified as significant peaks by the MACS algorithm, were recovered by Affinity-seq as strong CTCF binding sites. Furthermore, these peaks are also identified as cell-specific CTCF binding sites in other cell types, where they display high CTCF occupancy. This low CTCF occupancy is indicated by yellow arrows in Figure 1C,D and Supplementary Figures S1A, S1D, S2, S3A, S4C, S5C, S5D, S6. Based on these observations, we suggest that there are many more CTCF binding sites in vertebrate cells than previously recognized. These sites may not be classified as significant ChIP-seq peaks but still bind CTCF briefly and specifically, provided that the CTCF consensus is not methylated. The repressive status of the CpN appears to inhibit stronger CTCF binding at these sites. The functionality of these sites remains an open question, but they do show a low enrichment of cohesin, suggesting their potential and brief involvement in 3D genome organization (Supplementary Figure S1D).

To analyze these low-occupancy CTCF sites in greater detail (the group#2 from Figure 1D), we clustered them into three groups based on the enrichment of CTCF ChIP-seq reads (Supplementary Figure S6A). Regarding nucleosome positioning and histone modifications, these low-occupancy CTCF sites did not exhibit significant differences in the enrichment of H2A.Z, H3K9me3, and CpG methylation across the three different clusters (Supplementary Figure S6A), unlike the distinct patterns observed for the clustering of CTCF-bound sites (Supplementary Figure S3A). Motif analysis of the different clusters of CTCF-bound sites versus Affinity-seq-recovered CTCF non-bound sites revealed differences in the grammar of the 14-bp core CTCF consensus sequence. High-occupancy CTCF-bound sites preferentially featured C, G, and G nucleotides at positions 4, 8, and 13, respectively, within the 14-bp CTCF consensus sequence (Supplementary Figure S3A). In contrast, low-occupancy, cell-specific, and Affinity-seq only CTCF binding sites predominantly contained G, A, and A nucleotides at these critical positions (Supplementary Figures S1D, S3A-B, S4C, S6A). These findings suggest that the intrinsic DNA-binding affinity of the CTCF zinc finger domains is one of the key factors influencing its occupancy in vivo.

Since CTCF binding in chromatin is linked to the chromatin environment, we further clustered 191,231 CTCF binding sites, mapped by Affinity-seq, into seven clusters based on their binding strength (Affinity-seq read enrichment) (Supplementary Figure S6B). This clustering, reflecting interactions of the 11ZFs of CTCF with naked DNA, reinforced the initial observation regarding the grammar of the CTCF motif. The strongest CTCF binding, independent of chromatin context, was consistently associated with C, G, and G nucleotides at positions 4, 8, and 13, respectively (Supplementary Figure S6B). Sites with weaker CTCF binding progressively exhibited more frequent G, A, and A nucleotides at these positions (Supplementary Figure S6B). The read enrichment of CTCF measured by Affinity-seq was also reflected in CTCF ChIP-seq data across different cell lines (Supplementary Figure S6B), further supporting the conclusion that the grammar of the CTCF consensus sequence significantly influences CTCF binding both in vitro and in vivo.

### Cell-specific CTCF occupancy is pre-marked and stable within each cell type

Our findings outlined above suggest that cell-specific CTCF occupancy is influenced by the epigenetic status of the CPN. It is known that each cell type exhibits a unique pattern of CTCF occupancy, which plays a crucial role in 3D genome organization and the maintenance of cell identity (71). During mitosis, the majority of CTCF is released from chromatin, necessitating the re-establishment of the same CTCF binding pattern in daughter cells post-mitosis as was present in the mother cell (46). The use of auxin-degron systems to analyze CTCF dynamics has shown that, upon withdrawal of auxin, CTCF restores its binding pattern to its pre-depletion state (47). These observations suggest that cell-specific CTCF binding patterns are preprogrammed and preserved within the chromatin state of each cell type.

In this study, we sought to explore the mechanisms by which cell-specific CTCF occupancy is maintained and re-established, focusing particularly on the potential role of the CPN in pre-marking future CTCF binding sites. We hypothesized that the status of the CPN may serve as a memory marker for CTCF binding, preserving the specific occupancy pattern seen in each cell type across cell divisions. To test this hypothesis, we utilized a model system involving wild-type (wt) CH12 cells and mutant (mut) CH12 cells. The mutCH12 cells were previously generated through the homozygous deletion of CTCF zinc fingers (ZFs) 9-11, resulting in a truncated CTCF protein (Figure 3A, schematic) (49). This modification leads to the loss of mutant CTCF binding at approximately 5,000 (5K) CTCF sites in mutCH12 cells, as observed in comparative analyses of CTCF ChIP-seq data with wtCH12 cells (Figure 3B) and in (48). Previously, we demonstrated that ectopic expression of full-length (FL) CTCF in mutCH12 cells restores CTCF occupancy at these 5K lost sites, replicating the pattern observed in wtCH12 cells, except for 199 CTCF sites that were entirely lost (48). This suggests that these 5K sites hold inherent chromatin features that mark them for future CTCF occupancy, regardless of the loss of CTCF binding.

To investigate the chromatin features that mark these sites for CTCF binding, we also utilized various truncated forms of CTCF, lacking either the N-terminal (ΔN) or C-terminal (ΔC) regions, while retaining all 11 ZFs necessary for DNA binding to the 5K sites (Supplementary Figure S7A-C). This approach allowed us to dissect the roles of the C- and N-terminal domains of CTCF in the re-establishment of occupancy at the 5K sites. Furthermore, we explored the role of BORIS (CTCFL), a paralog of CTCF that shares a similar 11 ZF DNA-binding domain but differs in the C- and N-terminal regions (72,73). We previously showed that BORIS can bind to the 5K lost CTCF sites in mutCH12 cells, but not in wtCH12 cells, where it would normally compete with CTCF for binding (48).

To comprehensively analyze the chromatin and DNA features associated with the 5K lost CTCF sites, we performed the following assays in wtCH12 and mutCH12 cells: ATAC-seq (to assess chromatin accessibility), MNase-assisted H3 ChIP-seq (to profile nucleosome occupancy), MeDIP-seq (to map DNA methylation), H3K9me3 and H2A.Z ChIP-seq (to identify repressive and active histone modifications). The use of wtCH12 and mutCH12 cells provides a unique system to investigate how the chromatin and DNA status changed with the loss of CTCF binding and the subsequent re-occupancy by CTCF and BORIS proteins (48).

### Loss of CTCF occupancy results in the loss of chromatin accessibility, a state that cannot be restored at the CTCF binding sites without the N-terminus of CTCF

To investigate the changes in chromatin status following the loss of CTCF occupancy at the 5K CTCF sites in mutCH12 cells, we performed ATAC-seq analysis to compare chromatin accessibility in wtCH12 and mutCH12 cells. ATAC-seq using paired-end sequencing allowed us to categorize the reads into short fragments (<130bp), indicative of CTCF footprint, and nucleosome-sized fragments (>160bp) to determine nucleosome positioning (Figure 3C-H). We analyzed chromatin accessibility at two groups of CTCF binding sites: 1). Remaining CTCF Sites (56K): these sites are bound by CTCF in both wtCH12 and mutCH12 cells and serve as a control; 2). Lost 5K CTCF Sites: these sites are bound by CTCF in wtCH12 cells but lose CTCF occupancy in mutCH12 cells (Figure 3C-D).

It is well-documented that CTCF bound sites are located in open chromatin regions with well-positioned nucleosomes flanking these sites (33). Our ATAC-seq analysis confirmed this at the remaining CTCF sites in both wtCH12 and mutCH12 cells, as well as at the 5K lost CTCF sites in wtCH12 cells, showing clear footprints of open chromatin and nucleosome positioning at the flank of CTCF sites (Figure 3C-D and Supplementary Figure S7D). However, in mutCH12 cells, the loss of CTCF occupancy at the 5K sites resulted in a complete loss of both open chromatin and nucleosome positioning around these sites (Figure 3D, right panel). This indicates that CTCF is crucial for maintaining an open chromatin state at its binding sites. Ectopic expression of FL-CTCF (tagged with V5-tag) in mutCH12 cells restored not only CTCF occupancy at the vast majority of the 5K lost sites (as mapped using V5-tag antibodies) but also reestablished open chromatin and nucleosome positioning around CTCF sites (Figure 3E). This restoration suggests that CTCF functions as a pioneer factor capable of binding and opening closed chromatin.

To determine which domain of CTCF is essential for chromatin opening and nucleosome repositioning, we expressed truncated versions of CTCF, tagged with V5, in mutCH12 cells. We found that truncations of either the C-terminus (deltaC) or the N-terminus (deltaN) led to less stable CTCF binding compared to FL-CTCF (Figure 3E-F and Supplementary Figure S7C). Importantly, CTCF lacking the C-terminus was still capable of opening chromatin and repositioning nucleosomes, but CTCF lacking the N-terminus was not (Figure 3F,G). This demonstrates that the N-terminus of CTCF is essential for its ability to open chromatin and reposition nucleosomes. Unexpectedly, the N-terminus truncated version of CTCF was still able to bind the 5K lost CTCF sites without opening of chromatin (Figure 3G). Furthermore, we investigated the role of BORIS, a paralog of CTCF with a different N- and C-termini. Our analysis revealed that while BORIS could not open the chromatin at the 5K lost CTCF sites, it could still bind closed chromatin (as shown by BORIS ChIP-seq peaks) at the 5K sites in mut CH12 cells (Figure 3H). These data are corroborating previous findings regarding the N-terminus of CTCF and BORIS (74).

In conclusion, this analysis suggests that CTCF acts as a pioneer transcription factor capable of opening closed chromatin and repositioning nucleosomes. This pioneer activity is dependent on the N-terminus of CTCF, highlighting its critical role in chromatin dynamics and gene regulation.

### Nucleosome positioning around CTCF binding sites depends on the N-terminus of CTCF

Our ATAC-seq data analysis demonstrated that CTCF-bound sites are associated with open chromatin and flanked by well-positioned nucleosomes, with these chromatin states being influenced by the N-terminus of CTCF (Figure 3). To validate the ATAC-seq findings, we performed micrococcal nuclease digestion followed by H3 histone ChIP sequencing (MNase-H3-ChIP-seq) to assess nucleosome status in wtCH12 versus mutCH12 cells. The MNase-H3-ChIP-seq analysis revealed that several well-positioned nucleosomes flank the 56K remaining CTCF sites in both wtCH12 and mutCH12 cells (Supplementary Figure S7E). However, at the 5K lost CTCF sites in mutCH12 cells, the nucleosome positioning observed in wtCH12 cells was disrupted (Figure 4A-B). In mutCH12 cells, we detected only one well-positioned nucleosome located at the N-terminus of CTCF relative to the 5K lost motif orientation (Figure 4B, blue arrow). The same N-terminal nucleosome was also observed at low occupancy CTCF sites in wtCH12 cells (Figure 4A). By combining MNase-H3-ChIP-seq data at the 5K lost CTCF sites in both wtCH12 and mutCH12 cells, we confirmed that CTCF motifs are located at the entry side of the well-positioned nucleosome in mutCH12 cells, while the same sequences are nucleosome-free when bound by CTCF in wtCH12 cells (Figure 4C). Thus, similar to the Affinity-seq only CTCF sites shown in Figure 1D-E (group #2), in the absence of strong CTCF binding, the N-terminal nucleosome shifts toward the CTCF motif, positioning it at the entrance of what we term the CTCF priming nucleosome (CPN).

**Figure 4.**
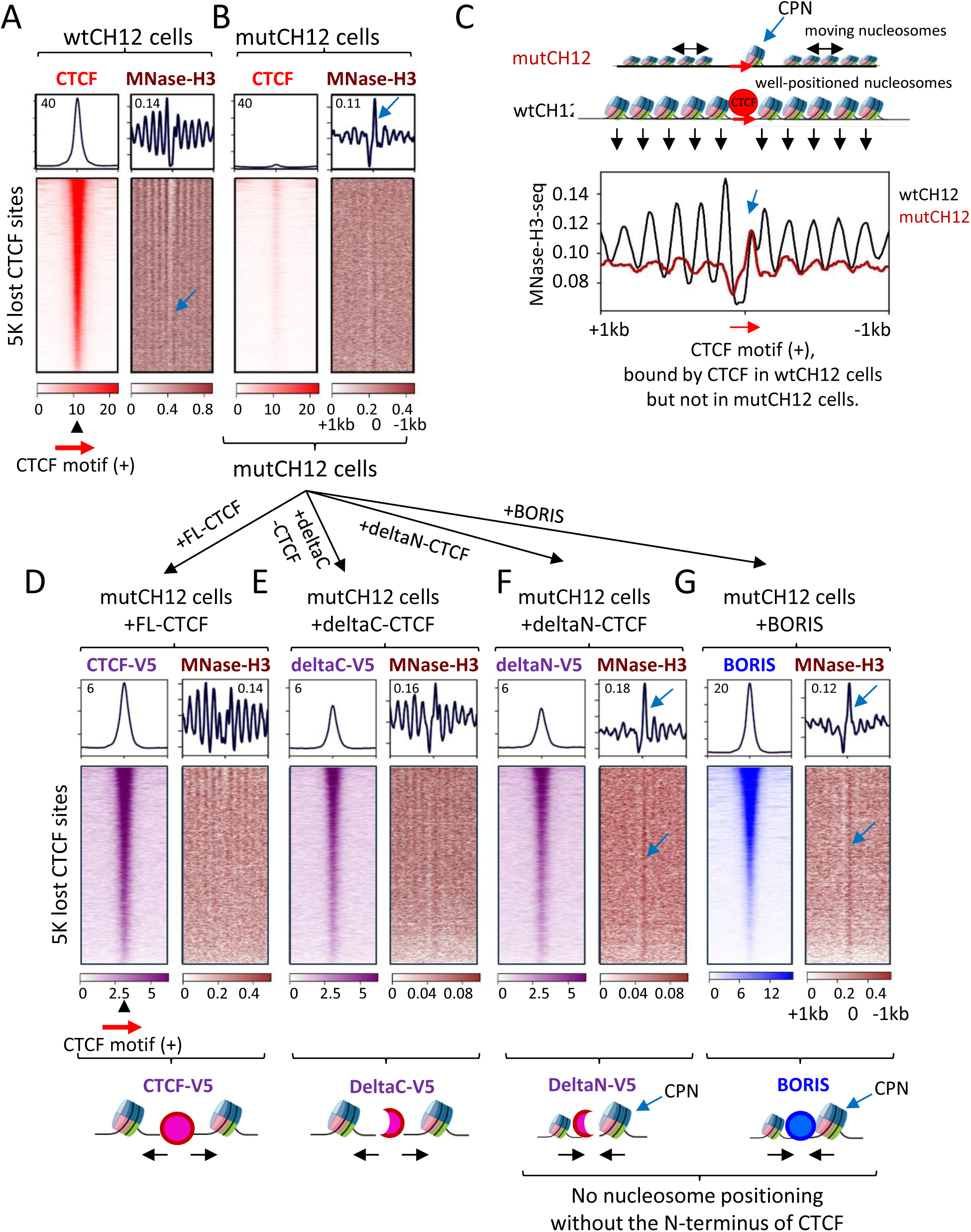
Nucleosome positioning around CTCF binding sites depends on the N-terminus of CTCF. **(A, B)** Heatmaps showing CTCF occupancy (ChIP-seq) combined with nucleosome positioning (MNase-H3), centered on the CTCF motif in the plus (sense) orientation. (**A)** Sites bound by CTCF in wtCH12 cells. (**B)** Sites lost in mutCH12 cells. **(C)** MNase-H3 ChIP-seq profile from panels **(A)** and **(B)** combined into a single plot, comparing nucleosome occupancy at the 5K lost CTCF sites in wtCH12 and mutCH12 cells. A schematic representation above the plot illustrates nucleosome positioning relative to the CTCF motif (red arrow) at the lost sites. **(D–G)** Heatmaps displaying V5-tag density (D–F) or BORIS antibody (Ab) density combined with MNase-H3, following ectopic expression of the indicated CTCF vectors or BORIS in mutCH12 cells. Schematics at the bottom of the heatmaps illustrate the structural outcomes of CTCF truncations, with black arrows indicating the changes in nucleosome positioning and movement. **(A–G)** The CTCF priming nucleosome (CPN) is indicated by blue arrows.

Ectopic expression of full-length CTCF (FL-CTCF) in mutCH12 cells restored the open chromatin state and the positioning of several nucleosomes on each side of the 5K lost CTCF sites (Figures 3E, 4D), further corroborating that both chromatin states depend on CTCF occupancy. When examining nucleosome positioning in mutCH12 cells expressing either a C-terminal truncation (deltaC) or an N-terminal truncation (deltaN) of CTCF, we found that the truncation of the N-terminus prevented the restoration of open chromatin and proper nucleosome positioning around CTCF binding sites (Figures, 3F-G, 4E-F). The deltaC truncation also affected nucleosome positioning but to a lesser extent compared to the N-terminus (Figure 4E). BORIS, like the N-terminus truncation of CTCF, failed to open chromatin or create phased nucleosome arrays (Figures 3H, 4G). Therefore, the MNase-H3-ChIP-seq analysis corroborates our ATAC-seq data, demonstrating that the N-terminus of CTCF is essential for maintaining an open chromatin state and phased nucleosome positioning.

### The N-terminus of CTCF is necessary for the recruitment of the chromatin remodeling factor SMARCA5 (Snf2h) at CTCF binding sites

If we assume that CTCF competes with the CPN for DNA binding, an important question arises: how does CTCF displace the CPN downstream to create open chromatin around its binding site? Previous studies have demonstrated that the chromatin remodeling factor SMARCA5 (Snf2h) is required for CTCF occupancy and for maintaining nucleosome spacing (27,44,45). To investigate how SMARCA5 influences CTCF binding, we analyzed several cell lines in which SMARCA5 depletion was achieved using different strategies. Reanalyzing data from three independent studies (27,44,45), using the same analytical framework as in Figure 1D, we confirmed that SMARCA5 depletion consistently led to a substantial, though incomplete, loss of CTCF occupancy at its binding sites across datasets from both mouse and human cells (Figure 5A-B; Supplementary Figure S8). This loss was accompanied by the appearance of a nucleosome at the CTCF binding site (Figure 5A-C, Supplementary Figure S8) and a reduction in chromatin accessibility (Supplementary Figure S8E).

**Figure 5.**
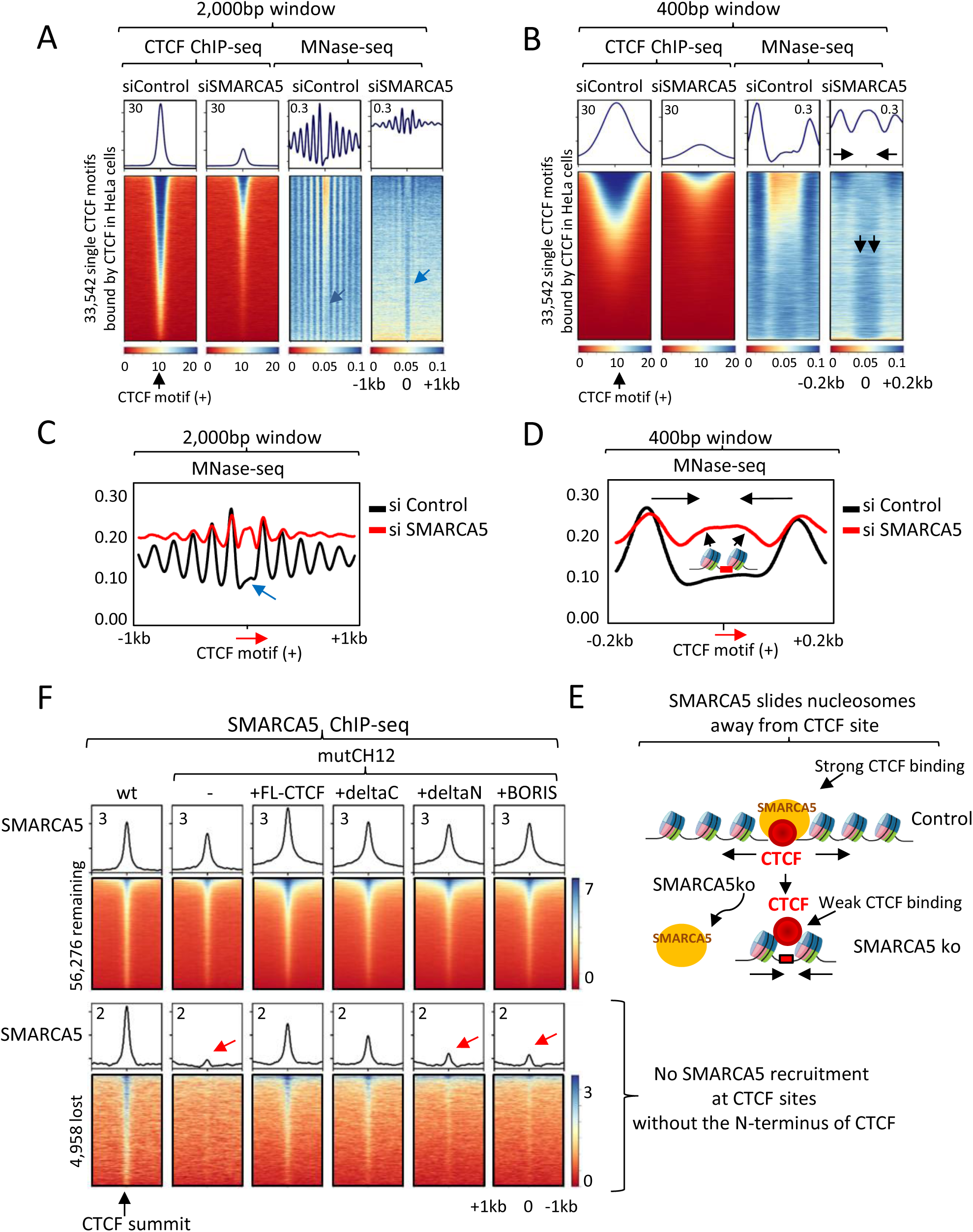
SMARCA5 is recruited by the N-terminus of CTCF to open chromatin around CTCF binding sites. **(A, B)** Heatmaps showing CTCF occupancy (ChIP-seq) and nucleosome positioning (MNase-seq) in HeLa cells with SMARCA5 depletion (siSMARCA5) compared to control HeLa cells (siControl). The heatmaps are centered on CTCF bound motifs inside ChIP-seq peaks in HeLa cells within a window of 2,000 bp **(A)** or 400 bp **(B).** The CTCF priming nucleosome (CPN) is indicated by blue arrows, and the two flanking CTCF nucleosomes shifted toward the CTCF motif are marked by black arrows. **(C, D)** MNase-seq profiles of nucleosome positioning at CTCF-bound motifs in the plus orientation in HeLa cells with (red) and without (black) SMARCA5 depletion. Profiles are shown for windows of 2,000 bp **(C)** and 400 bp **(D). (E)** Schematic representation illustrating the effects of SMARCA5 depletion on CTCF binding and nucleosome occupancy. **(F)** Heatmaps of SMARCA5 ChIP-seq data across two groups of CTCF binding sites: remaining CTCF sites (56,276) bound by CTCF in both wtCH12 and mutCH12 cells (top panel); 4,958 (5K) lost CTCF sites in mutCH12 cells but restored by ectopic expression of FL-CTCF (lower panel). Red arrows indicate the absence of SMARCA5 occupancy at the lost CTCF sites in mutCH12 cells, ectopically expressing either the deltaN CTCF variant or BORIS.

Visualization of nucleosome positioning within a 2,000 bp window revealed that high CTCF occupancy, prior to SMARCA5 depletion, corresponded to a well-ordered array of nucleosomes flanking the CTCF motif (Figure 5A, Supplementary Figure S8A,F). Similar to the observations in Figure 1D, low-occupancy CTCF sites displayed the CPN nucleosome shifted toward the CTCF motif (Figure 5A, blue arrow). However, following SMARCA5 degradation, the ordered nucleosome array was disrupted. Only the two nucleosomes directly flanking the CTCF motif shifted inward, effectively squeezing CTCF out of the binding site and reducing its occupancy (Figure 5A-D). This effect was more pronounced within a 400 bp window, where the inward shift of the flanking nucleosomes correlated precisely with the loss of CTCF occupancy (Figure 5B, D, Supplementary Figure S8C,D).

Since SMARCA5 depletion precedes the reduction in CTCF occupancy, we interpret this as CTCF’s inability to recruit SMARCA5 to displace the CPN nucleosome and establish open chromatin (schematic presentation in Figure 5E). This failure likely results in the observed reduction in CTCF binding. Regarding the CPN nucleosome, CTCF-bound motifs were nucleosome-free prior to SMARCA5 degradation but shifted to the nucleosome entry side post-degradation, weakening CTCF binding (Figure 5A-D; Supplementary Figure S8). These findings align with our data from the 5K lost CTCF sites in mutCH12 cells (Figures 3, 4). A critical distinction between the stable loss of CTCF occupancy in mutCH12 cells and the transient loss resulting from siRNA- or PROTAC-mediated SMARCA5 degradation lies in the behavior of the C-terminal nucleosome. Upon SMARCA5 depletion, this nucleosome also shifts inward toward the CTCF motif. Similar inward repositioning of flanking nucleosomes was observed with auxin-mediated CTCF degradation (Supplementary Figure S3C-D) and nanoNOMe-seq single-molecule analysis of dynamic CTCF binding (75). This difference likely reflects chromatin’s ability to adapt to stable versus transient changes in CTCF binding. Notably, even with SMARCA5 depletion, CTCF retains the ability to recognize and bind its target sites. This conclusion is supported by our data showing that a CTCF variant with a truncated N-terminus can bind the 5K lost sites in mutCH12 cells (Figure 3G), albeit with lower occupancy than full-length CTCF (Supplementary Figure S7C) but fails to reposition the CPN nucleosome for strong binding (Figures 3G, 4F).

SMARCA5 has been shown to directly interact with CTCF by co-IP (76); however, the interacting domains between the two proteins have not been mapped. Using AlphaFold3 predictions, we identified a potential interaction between the N-terminus of CTCF (approximately 155–177 amino acids) and the P-loop-containing nucleoside triphosphate hydrolase domain of SMARCA5 (approximately 160–680 amino acids) (Supplementary Figure S9A). While the confidence of this prediction was low, with an Interface Predicted Template Modeling (ipTM) score of 0.25, it was notably stronger compared to the lack of any predicted interaction between BORIS and SMARCA5 (Supplementary Figure S9B).

To experimentally test the AlphaFold3 prediction and further analyze how SMARCA5 occupancy correlates with CTCF binding, we performed ChIP-seq with SMARCA5 antibodies in both wtCH12 and mutCH12 cells. We found that SMARCA5 occupancy follows CTCF occupancy in all types of CH12 cells at the remaining CTCF binding sites (Figure 5F, top panel). With the loss of CTCF binding at the 5K sites, SMARCA5 occupancy is lost, suggesting that SMARCA5 is recruited by CTCF to chromatin (Figure 5F, lower panel). The ectopic expression of FL-CTCF in mutCH12 cells, restores SMARCA5 occupancy at the 5K lost CTCF sites (Figure 5F, lower panel). However, the truncation of the N-terminus of CTCF or ectopic expression of BORIS in mutCH12 cells failed to restore SMARCA5 occupancy at the 5K CTCF sites (Figure 5F, red arrows). At the same time, the truncation of the C-terminus affected SMARCA5 enrichment, but at lesser extent, probably, because it is affected CTCF binding overall (Figure 5F and Supplementary Figure S7C). As an additional control, we analyzed SMARCA5 occupancy at 199 CTCF binding sites that were irreversibly lost and not restored by ectopic FL-CTCF expression in mutCH12 cells (Supplementary Figure S9C). At these sites, SMARCA5 occupancy was completely absent and not restored, mirroring the pattern observed with the ectopically expressed deltaN CTCF and BORIS. These data suggest that the N-terminus of CTCF is essential for recruiting SMARCA5, which is necessary for opening chromatin and phasing nucleosomes around CTCF binding sites.

### The nucleosome-free accessible chromatin facilitates the halting of cohesin extrusion at CTCF binding sites

The N-terminus of CTCF is crucial for halting cohesin-mediated chromatin extrusion, a process essential for mediating 3D genome organization (48,77–79). Moreover, as demonstrated in this study, the N-terminus of CTCF is also required for nucleosome repositioning and the establishment of open chromatin around CTCF binding sites (Figures 3,4). This observation raises an important question: are these two events, chromatin accessibility and cohesin retention at CTCF sites, interconnected?

Recent studies have shown that chromatin accessibility, or open chromatin, is crucial for cohesin retention at CTCF binding sites (80,81). Thus, the retention of cohesin at CTCF sites depends not only on the direct interaction between the SA2–SCC1 subunits and the N-terminus of CTCF, as proposed in Li et.al., (77), but also on the chromatin state of CTCF bound sites.

To evaluate this hypothesis, we examined CTCF binding sites that are unable to retain cohesin. In each cell type, there is a subset of CTCF sites, referred to as CTCF sites depleted of cohesin (CTCFnotRAD21), which neither exhibit cohesin enrichment nor participate in 3D genome organization (48). Genome browser analysis of these CTCFnotRAD21 sites, combined with ATAC-seq data, revealed that they are generally located in regions inaccessible to Tn5-transposase (Figure 6A). Motif analysis of CTCFnotRAD21sites identified the same core CTCF consensus sequence as found in CTCF sites occupied by cohesin (CTCF/RAD21) (Figure 6B), suggesting that the DNA sequence alone is not sufficient to explain why some CTCF sites are excluded from cohesin occupancy. To extend this analysis on a genome-wide scale, we selected 1,000 CTCF-bound sites with and without cohesin occupancy in wtCH12 cells (Figure 6C,D). These two groups of CTCF sites were analyzed for nucleosome positioning (MNase-H3) and chromatin accessibility (ATAC-seq), confirming the initial observation that low or absent cohesin occupancy correlates with closed chromatin around CTCF sites (Figure 6C,D). Interestingly, SMARCA5 occupancy was equally present at both groups of CTCF sites (Figure 6D). The same analysis, conducted on two additional cell lines, mES and NIH3T3, yielded similar conclusions: CTCF sites lacking cohesin are linked to reduced chromatin accessibility (Supplementary Figure S10).

**Figure 6.**
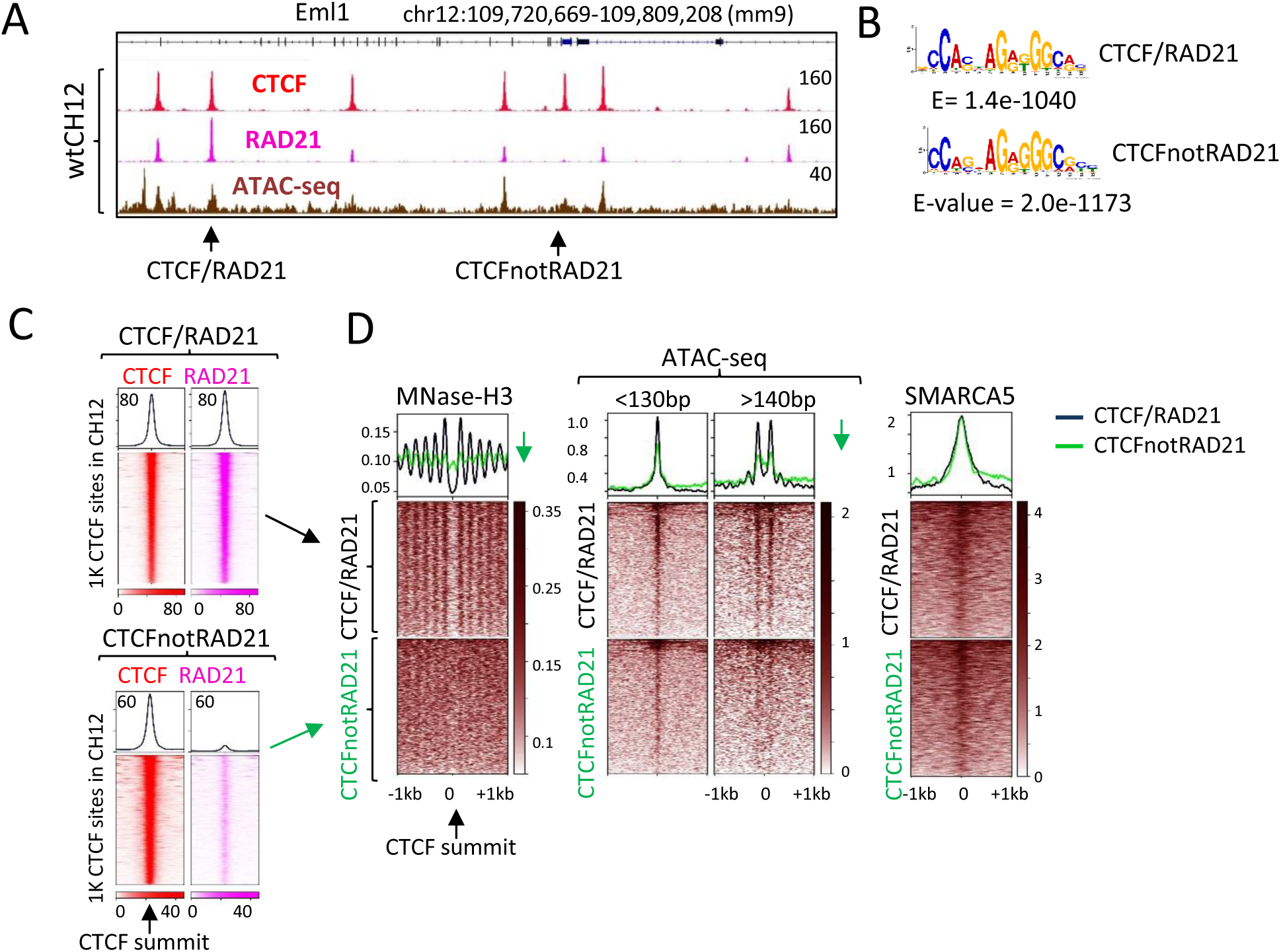
The nucleosome-free accessible chromatin facilitates the halting of cohesin extrusion at CTCF binding sites. **(A)** Genome browser view displaying CTCF (red) and RAD21 (pink) ChIP-seq data alongside ATAC-seq data (brown) in wtCH12 cells. CTCF sites either depleted of cohesin (CTCFnotRAD21) or enriched with cohesin (CTCF/RAD21) are indicated by black arrows. **(B)** CTCF motifs identified within CTCFnotRAD21 and CTCF/RAD21 ChIP-seq peaks in wtCH12 cells. **(C)** Heatmap of CTCF (red) and RAD21 (pink) ChIP-seq occupancy at 1,000 CTCF/RAD21 and CTCFnotRAD21 binding sites in wtCH12 cells, centered at the summit of CTCF peaks. **(D)** Heatmap showing MNase-H3 nucleosome positioning, ATAC-seq (with reads <130 bp indicative of CTCF footprint, and nucleosome-sized fragments >140 bp to determine nucleosome positioning), SMARCA5 ChIP-seq at CTCF/RAD21 (black) and CTCFnotRAD21 (green) binding sites in wtCH12 cells. Panel **(C)** is connected to the data presented in panel **(D)**. Green downward arrows summarize the reduced accessibility (less open chromatin, less positioned array of flanking nucleosomes) at CTCFnotRAD21 sites compared to CTCF/RAD21 sites.

Moreover, in our recent publication, we showed that the ectopic expression of BORIS in NIH3T3 cells increased CTCF occupancy and chromatin accessibility around CTCF sites via the SRCAP-H2A.Z pathway (24). Cohesin occupancy analysis at these sites revealed that some CTCF sites, depleted of cohesin in NIH3T3 cells (before BORIS expression) converted into CTCF cohesin enriched sites with BORIS expression (NIH3T3+BORIS cells). This conversion was accompanied by chromatin opening around CTCF sites (Supplementary Figure S11A). Conversely, some CTCF/RAD21 sites were converted into CTCFnotRAD21 sites in NIH3T3+BORIS cells compared to NIH3T3+EV cells, a process accompanied by a loss of chromatin accessibility at these CTCF sites (Supplementary Figure S11B).

Thus, nucleosome repositioning and chromatin opening at CTCF binding sites appear to be integral to the halting of cohesin extrusion during 3D genome organization.

### CTCF occupancy is largely determined by the permissive epigenetic status around its binding sites

To investigate what preconditions the 5K lost CTCF sites in mutCH12 cells for future CTCF occupancy, we analyzed the CpG methylation status using 5mC MeDIP-seq in both wtCH12 and mutCH12 cells. First, we compared CpG methylation enrichment at CTCF motifs that were either bound or not bound by CTCF in wtCH12 cells (Figure 7A). This analysis revealed a stark difference: CTCF-bound sites were completely depleted of CpG methylation, whereas not bound CTCF sites were enriched with CpG methylation. This finding aligns with the well-established fact that CpG methylation negatively affects CTCF binding (21,22). Next, we examined the CpG methylation status across three distinct groups of CTCF binding sites in CH12 cells: (#1) Remaining CTCF sites: 56,276 sites that are bound by CTCF in both wtCH12 and mutCH12 cells; (#2) 5K lost CTCF sites: 4,958 sites that lost CTCF binding in mutCH12 cells but could be restored with ectopic expression of FL-CTCF; (#3) Irreversibly lost CTCF sites: 199 sites that were irreversibly lost in mutCH12 cells and could not be restored with FL-CTCF expression (Figure 7B). The MeDIP-seq analysis demonstrated that while the remaining CTCF sites (#1) were completely devoid of CpG methylation, the 5K lost CTCF sites (#2) began to acquire some CpG methylation in the absence of CTCF, as indicated by the enrichment of MeDIP-seq reads at the CTCF motifs in mutCH12 cells compared to wtCH12 cells (Figure 7B and Supplementary Figure S12A, shown by the black arrow). In contrast, the irreversibly lost CTCF sites (#3) exhibited a more pronounced gain of CpG methylation (Figure 7B, red arrow). Since CTCF binding is methylation-sensitive, the observed methylation patterns across the three groups (Figure 7B) may explain the restoration of CTCF binding in mutCH12 cells. The persistence of low methylation levels at the 5K lost sites (Group #2) in mutCH12 cells permits potential CTCF re-binding when full-length 11ZFs CTCF is ectopically expressed. Conversely, the gain of CpG methylation at the 199 sites prevents CTCF re-binding, highlighting the critical role of methylation in regulating CTCF binding.

**Figure 7.**
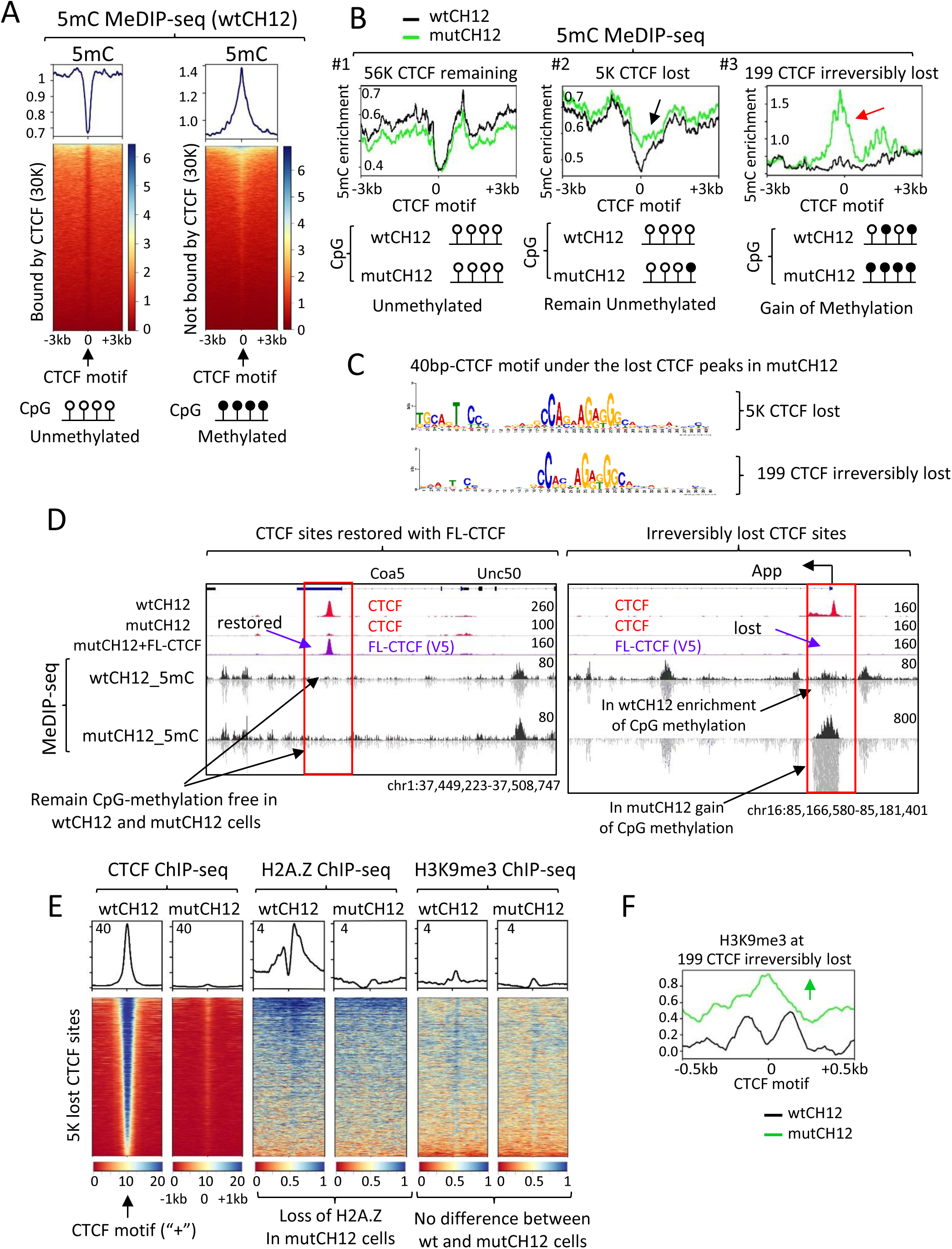
CTCF occupancy is shaped by the permissive epigenetic status around CTCF binding sites. **(A)** Heatmaps showing 5mC MeDIP-seq read alignment centered at CTCF motifs either bound by CTCF (left panel) or not bound by CTCF (right panel) in wtCH12 cells. Below the heatmaps, a schematic representation of CpG methylation status is provided, with open circles indicating unmethylated CpGs and filled black circles representing methylated CpGs, summarizing the data from the corresponding heatmaps. **(B)** Profile of 5mC MeDIP-seq across three groups of CTCF binding sites: (left panel) remaining CTCF sites (56,276) bound by CTCF in both wtCH12 and mutCH12 cells; (middle panel) 4,958 lost CTCF sites (5K) in mutCH12 cells, which are restored by ectopic expression of FL-CTCF; and (right panel) 199 irreversibly lost CTCF sites. Black and red arrows indicate the gain of CpG methylation at the 5K lost CTCF sites and at the irreversibly lost CTCF sites, respectively. A schematic representation of CpG methylation status at the three groups of CTCF sites is shown below the 5mC profiles, highlighting that the irreversibly lost CTCF sites gained CpG methylation (filled black circles) in mutCH12 cells compared to wtCH12 cells. **(C)** The 40-bp MEME-recovered CTCF consensus sequence under the 5K lost CTCF sites compared to the 199 irreversibly lost CTCF sites. **(D)** Genome browser view of CTCF (red) and FL-CTCF-V5 (purple) ChIP-seq data combined with 5mC MeDIP-seq data in wtCH12 cells and mutCH12 cells. The restoration status of CTCF occupancy with ectopic expression of FL-CTCF (V5-Tag antibodies) in mutCH12 cells is indicated by purple arrows. Black arrows show the status of CpG methylation at the lost CTCF sites in wtCH12 versus mutCH12 cells. **(E)** Heatmaps of CTCF, H2A.Z, and H3K9me3 ChIP-seq data in wtCH12 cells versus mutCH12 cells at the 5K lost CTCF sites, centered on the CTCF motifs in the plus (sense) orientation. **(F)** Profile of H3K9me3 ChIP-seq data at the 199 irreversibly lost CTCF sites in wtCH12 cells (black) versus mutCH12 cells (green), centered at CTCF motifs. The green arrow highlights a gain of H3K9me3 in mutCH12 cells compared to wtCH12 cells.

Previous studies have shown that CTCF binding is essential for maintaining the unmethylated state of its target sites (71,82). However, at the 5K lost CTCF sites, these regions largely remain unmethylated even in the absence of CTCF. Analysis of ATAC-seq data (Figure 3D) showed that the chromatin surrounding the 5K sites transitions to a closed state following CTCF loss, thereby preventing other chromatin-binding factors from sustaining the unmethylated state of these sites. This raises the question of why only a small subset (199 CTCF sites) of the lost 5K CTCF sites gained CpG methylation, while the remaining 4,958 sites remained largely methylation-free. To investigate this, we analyzed the sequences of the two groups (#2 and #3, Figure 7B) of CTCF sites to determine whether the irreversibly lost CTCF sites might have a higher presence of CpG sites within the consensus sequence, as previously reported (30). No significant differences were observed between these two groups that could explain their differential methylation status (Figure 7C). Furthermore, the irreversibly lost CTCF sites were not associated with CpG islands, with only 3.2% of them being part of such regions. A manual analysis of the most representative examples of irreversibly lost CTCF sites revealed that these 199 sites already exhibited some level of CpG methylation in wtCH12 cells, particularly in the sequences surrounding the CTCF-bound sites (Figure 7B, D and Supplementary Figure S12B, shown by black arrows). This contrasts sharply with the 5K lost sites, which could be restored with FL-CTCF expression and were generally devoid of CpG methylation in both wtCH12 and mutCH12 cells (Figure 7B, D).

Thus, certain CTCF sites are more susceptible to acquiring CpG methylation upon the loss of CTCF binding. Based on the observed CpG methylation enrichment (Figure 7B,D and Supplementary Figure S12B), these sites may initially be hemi-methylated (with only one strand of the DNA being methylated, as shown in Supplementary Figure S12B). Over time, with the loss of CTCF binding, these sites are more likely to become fully CpG methylated, which renders them inaccessible to CTCF binding.

As we have shown that the epigenetic status of CPN influences CTCF occupancy (Figure 1D and Supplementary Figures S2, S3A, S4C, S5D), we analyzed the enrichment patterns of H3K9me3 and H2A.Z around CTCF sites in wtCH12 and mutCH12 cells. A comparison of H3K9me3 levels at the remaining and lost CTCF sites in these cells showed that the epigenetic chromatin status did not change with the deletion of ZFs9-11, nor with the loss of CTCF occupancy at the 5K sites (Figure 7E and Supplementary Figure S12C). This may explain why these 5K lost CTCF sites remain available for future CTCF occupancy in mutCH12 cells. However, the 199 irreversibly lost CTCF sites did acquire the repressive histone mark, rendering these sites inaccessible for CTCF occupancy (Figure 7F). As high CTCF occupancy was associated with the gain of the H2A.Z histone variant (Figure 1D, Supplementary Figure S3A), we investigated whether H2A.Z pre-marks the 5K lost CTCF sites in mutCH12 cells for future CTCF occupancy, as previously suggested (46). However, analysis of H2A.Z in wtCH12 versus mutCH12 cells revealed that this histone variant was lost following the loss of CTCF occupancy at the 5K sites in mutCH12 cells (Figure 7E, Supplementary Figure S12D). This finding supports the conclusion that the absence of repressive histone modifications may maintain the 5K sites in a state permissive for future CTCF binding in mutCH12 cells, even though active histone variants like H2A.Z are completely lost.

Thus, in this study, we confirmed our initial hypothesis regarding the regulation of CTCF occupancy by CPN. We demonstrated that CTCF binding to the 5K lost CTCF sites in mutCH12 cells is preprogrammed not only by the CpG methylation status of the CTCF site itself but also by the CpG methylation status and histone modifications of the CPN. The preservation of the unchanged CpG methylation status and H3K9me3 histone modifications at the 5K lost CTCF sites in mutCH12 cells, as observed in wtCH12 cells, makes these sites available for CTCF occupancy (Figure 7). This finding highlights the importance of the epigenetic landscape in regulating CTCF binding and further emphasizes the role of chromatin accessibility in CTCF function.

## Discussion

CTCF binding patterns are closely linked to cell fate during differentiation, playing a critical role in 3D genome organization and cell-specific gene transcription (71,83). In this study, we sought to investigate how these cell-specific CTCF binding patterns are established and maintained to preserve cell identity. First, we compared CTCF binding to naked DNA versus chromatin and proposed a model for CTCF occupancy. According to this model, CTCF binding sites are strategically positioned at the entry side of a well-positioned nucleosome, such that upon CTCF binding, its N-terminus is oriented towards the nucleosome. We refer to this nucleosome as the CTCF-priming nucleosome (CPN). CTCF can either shift this nucleosome downstream from its binding site or not, depending on the epigenetic state of the nucleosome.

CTCF displaces nucleosomes when the DNA is unmethylated at CpG sites and the nucleosomes lack repressive histone marks, such as H3K9me3. Under these conditions, CTCF recruits SMARCA5 via its N-terminus, pushing the nucleosome downstream and enabling stable binding to the nucleosome-free sequence. Conversely, when nucleosomal DNA is CpG-methylated and enriched with repressive histone marks such as H3K9me3, CTCF cannot effectively displace the nucleosome. In such cases, the nucleosome remains in place, leading to weak and transient CTCF binding. Based on our analysis of the experimental data presented here and multiple published datasets, we propose that the contributions of the C-terminal and N-terminal nucleosomes flanking CTCF are unequal. Specifically, the epigenetic status of the N-terminal nucleosome (CPN) plays a more critical role in determining CTCF binding. This distinction is further supported by differences in their positioning relative to the CTCF binding site: 1). upon CTCF binding, the N-terminal nucleosome is shifted further downstream compared to the C-terminal nucleosome (75); 2). mapping CTCF occupancy using the CUT&RUN approach often results in the asymmetric release of the C-terminal nucleosome along with the CTCF footprint (84), suggesting that the C-terminal nucleosome is positioned closer to the CTCF binding site.

To confirm our model, we used wtCH12 and mutCH12 cells and demonstrated that as long as the CPN’s epigenetic status remains favorable for CTCF binding, these sites remain pre-marked for CTCF occupancy, even in the absence of CTCF itself. Our data suggest that both the CTCF consensus sequence and the CPN must be free of repressive epigenetic marks to permit stable CTCF binding. Active epigenetic marks, such as H2A.Z histone variant, appear to be linked to CTCF binding itself rather than serving as pre-markers for future CTCF occupancy. We also speculate that the mechanism by which priming nucleosomes regulate CTCF occupancy may extend to other pioneer factors, as REST (85), PU.1 (86), and ESRRB (87), which are capable of displacing nucleosomes and associating with well-positioned nucleosomes. However, the exact mechanism of nucleosome displacement likely varies, depending on the specific protein partners of each factor.

### Do small CTCF peaks matter?

Cross-referencing ChIP-seq (*in vivo*) with Affinity-seq (*in vitro*) from the same cell type revealed that CTCF sites detected strongly *in vitro* but not *in vivo* are still transiently bound by CTCF *in vivo*. However, this transient CTCF occupancy is often insufficient for these sites to be classified as significantly bound by standard peak-calling algorithms, such as MACS. Similarly, when comparing CTCF occupancy across different cell lines, a closer examination of cell-specific CTCF binding sites shows that these sites are also transiently bound by CTCF in other cell types but are not identified as significant peaks in ChIP-seq data.

In Figure 1C-D and Supplementary Figures S1-S6, we have highlighted these low-occupancy CTCF sites with yellow arrows. According to our model, CTCF can transiently recognize and bind to these sites without inducing chromatin opening. The repressive nature of the CPN, which incorporates the H3K9me3 repressive histone mark, prevents strong and sustained CTCF occupancy. Thus, we propose that there are many more CTCF binding sites than are currently detected by ChIP-seq using peak-calling algorithms like MACS.

Do these low-occupancy CTCF sites matter? Our data suggest they do. We observed that low CTCF occupancy is associated with low cohesin enrichment, indicating that these sites might contribute to loop formation, albeit transiently and for short durations. A growing body of research supports the idea that low-occupancy CTCF sites play a role in cell-specific transcription (88), serve as targets for JPX RNA (89). Additionally, recent findings indicate that weak CTCF sites help define H3K9me2/3 nanodomain structures within the genome (90). In summary, these low-occupancy CTCF sites should not be disregarded, as they may contribute to 3D genome organization and gene transcription.

### Why does the enrichment of H3K9me3 and H2A.Z at the CPN affect CTCF occupancy?

In this study, we demonstrated that the presence of the H3K9me3 histone modification on the CTCF priming nucleosome (CPN) either reduces or completely inhibits CTCF binding to chromatin. This finding is consistent with several previous studies that have identified a mutually exclusive relationship between H3K9me3 enrichment and CTCF occupancy (28,90–92). For example, during the differentiation of human embryonic stem cells into pancreatic islet organoids, CTCF recruitment to new binding sites is facilitated by the demethylation of H3K9me3 to H3K9me2 on nucleosomes flanking these sites (91). The H3K9me3 modification can be established by at least six enzymes, including SUV39H1, SUV39H2, SETDB1, SETDB2, G9A, and GLP, which exhibit redundant functions (93). The single knockout of several of these enzymes has been shown to increase CTCF occupancy. For instance, SETDB1 depletion in mouse cells led to the increased recruitment of CTCF at more than 1,600 CTCF motifs within SINE B2 repeats (94). Similar results confirmed that SETDB1 knockout caused a global increase in CTCF occupancy, accompanied by changes in nucleosome positioning around newly occupied CTCF sites, independent of DNA methylation (28). Furthermore, G9A knockout also resulted in enhanced CTCF occupancy (95).

Thus, the removal of H3K9me3 consistently increases CTCF occupancy at thousands of CTCF binding sites, though not universally. This pattern mirrors that observed with CpG methylation, which influences CTCF occupancy but is not sufficient to explain its genome-wide distribution (96). We speculate that the CPN may harbor additional repressive histone marks not mapped in this study.

The reduced CTCF occupancy at sites flanked by H3K9me3-modified nucleosomes suggests that CTCF may bind these sites transiently but is unable to reposition the CPN. As previously shown (27,44), and confirmed by our study, SMARCA5 (SNF2H) is required for CTCF to displace nucleosomes at its binding sites. Interestingly, histone deacetylation has been shown to block SWI/SNF remodelers from accessing chromatin (97), suggesting that repressive histone modifications may hinder SMARCA5-mediated nucleosome displacement, thereby preventing the strong and stable occupancy of CTCF. Conversely, active histone variants such as H2A.Z may facilitate nucleosome displacement by SMARCA5 (69), promoting chromatin opening around CTCF sites and enhancing CTCF binding stability.

A key question remains: what initiates strong CTCF occupancy? In this study, we demonstrated that the grammar of the CTCF core motif plays a critical role. The affinity of CTCF binding varies depending on the strength of the CTCF motif. CTCF likely demonstrates higher affinity for stronger motifs and, upon binding, influences the epigenetic status of flanking nucleosomes by incorporating H2A.Z through the recruitment of chromatin remodeling factors, such as SRCAP (24). This incorporation, in turn, further facilitates chromatin opening via SMARCA5 activity. In contrast, weaker CTCF motifs are more susceptible to cell-specific regulation of CTCF occupancy through repressive epigenetic modifications, as H3K9me3 and CpG methylation.

### Why is the CTCF priming nucleosome well-positioned relative to the CTCF consensus?

Our proposed model supports the CTCF-nucleosome competition model, which has been explored in prior studies (36,40,41). An analysis of MNase-seq data from experiments where CTCF was depleted using either the auxin-degron system (57) or a conditional mouse model (98), or where CTCF binding was lost during cell differentiation (42), revealed increased nucleosome occupancy directly at CTCF binding sites in the absence of CTCF. In this study, by focusing on CTCF sites in the plus orientation, we found that the CTCF consensus typically resides on the entry side of the well-positioned nucleosome, with the N-terminus of CTCF facing the nucleosome upon binding.

The next question is: Why is the CTCF priming nucleosome well-positioned? The fact that nucleosomes remain well-positioned even in the absence of CTCF suggests that intrinsic DNA properties, such as DNA conformation and bendability, play a key role in nucleosome positioning. Indeed, CTCF binding sequences have been shown to possess properties that enable them to stall RNA polymerase II independently of CTCF binding (99) and help retain cohesin at CTCF-bound sites (48). Notably, even when CTCF’s N-terminus, which interacts with cohesin, is directed to alternative target sites, cohesin is not retained at non-CTCF sequences (48,80), indicating that the specific conformation of the CTCF-DNA complex is crucial. Moreover, the CTCF motif sequence itself has been found to exhibit significantly higher DNA bendability compared to the surrounding 200 bp of DNA (100). DNA bendability is a key factor in nucleosome positioning, and sequences with greater flexibility often favor specific rotational settings within nucleosomes (101). Thus, the high flexibility of the DNA at CTCF consensus sites may naturally favor its positioning near the entry side of the nucleosome.

Another potential explanation for the well-positioned nucleosome could be related to non-B DNA motifs, which are found to be highly enriched at CTCF sites (102,103). Non-B DNA structures are known to disrupt nucleosome positioning (104). If CTCF binding sites adopt a non-B DNA structure, this could act as a repellent for nucleosome assembly, further contributing to the shift in nucleosome positioning relative to the CTCF site.

In summary, our study underscores the critical role of the well-positioned nucleosome in establishing and maintaining CTCF binding patterns, which are crucial for cell identity. We propose that CTCF binding is tightly regulated by the epigenetic state of the priming nucleosome. The next question is: What factors guide and establish these epigenetic marks during cell differentiation?

### Limitations of the Study

All analyses in this study were performed on total cell populations. While informative, single-molecule approaches could provide more detailed insights into the nucleosome repositioning dynamics associated with CTCF binding to chromatin. In investigating CTCF sites bound in vitro but not in vivo, we relied on multiple ENCODE datasets and other published ChIP-seq data for various epigenetic marks to explore the differences between chromatin-bound and DNA-only sites. However, we cannot rule out the potential influence of additional, yet-undiscovered epigenetic modifications that may contribute to these differences. Additionally, the analysis of wild-type versus mutant CH12 cells focused exclusively on the 5K CTCF sites with upstream CTCF consensus motifs bound by ZFs9-11. This scope may not fully capture the broader diversity of CTCF binding sites across the genome.

## Data availability

All raw and processed sequencing data used in this study are available through NCBI GEO database (GSE277826).

## Supplementary data

Supplementary Figures S1-S12.

## Acknowledgments

This study used the Office of Cyber Infrastructure and Computational Biology High Performance Computing cluster at NIAID, and high-performance computational capabilities of the Biowulf Linux cluster at NIH. We would like to express our sincere gratitude to the four anonymous reviewers whose insightful comments significantly enhanced the quality of our study. Author Contributions: M.T., D.N.B., and Y.J. conducted the experiments. E.M.P. and L.R. performed the bioinformatics analyses. M.T. and E.M.P. conceived and designed the project. V.V.L. provided overall supervision and support. E.M.P. wrote the paper with contributions and corrections from M.T., D.N.B., E.P., Y.J., D.L., V.B.T. All authors reviewed the manuscript.

## References

1. de Wit, E., Vos, E.S., Holwerda, S.J., Valdes-Quezada, C., Verstegen, M.J., Teunissen, H., Splinter, E., Wijchers, P.J., Krijger, P.H. and de Laat, W. (2015) CTCF Binding Polarity Determines Chromatin Looping. Mol Cell, 60, 676–684.

2. Rao, S.S., Huntley, M.H., Durand, N.C., Stamenova, E.K., Bochkov, I.D., Robinson, J.T., Sanborn, A.L., Machol, I., Omer, A.D., Lander, E.S. et al. (2014) A 3D map of the human genome at kilobase resolution reveals principles of chromatin looping. Cell, 159, 1665–1680.

3. Klenova, E.M., Nicolas, R.H., Paterson, H.F., Carne, A.F., Heath, C.M., Goodwin, G.H., Neiman, P.E. and Lobanenkov, V.V. (1993) CTCF, a conserved nuclear factor required for optimal transcriptional activity of the chicken c-myc gene, is an 11-Zn-finger protein differentially expressed in multiple forms. Mol Cell Biol, 13, 7612–7624.

4. Kim, T.H., Abdullaev, Z.K., Smith, A.D., Ching, K.A., Loukinov, D.I., Green, R.D., Zhang, M.Q., Lobanenkov, V.V. and Ren, B. (2007) Analysis of the vertebrate insulator protein CTCF-binding sites in the human genome. Cell, 128, 1231–1245.

5. Chen, H., Tian, Y., Shu, W., Bo, X. and Wang, S. (2012) Comprehensive identification and annotation of cell type-specific and ubiquitous CTCF-binding sites in the human genome. PLoS One, 7, e41374.

6. Ohlsson, R., Lobanenkov, V. and Klenova, E. (2010) Does CTCF mediate between nuclear organization and gene expression? Bioessays, 32, 37–50.

7. Bell, A.C., West, A.G. and Felsenfeld, G. (1999) The protein CTCF is required for the enhancer blocking activity of vertebrate insulators. Cell, 98, 387–396.

8. Kurukuti, S., Tiwari, V.K., Tavoosidana, G., Pugacheva, E., Murrell, A., Zhao, Z., Lobanenkov, V., Reik, W. and Ohlsson, R. (2006) CTCF binding at the H19 imprinting control region mediates maternally inherited higher-order chromatin conformation to restrict enhancer access to Igf2. Proc Natl Acad Sci U S A, 103, 10684–10689.

9. Lang, F., Li, X., Zheng, W., Li, Z., Lu, D., Chen, G., Gong, D., Yang, L., Fu, J., Shi, P. et al. (2017) CTCF prevents genomic instability by promoting homologous recombination-directed DNA double-strand break repair. Proc Natl Acad Sci U S A, 114, 10912–10917.

10. Phillips, J.E. and Corces, V.G. (2009) CTCF: master weaver of the genome. Cell, 137, 1194–1211.

11. Filippova, G.N., Thienes, C.P., Penn, B.H., Cho, D.H., Hu, Y.J., Moore, J.M., Klesert, T.R., Lobanenkov, V.V. and Tapscott, S.J. (2001) CTCF-binding sites flank CTG/CAG repeats and form a methylation-sensitive insulator at the DM1 locus. Nat Genet, 28, 335–343.

12. Pugacheva, E.M., Tiwari, V.K., Abdullaev, Z., Vostrov, A.A., Flanagan, P.T., Quitschke, W.W., Loukinov, D.I., Ohlsson, R. and Lobanenkov, V.V. (2005) Familial cases of point mutations in the XIST promoter reveal a correlation between CTCF binding and pre-emptive choices of X chromosome inactivation. Hum Mol Genet, 14, 953–965.

13. Price, E., Fedida, L.M., Pugacheva, E.M., Ji, Y.J., Loukinov, D. and Lobanenkov, V.V. (2023) An updated catalog of CTCF variants associated with neurodevelopmental disorder phenotypes. Front Mol Neurosci, 16, 1185796.

14. Fang, C., Wang, Z., Han, C., Safgren, S.L., Helmin, K.A., Adelman, E.R., Serafin, V., Basso, G., Eagen, K.P., Gaspar-Maia, A. et al. (2020) Cancer-specific CTCF binding facilitates oncogenic transcriptional dysregulation. Genome Biol, 21, 247.

15. Chen, W., Zeng, Y.C., Achinger-Kawecka, J., Campbell, E., Jones, A.K., Stewart, A.G., Khoury, A. and Clark, S.J. (2024) Machine learning enables pan-cancer identification of mutational hotspots at persistent CTCF binding sites. Nucleic Acids Res, 52, 8086–8099.

16. Hansen, A.S. (2020) CTCF as a boundary factor for cohesin-mediated loop extrusion: evidence for a multi-step mechanism. Nucleus, 11, 132–148.

17. Liu, Y., Wan, X., Li, H., Chen, Y., Hu, X., Chen, H., Zhu, D., Li, C. and Zhang, Y. (2023) CTCF coordinates cell fate specification via orchestrating regulatory hubs with pioneer transcription factors. Cell Rep, 42, 113259.

18. Dixon, J.R., Jung, I., Selvaraj, S., Shen, Y., Antosiewicz-Bourget, J.E., Lee, A.Y., Ye, Z., Kim, A., Rajagopal, N., Xie, W. et al. (2015) Chromatin architecture reorganization during stem cell differentiation. Nature, 518, 331–336.

19. Stefanova, M.E., Ing-Simmons, E., Stefanov, S., Flyamer, I., Dorado Garcia, H., Schopflin, R., Henssen, A.G., Vaquerizas, J.M. and Mundlos, S. (2023) Doxorubicin Changes the Spatial Organization of the Genome around Active Promoters. Cells, 12.

20. Kantidze, O.L., Luzhin, A.V., Nizovtseva, E.V., Safina, A., Valieva, M.E., Golov, A.K., Velichko, A.K., Lyubitelev, A.V., Feofanov, A.V., Gurova, K.V. et al. (2019) The anti-cancer drugs curaxins target spatial genome organization. Nat Commun, 10, 1441.

21. Kanduri, C., Pant, V., Loukinov, D., Pugacheva, E., Qi, C.F., Wolffe, A., Ohlsson, R. and Lobanenkov, V.V. (2000) Functional association of CTCF with the insulator upstream of the H19 gene is parent of origin-specific and methylation-sensitive. Curr Biol, 10, 853–856.

22. Monteagudo-Sanchez, A., Noordermeer, D. and Greenberg, M.V.C. (2024) The impact of DNA methylation on CTCF-mediated 3D genome organization. Nat Struct Mol Biol, 31, 404–412.

23. Hark, A.T., Schoenherr, C.J., Katz, D.J., Ingram, R.S., Levorse, J.M. and Tilghman, S.M. (2000) CTCF mediates methylation-sensitive enhancer-blocking activity at the H19/Igf2 locus. Nature, 405, 486–489.

24. Pugacheva, E.M., Bhatt, D.N., Rivero-Hinojosa, S., Tajmul, M., Fedida, L., Price, E., Ji, Y., Loukinov, D., Strunnikov, A.V., Ren, B. et al. (2024) BORIS/CTCFL epigenetically reprograms clustered CTCF binding sites into alternative transcriptional start sites. Genome Biol, 25, 40.

25. Dephoure, N., Zhou, C., Villen, J., Beausoleil, S.A., Bakalarski, C.E., Elledge, S.J. and Gygi, S.P. (2008) A quantitative atlas of mitotic phosphorylation. Proc Natl Acad Sci U S A, 105, 10762–10767.

26. Ramirez-Cuellar, J., Ferrari, R., Sanz, R.T., Valverde-Santiago, M., Garcia-Garcia, J., Nacht, A.S., Castillo, D., Le Dily, F., Neguembor, M.V., Malatesta, M., et al. (2024) LATS1 controls CTCF chromatin occupancy and hormonal response of 3D-grown breast cancer cells. EMBO J, 43, 1770–1798.

27. Bomber, M.L., Wang, J., Liu, Q., Barnett, K.R., Layden, H.M., Hodges, E., Stengel, K.R. and Hiebert, S.W. (2023) Human SMARCA5 is continuously required to maintain nucleosome spacing. Mol Cell, 83, 507–522 e506.

28. Tam, P.L.F., Cheung, M.F., Chan, L.Y. and Leung, D. (2024) Cell-type differential targeting of SETDB1 prevents aberrant CTCF binding, chromatin looping, and cis-regulatory interactions. Nat Commun, 15, 15.

29. Weth, O. and Renkawitz, R. (2011) CTCF function is modulated by neighboring DNA binding factors. Biochem Cell Biol, 89, 459–468.

30. Hashimoto, H., Wang, D., Horton, J.R., Zhang, X., Corces, V.G. and Cheng, X. (2017) Structural Basis for the Versatile and Methylation-Dependent Binding of CTCF to DNA. Mol Cell, 66, 711–720 e713.

31. Filippova, G.N., Fagerlie, S., Klenova, E.M., Myers, C., Dehner, Y., Goodwin, G., Neiman, P.E., Collins, S.J. and Lobanenkov, V.V. (1996) An exceptionally conserved transcriptional repressor, CTCF, employs different combinations of zinc fingers to bind diverged promoter sequences of avian and mammalian c-myc oncogenes. Mol Cell Biol, 16, 2802–2813.

32. Nakahashi, H., Kieffer Kwon, K.R., Resch, W., Vian, L., Dose, M., Stavreva, D., Hakim, O., Pruett, N., Nelson, S., Yamane, A. et al. (2013) A genome-wide map of CTCF multivalency redefines the CTCF code. Cell Rep, 3, 1678–1689.

33. Fu, Y., Sinha, M., Peterson, C.L. and Weng, Z. (2008) The insulator binding protein CTCF positions 20 nucleosomes around its binding sites across the human genome. PLoS Genet, 4, e1000138.

34. Stergachis, A.B., Debo, B.M., Haugen, E., Churchman, L.S. and Stamatoyannopoulos, J.A. (2020) Single-molecule regulatory architectures captured by chromatin fiber sequencing. Science, 368, 1449–1454.

35. Lai, B., Gao, W., Cui, K., Xie, W., Tang, Q., Jin, W., Hu, G., Ni, B. and Zhao, K. (2018) Principles of nucleosome organization revealed by single-cell micrococcal nuclease sequencing. Nature, 562, 281–285.

36. Sonmezer, C., Kleinendorst, R., Imanci, D., Barzaghi, G., Villacorta, L., Schubeler, D., Benes, V., Molina, N. and Krebs, A.R. (2021) Molecular Co-occupancy Identifies Transcription Factor Binding Cooperativity In Vivo. Mol Cell, 81, 255–267 e256.

37. Hansen, A.S., Pustova, I., Cattoglio, C., Tjian, R. and Darzacq, X. (2017) CTCF and cohesin regulate chromatin loop stability with distinct dynamics. Elife, 6.

38. Soochit, W., Sleutels, F., Stik, G., Bartkuhn, M., Basu, S., Hernandez, S.C., Merzouk, S., Vidal, E., Boers, R., Boers, J. et al. (2021) CTCF chromatin residence time controls three-dimensional genome organization, gene expression and DNA methylation in pluripotent cells. Nat Cell Biol, 23, 881–893.

39. Mach, P., Kos, P.I., Zhan, Y., Cramard, J., Gaudin, S., Tunnermann, J., Marchi, E., Eglinger, J., Zuin, J., Kryzhanovska, M. et al. (2022) Cohesin and CTCF control the dynamics of chromosome folding. Nat Genet, 54, 1907–1918.

40. Voong, L.N., Xi, L., Sebeson, A.C., Xiong, B., Wang, J.P. and Wang, X. (2016) Insights into Nucleosome Organization in Mouse Embryonic Stem Cells through Chemical Mapping. Cell, 167, 1555–1570 e1515.

41. Wiehle, L., Thorn, G.J., Raddatz, G., Clarkson, C.T., Rippe, K., Lyko, F., Breiling, A. and Teif, V.B. (2019) DNA (de)methylation in embryonic stem cells controls CTCF-dependent chromatin boundaries. Genome Res, 29, 750–761.

42. Clarkson, C.T., Deeks, E.A., Samarista, R., Mamayusupova, H., Zhurkin, V.B. and Teif, V.B. (2019) CTCF-dependent chromatin boundaries formed by asymmetric nucleosome arrays with decreased linker length. Nucleic Acids Res, 47, 11181–11196.

43. Marino, M.M., Rega, C., Russo, R., Valletta, M., Gentile, M.T., Esposito, S., Baglivo, I., De Feis, I., Angelini, C., Xiao, T., et al. (2019) Interactome mapping defines BRG1, a component of the SWI/SNF chromatin remodeling complex, as a new partner of the transcriptional regulator CTCF. J Biol Chem, 294, 861–873.

44. Wiechens, N., Singh, V., Gkikopoulos, T., Schofield, P., Rocha, S. and Owen-Hughes, T. (2016) The Chromatin Remodelling Enzymes SNF2H and SNF2L Position Nucleosomes adjacent to CTCF and Other Transcription Factors. PLoS Genet, 12, e1005940.

45. Barisic, D., Stadler, M.B., Iurlaro, M. and Schubeler, D. (2019) Mammalian ISWI and SWI/SNF selectively mediate binding of distinct transcription factors. Nature, 569, 136–140.

46. Oomen, M.E., Hansen, A.S., Liu, Y., Darzacq, X. and Dekker, J. (2019) CTCF sites display cell cycle-dependent dynamics in factor binding and nucleosome positioning. Genome Res, 29, 236–249.

47. Nora, E.P., Goloborodko, A., Valton, A.L., Gibcus, J.H., Uebersohn, A., Abdennur, N., Dekker, J., Mirny, L.A. and Bruneau, B.G. (2017) Targeted Degradation of CTCF Decouples Local Insulation of Chromosome Domains from Genomic Compartmentalization. Cell, 169, 930–944 e922.

48. Pugacheva, E.M., Kubo, N., Loukinov, D., Tajmul, M., Kang, S., Kovalchuk, A.L., Strunnikov, A.V., Zentner, G.E., Ren, B. and Lobanenkov, V.V. (2020) CTCF mediates chromatin looping via N-terminal domain-dependent cohesin retention. Proc Natl Acad Sci U S A, 117, 2020–2031.

49. Vian, L., Pekowska, A., Rao, S.S.P., Kieffer-Kwon, K.R., Jung, S., Baranello, L., Huang, S.C., El Khattabi, L., Dose, M., Pruett, N. et al. (2018) The Energetics and Physiological Impact of Cohesin Extrusion. Cell, 173, 1165–1178 e1120.

50. Buenrostro, J.D., Giresi, P.G., Zaba, L.C., Chang, H.Y. and Greenleaf, W.J. (2013) Transposition of native chromatin for fast and sensitive epigenomic profiling of open chromatin, DNA-binding proteins and nucleosome position. Nat Methods, 10, 1213–1218.

51. Zuo, Z., Billings, T., Walker, M., Petkov, P.M., Fordyce, P.M. and Stormo, G.D. (2023) On the dependent recognition of some long zinc finger proteins. Nucleic Acids Res, 51, 5364–5376.

52. Shen, Y., Yue, F., McCleary, D.F., Ye, Z., Edsall, L., Kuan, S., Wagner, U., Dixon, J., Lee, L., Lobanenkov, V.V. et al. (2012) A map of the cis-regulatory sequences in the mouse genome. Nature, 488, 116–120.

53. Cattoglio, C., Pustova, I., Walther, N., Ho, J.J., Hantsche-Grininger, M., Inouye, C.J., Hossain, M.J., Dailey, G.M., Ellenberg, J., Darzacq, X. et al. (2019) Determining cellular CTCF and cohesin abundances to constrain 3D genome models. Elife, 8.

54. Sundaram, V., Cheng, Y., Ma, Z., Li, D., Xing, X., Edge, P., Snyder, M.P. and Wang, T. (2014) Widespread contribution of transposable elements to the innovation of gene regulatory networks. Genome Res, 24, 1963–1976.

55. Wen, Z., Zhang, L., Ruan, H. and Li, G. (2020) Histone variant H2A.Z regulates nucleosome unwrapping and CTCF binding in mouse ES cells. Nucleic Acids Res, 48, 5939–5952.

56. Papin, C., Le Gras, S., Ibrahim, A., Salem, H., Karimi, M.M., Stoll, I., Ugrinova, I., Schroder, M., Fontaine-Pelletier, E., Omran, Z., et al. (2021) CpG Islands Shape the Epigenome Landscape. J Mol Biol, 433, 166659.

57. Owens, N., Papadopoulou, T., Festuccia, N., Tachtsidi, A., Gonzalez, I., Dubois, A., Vandormael-Pournin, S., Nora, E.P., Bruneau, B.G., Cohen-Tannoudji, M. et al. (2019) CTCF confers local nucleosome resiliency after DNA replication and during mitosis. Elife, 8.

58. Nakajima, T., Kanno, T., Ueda, Y., Miyako, K., Endo, T., Yoshida, S., Yokoyama, S., Asou, H.K., Yamada, K., Ikeda, K. et al. (2024) Fatty acid metabolism constrains Th9 cell differentiation and antitumor immunity via the modulation of retinoic acid receptor signaling. Cell Mol Immunol, 21, 1266–1281.

59. Fischer, V., Plassard, D., Ye, T., Reina-San-Martin, B., Stierle, M., Tora, L. and Devys, D. (2021) The related coactivator complexes SAGA and ATAC control embryonic stem cell self-renewal through acetyltransferase-independent mechanisms. Cell Rep, 36, 109598.

60. Langmead, B., Trapnell, C., Pop, M. and Salzberg, S.L. (2009) Ultrafast and memory-efficient alignment of short DNA sequences to the human genome. Genome Biol, 10, R25.

61. Li, H., Handsaker, B., Wysoker, A., Fennell, T., Ruan, J., Homer, N., Marth, G., Abecasis, G., Durbin, R. and Genome Project Data Processing, S. (2009) The Sequence Alignment/Map format and SAMtools. Bioinformatics, 25, 2078–2079.

62. Ramirez, F., Dundar, F., Diehl, S., Gruning, B.A. and Manke, T. (2014) deepTools: a flexible platform for exploring deep-sequencing data. Nucleic Acids Res, 42, W187–191.

63. Zhang, Y., Liu, T., Meyer, C.A., Eeckhoute, J., Johnson, D.S., Bernstein, B.E., Nusbaum, C., Myers, R.M., Brown, M., Li, W. et al. (2008) Model-based analysis of ChIP-Seq (MACS). Genome Biol, 9, R137.

64. Robinson, J.T., Thorvaldsdottir, H., Winckler, W., Guttman, M., Lander, E.S., Getz, G. and Mesirov, J.P. (2011) Integrative genomics viewer. Nat Biotechnol, 29, 24–26.

65. Quinlan, A.R. and Hall, I.M. (2010) BEDTools: a flexible suite of utilities for comparing genomic features. Bioinformatics, 26, 841–842.

66. Khan, A. and Mathelier, A. (2017) Intervene: a tool for intersection and visualization of multiple gene or genomic region sets. BMC Bioinformatics, 18, 287.

67. Bailey, T.L., Boden, M., Buske, F.A., Frith, M., Grant, C.E., Clementi, L., Ren, J., Li, W.W. and Noble, W.S. (2009) MEME SUITE: tools for motif discovery and searching. Nucleic Acids Res, 37, W202–208.

68. Abramson, J., Adler, J., Dunger, J., Evans, R., Green, T., Pritzel, A., Ronneberger, O., Willmore, L., Ballard, A.J., Bambrick, J. et al. (2024) Accurate structure prediction of biomolecular interactions with AlphaFold 3. Nature, 630, 493–500.

69. Goldman, J.A., Garlick, J.D. and Kingston, R.E. (2010) Chromatin remodeling by imitation switch (ISWI) class ATP-dependent remodelers is stimulated by histone variant H2A.Z. J Biol Chem, 285, 4645–4651.

70. Lehnertz, B., Ueda, Y., Derijck, A.A., Braunschweig, U., Perez-Burgos, L., Kubicek, S., Chen, T., Li, E., Jenuwein, T. and Peters, A.H. (2003) Suv39h-mediated histone H3 lysine 9 methylation directs DNA methylation to major satellite repeats at pericentric heterochromatin. Curr Biol, 13, 1192–1200.

71. Xiang, J.F. and Corces, V.G. (2021) Regulation of 3D chromatin organization by CTCF. Curr Opin Genet Dev, 67, 33–40.

72. Pugacheva, E.M., Rivero-Hinojosa, S., Espinoza, C.A., Mendez-Catala, C.F., Kang, S., Suzuki, T., Kosaka-Suzuki, N., Robinson, S., Nagarajan, V., Ye, Z. et al. (2015) Comparative analyses of CTCF and BORIS occupancies uncover two distinct classes of CTCF binding genomic regions. Genome Biol, 16, 161.

73. Loukinov, D.I., Pugacheva, E., Vatolin, S., Pack, S.D., Moon, H., Chernukhin, I., Mannan, P., Larsson, E., Kanduri, C., Vostrov, A.A. et al. (2002) BORIS, a novel male germ-line-specific protein associated with epigenetic reprogramming events, shares the same 11-zinc-finger domain with CTCF, the insulator protein involved in reading imprinting marks in the soma. Proc Natl Acad Sci U S A, 99, 6806–6811.

74. Weth, O., Paprotka, C., Gunther, K., Schulte, A., Baierl, M., Leers, J., Galjart, N. and Renkawitz, R. (2014) CTCF induces histone variant incorporation, erases the H3K27me3 histone mark and opens chromatin. Nucleic Acids Res, 42, 11941–11951.

75. Do, C., Jiang, G., Zappile, P., Heguy, A. and Skok, J.A. (2024) A genome wide code to define cell-type specific CTCF binding and chromatin organization. bioRxiv.

76. Song, Y., Liang, Z., Zhang, J., Hu, G., Wang, J., Li, Y., Guo, R., Dong, X., Babarinde, I.A., Ping, W. et al. (2022) CTCF functions as an insulator for somatic genes and a chromatin remodeler for pluripotency genes during reprogramming. Cell Rep, 39, 110626.

77. Li, Y., Haarhuis, J.H.I., Sedeno Cacciatore, A., Oldenkamp, R., van Ruiten, M.S., Willems, L., Teunissen, H., Muir, K.W., de Wit, E., Rowland, B.D., et al. (2020) The structural basis for cohesin-CTCF-anchored loops. Nature, 578, 472–476.

78. Nora, E.P., Caccianini, L., Fudenberg, G., So, K., Kameswaran, V., Nagle, A., Uebersohn, A., Hajj, B., Saux, A.L., Coulon, A. et al. (2020) Molecular basis of CTCF binding polarity in genome folding. Nat Commun, 11, 5612.

79. Nishana, M., Ha, C., Rodriguez-Hernaez, J., Ranjbaran, A., Chio, E., Nora, E.P., Badri, S.B., Kloetgen, A., Bruneau, B.G., Tsirigos, A. et al. (2020) Defining the relative and combined contribution of CTCF and CTCFL to genomic regulation. Genome Biol, 21, 108.

80. Do, C., Jiang, G., Cova, G., Katsifis, C.C., Narducci, D.N., Yang, J., Sakellaropoulos, T., Vidal, R., Lhoumaud, P., Tsirigos, A., et al. (2024) Brain and cancer associated binding domain mutations provide insight into CTCF’s relationship with chromatin and its ability to act as a chromatin organizer. Res Sq.

81. Iurlaro, M., Masoni, F., Flyamer, I.M., Wirbelauer, C., Iskar, M., Burger, L., Giorgetti, L. and Schubeler, D. (2024) Systematic assessment of ISWI subunits shows that NURF creates local accessibility for CTCF. Nat Genet, 56, 1203–1212.

82. Kemp, C.J., Moore, J.M., Moser, R., Bernard, B., Teater, M., Smith, L.E., Rabaia, N.A., Gurley, K.E., Guinney, J., Busch, S.E. et al. (2014) CTCF haploinsufficiency destabilizes DNA methylation and predisposes to cancer. Cell Rep, 7, 1020–1029.

83. Arzate-Mejia, R.G., Recillas-Targa, F. and Corces, V.G. (2018) Developing in 3D: the role of CTCF in cell differentiation. Development, 145.

84. Spector, B.M., Santana, J.F., Pufall, M.A. and Price, D.H. (2024) DFF-ChIP: a method to detect and quantify complex interactions between RNA polymerase II, transcription factors, and chromatin. Nucleic Acids Res, 52, e88.

85. Harwood, J.C., Kent, N.A., Allen, N.D. and Harwood, A.J. (2019) Nucleosome dynamics of human iPSC during neural differentiation. EMBO Rep, 20.

86. Barozzi, I., Simonatto, M., Bonifacio, S., Yang, L., Rohs, R., Ghisletti, S. and Natoli, G. (2014) Coregulation of transcription factor binding and nucleosome occupancy through DNA features of mammalian enhancers. Mol Cell, 54, 844–857.

87. Festuccia, N., Owens, N., Papadopoulou, T., Gonzalez, I., Tachtsidi, A., Vandoermel-Pournin, S., Gallego, E., Gutierrez, N., Dubois, A., Cohen-Tannoudji, M. et al. (2019) Transcription factor activity and nucleosome organization in mitosis. Genome Res, 29, 250–260.

88. Essien, K., Vigneau, S., Apreleva, S., Singh, L.N., Bartolomei, M.S. and Hannenhalli, S. (2009) CTCF binding site classes exhibit distinct evolutionary, genomic, epigenomic and transcriptomic features. Genome Biol, 10, R131.

89. Oh, H.J., Aguilar, R., Kesner, B., Lee, H.G., Kriz, A.J., Chu, H.P. and Lee, J.T. (2021) Jpx RNA regulates CTCF anchor site selection and formation of chromosome loops. Cell, 184, 6157–6173 e6124.

90. Thorn, G.J., Clarkson, C.T., Rademacher, A., Mamayusupova, H., Schotta, G., Rippe, K. and Teif, V.B. (2022) DNA sequence-dependent formation of heterochromatin nanodomains. Nat Commun, 13, 1861.

91. Lyu, X., Rowley, M.J., Kulik, M.J., Dalton, S. and Corces, V.G. (2023) Regulation of CTCF loop formation during pancreatic cell differentiation. Nat Commun, 14, 6314.

92. Jiang, Y., Loh, Y.E., Rajarajan, P., Hirayama, T., Liao, W., Kassim, B.S., Javidfar, B., Hartley, B.J., Kleofas, L., Park, R.B. et al. (2017) The methyltransferase SETDB1 regulates a large neuron-specific topological chromatin domain. Nat Genet, 49, 1239–1250.

93. Padeken, J., Methot, S.P. and Gasser, S.M. (2022) Establishment of H3K9-methylated heterochromatin and its functions in tissue differentiation and maintenance. Nat Rev Mol Cell Biol, 23, 623–640.

94. Gualdrini, F., Polletti, S., Simonatto, M., Prosperini, E., Pileri, F. and Natoli, G. (2022) H3K9 trimethylation in active chromatin restricts the usage of functional CTCF sites in SINE B2 repeats. Genes Dev, 36, 414–432.

95. Jiang, Q., Ang, J.Y.J., Lee, A.Y., Cao, Q., Li, K.Y., Yip, K.Y. and Leung, D.C.Y. (2020) G9a Plays Distinct Roles in Maintaining DNA Methylation, Retrotransposon Silencing, and Chromatin Looping. Cell Rep, 33, 108315.

96. Maurano, M.T., Wang, H., John, S., Shafer, A., Canfield, T., Lee, K. and Stamatoyannopoulos, J.A. (2015) Role of DNA Methylation in Modulating Transcription Factor Occupancy. Cell Rep, 12, 1184–1195.

97. Sahu, R.K., Dhakshnamoorthy, J., Jain, S., Folco, H.D., Wheeler, D. and Grewal, S.I.S. (2024) Nucleosome remodeler exclusion by histone deacetylation enforces heterochromatic silencing and epigenetic inheritance. Mol Cell.

98. Marina-Zarate, E., Rodriguez-Ronchel, A., Gomez, M.J., Sanchez-Cabo, F. and Ramiro, A.R. (2023) Low-affinity CTCF binding drives transcriptional regulation whereas high-affinity binding encompasses architectural functions. iScience, 26, 106106.

99. Luan, J., Xiang, G., Gomez-Garcia, P.A., Tome, J.M., Zhang, Z., Vermunt, M.W., Zhang, H., Huang, A., Keller, C.A., Giardine, B.M. et al. (2021) Distinct properties and functions of CTCF revealed by a rapidly inducible degron system. Cell Rep, 34, 108783.

100. Li, K., Carroll, M., Vafabakhsh, R., Wang, X.A. and Wang, J.P. (2022) DNAcycP: a deep learning tool for DNA cyclizability prediction. Nucleic Acids Res, 50, 3142–3154.

101. Biswas, A. and Basu, A. (2023) The impact of the sequence-dependent physical properties of DNA on chromatin dynamics. Curr Opin Struct Biol, 83, 102698.

102. Georgakopoulos-Soares, I., Victorino, J., Parada, G.E., Agarwal, V., Zhao, J., Wong, H.Y., Umar, M.I., Elor, O., Muhwezi, A., An, J.Y. et al. (2022) High-throughput characterization of the role of non-B DNA motifs on promoter function. Cell Genom, 2.

103. Hou, Y., Li, F., Zhang, R., Li, S., Liu, H., Qin, Z.S. and Sun, X. (2019) Integrative characterization of G-Quadruplexes in the three-dimensional chromatin structure. Epigenetics, 14, 894–911.

104. Kouzine, F., Wojtowicz, D., Baranello, L., Yamane, A., Nelson, S., Resch, W., Kieffer-Kwon, K.R., Benham, C.J., Casellas, R., Przytycka, T.M. et al. (2017) Permanganate/S1 Nuclease Footprinting Reveals Non-B DNA Structures with Regulatory Potential across a Mammalian Genome. Cell Syst, 4, 344–356 e347.

