## Supplemental Figures for "CTCF binding landscape is shaped by the epigenetic state of the N-terminal nucleosome in relation to CTCF motif orientation"

### Supplementary Figure S1

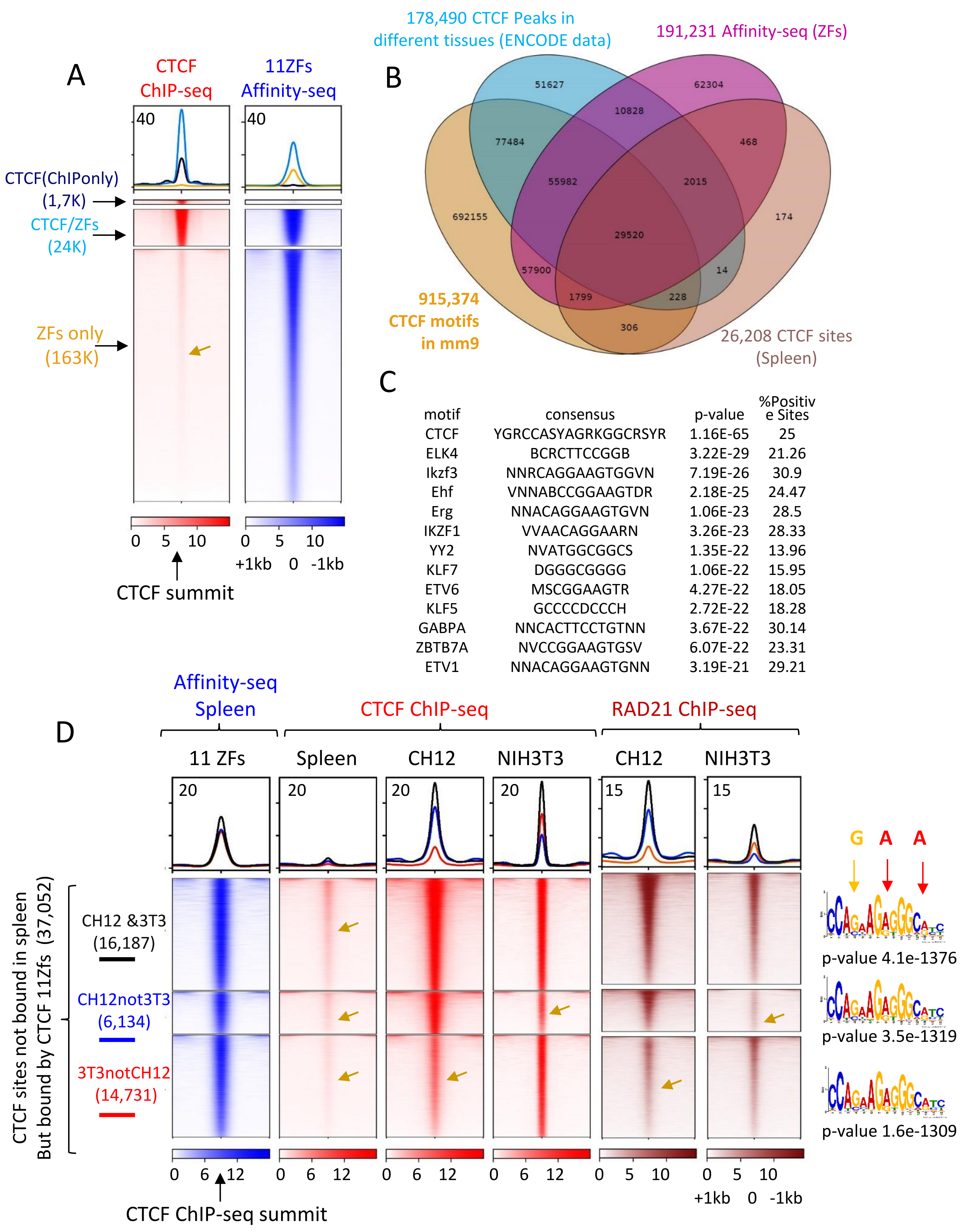


| motif | consensus | p-value | %Positive Sites |
|---|---|---|---|
| CTCF | YGRCCASYAGRKGGCRSYR | 1.16E-65 | 25 |
| ELK4 | BCRCTTCCGGB | 3.22E-29 | 21.26 |
| Ikzf3 | NNRCAGGAAGTGGVN | 7.19E-26 | 30.9 |
| Ehf | VNNABCCGGAAGTDR | 2.18E-25 | 24.47 |
| Erg | NNACAGGAAGTGVN | 1.06E-23 | 28.5 |
| IKZF1 | VVAACAGGAARN | 3.26E-23 | 28.33 |
| YY2 | NVATGGCGGCS | 1.35E-22 | 13.96 |
| KLF7 | DGGGCGGGG | 1.06E-22 | 15.95 |
| ETV6 | MSCGGAAGTR | 4.27E-22 | 18.05 |
| KLF5 | GCCCCDCCCH | 2.72E-22 | 18.28 |
| GABPA | NNCACTTCCTGTNN | 3.67E-22 | 30.14 |
| ZBTB7A | NVCCGGAAGTGSV | 6.07E-22 | 23.31 |
| ETV1 | NNACAGGAAGTGNN | 3.19E-21 | 29.21 |

**Supplementary Figure S1. Comparison of CTCF ChIP-seq and CTCF Affinity-seq data. (A)** Heatmap, based on the overlapping data from the main Figure 1B, is centered on the CTCF summit peak, showing CTCF binding detected by ChIP-seq (chromatin, red) and by Affinity-seq (naked DNA, blue) in mouse spleen cells. Approximately 1.7K CTCF sites were detected only by ChIP-seq, 24K CTCF sites were detected by both ChIP-seq and Affinity-seq (CTCF/ZFs), and approximately 163K CTCF sites were detected only by Affinity-seq (ZFs only). The low CTCF occupancy detected by ChIP-seq at Affinity-seq-only sites is indicated by yellow arrow. **(B)** Venn diagram showing the overlap of 915,374 15-bp CTCF motifs mapped in the genome (mm9) with CTCF ChIP-seq data from the indicated cell lines: CTCF ChIP-seq data combined from multiple cell types mapped by the ENCODE Project (178,490), with CTCF ChIP-seq in spleen (26,208), and with Affinity-seq data (191,231), performed on naked DNA from mouse spleen cells. **(C)** The list of transcriptional factors motif significantly enriched in 400bp window around the 1,7K CTCF sites mapped by ChIP-seq only (From Figure 1B). **(D)** Heatmaps centered on the CTCF summit peak showing that of the 37,052 CTCF binding sites detected by Affinity-seq only (11ZFs, blue) but not by CTCF ChIP-seq in mouse spleen (red), 16,187 are bound by CTCF in both CH12 and NIH3T3 cells (CH12&3T3), 6,134 are bound by CTCF only in CH12 cells (CH12not3T3), and 14,731 are bound by CTCF only in NIH3T3 cells (3T3notCH12). RAD21 (cohesin, brown) occupancy follows CTCF occupancy in a cell-specific pattern, with low but clear enrichment of reads at cell-specific CTCF ChIP-seq peaks not detected by MACS (indicated by yellow arrows). The low CTCF and RAD21 occupancy, not counted by ChIP-seq in mouse spleen or at cell-specific CTCF binding sites, is shown by yellow arrows. The CTCF core motif identified under cell-specific CTCF binding sites is shown on the right. At the top of the motif, yellow and red arrows indicate the presence of G, A, and A at positions 4, 8, and 13 within the 15-bp CTCF motif.

### Supplementary Figure S2

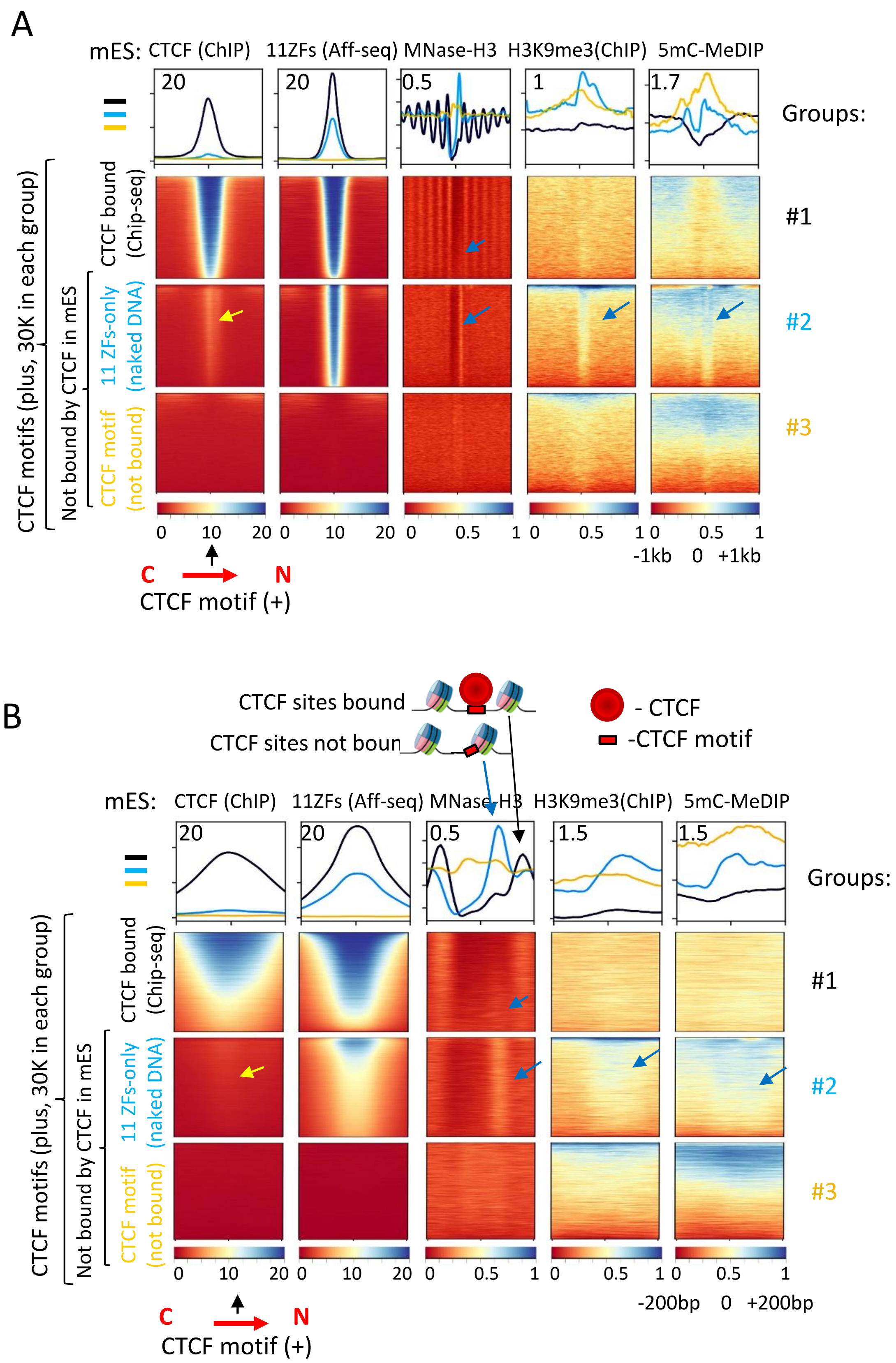

**Supplementary Figure S2. CTCF binding sites are strategically positioned at the entry site of well-positioned nucleosome in mEs cells. (A,B)** Heatmaps centered on the plus (sense) CTCF motif, comparing various chromatin features across three groups of CTCF binding sites in mES cells: #1 - CTCF-bound sites (black); #2 - CTCF motifs recovered by Affinity-seq but not identified as peaks by MACS in CTCF ChIP-seq in mES cells (blue); #3 - CTCF motifs neither bound by CTCF nor recovered by Affinity-seq (yellow). The heatmaps display the following data (from left to right): CTCF ChIP-seq (mES cells), CTCF Affinity-seq (spleen), MNase-H3 nucleosome positioning (mES cells), H3K9me3 ChIP-seq (mES cells), 5mC MeDIP-seq (mES cells). **(A)** A 2,000 bp window centered on the plus CTCF motif. **(B)** A zoomed-in view of the data from panel **(A)**, focusing on a 400 bp window centered on the plus CTCF motif. A schematic representation of bound and unbound CTCF sites is shown at the top, highlighting the relationship between CTCF binding and nucleosome positioning. The CTCF priming nucleosome (CPN) is indicated by a blue arrow.

### Supplementary Figure S3

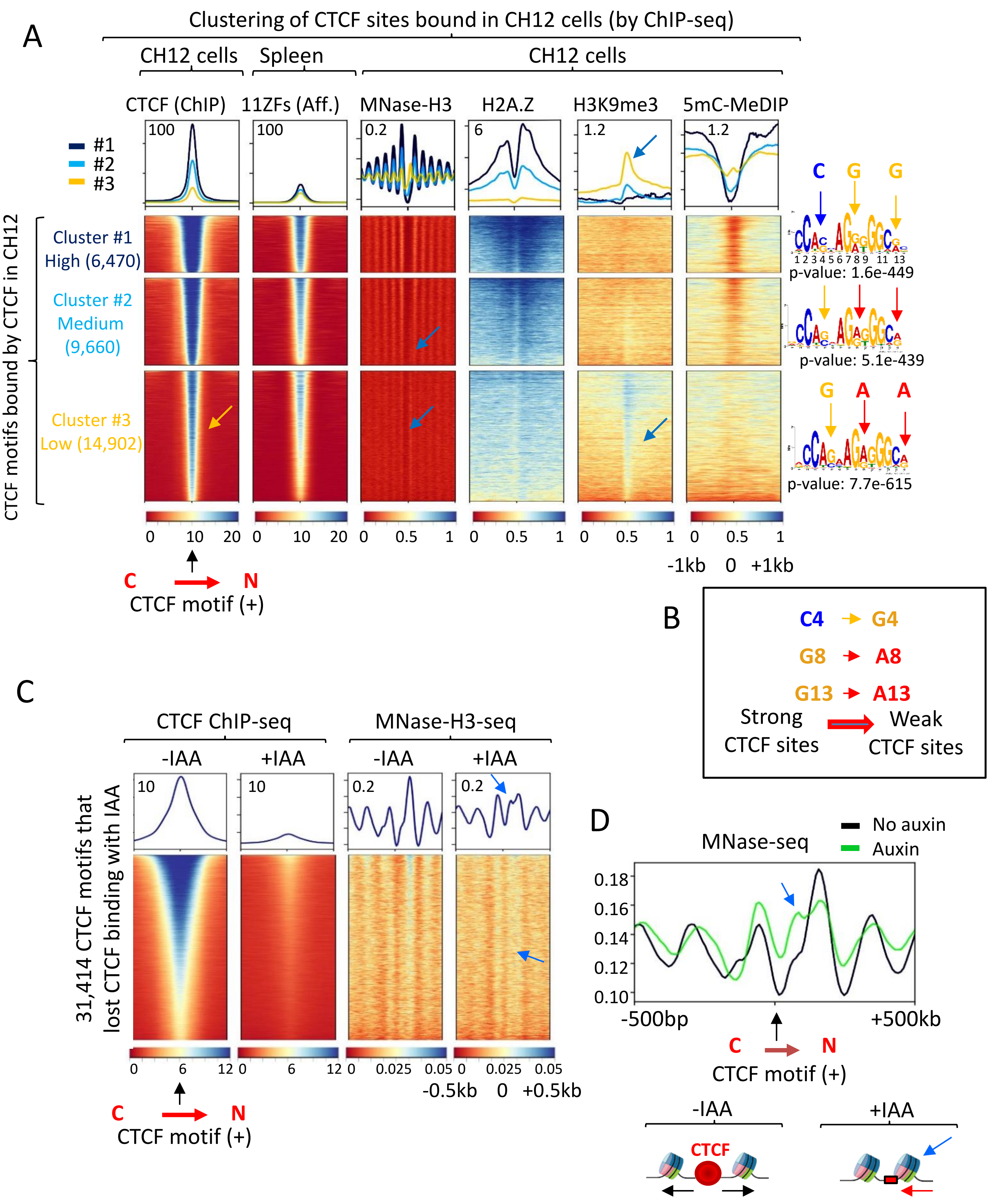

**Supplementary Figure S3. The N-terminal CTCF nucleosome affects CTCF binding in vivo. (A).** Heatmap centered on the plus-oriented CTCF motif of CTCF sites mapped by ChIP-seq in CH12 cells. The following datasets are displayed (from left to right): CTCF ChIP-seq (CH12 cells); CTCF Affinity-seq (spleen); MNase-H3 nucleosome positioning (CH12); H3K9me3 ChIP-seq (CH12); 5mC MeDIP-seq data (CH12 cells). The CTCF bound sites were classified into three clusters based on CTCF ChIP-seq tag enrichment: relatively high occupancy (black); medium occupancy (blue); lowest occupancy (yellow). On the right side of the heatmaps, CTCF motifs extracted from the corresponding clusters are shown. Yellow, blue, and red arrows indicate nucleotides at positions 4, 8, and 13 within the 14-bp CTCF consensus motif, where sequence variations correlate with differences in CTCF binding strength. **(B)** Summary of the CTCF motif "grammar" in relation to CTCF occupancy: High-occupancy CTCF-bound sites (Cluster #1) preferentially feature C, G, and G nucleotides at positions 4, 8, and 13, respectively; Low-occupancy, weak CTCF-bound sites (Clusters #2 and #3) contain G, A, and A nucleotides at these positions. **(C)** Heatmap centered on CTCF motifs of CTCF binding sites lost upon auxin (IAA)-mediated degradation of CTCF using auxin-inducible degron (AID) technology in mES cells. Only lost CTCF binding sites were included in the analysis. The following datasets are displayed (from left to right): CTCF ChIP-seq before and after auxin (IAA) treatment; MNase-H3-seq before and after auxin treatment. **(D)** Average plot showing nucleosome positioning (MNase-seq) around lost CTCF binding sites (motifs in plus orientation) before and after auxin treatment. At the bottom, a schematic representation of nucleosome shifts with CTCF loss is provided. The red arrow indicates the inward shift of the CTCF priming nucleosome (CPN) toward the CTCF motif. **(A, C, D)** The CTCF priming nucleosome (CPN) is marked by a blue arrow.

### Supplementary Figure S4

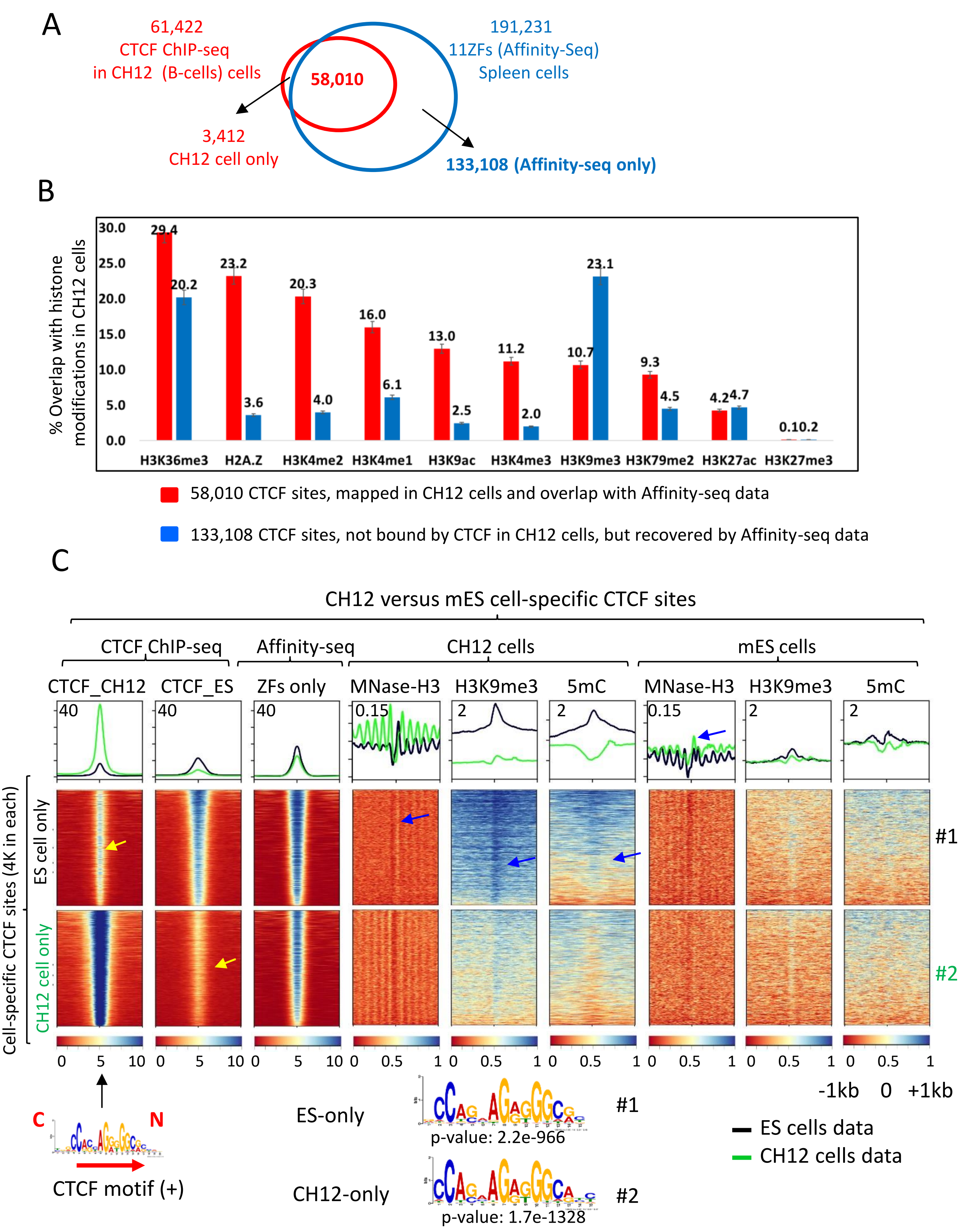

**Supplementary Figure S4. The epigenetic status of the N-terminus nucleosome affects CTCF occupancy in a cell-specific manner. (A)** Venn diagram showing the overlap of CTCF ChIP-seq in CH12 cells (red) with CTCF Affinity-seq data in spleen (blue). **(B)** Bar plot showing the percentage of overlapping for sites bound by CTCF (red) versus not bound by CTCF (blue) in CH12 cells (based on Panel A), analyzed in relation to various histone modifications mapped by ENCODE in CH12 cells. The difference between bound and unbound CTCF sites is statistically significant ($p$-value $< 0.001$) for all epigenetic marks examined, except H3K27ac. **(C)** Heatmaps centered on plus-oriented CTCF motifs, displaying (from left to right): CTCF ChIP-seq (CH12 cells), CTCF ChIP-seq (mES cells), CTCF Affinity-seq (spleen), MNase-H3 (CH12 cells), H3K9me3 ChIP-seq (CH12 cells), and 5mC MeDIP-seq data (CH12 cells), MNase-H3 (mES cells), H3K9me3 ChIP-seq (mES cells), and 5mC MeDIP-seq data (mES cells). These datasets are aligned across two groups of CTCF binding sites: (#1) CTCF sites bound in mES cells but not in CH12 cells (black), and (#2) CTCF sites bound in CH12 cells but not in mES cells (green). At the bottom of the heatmaps, CTCF motifs extracted from the corresponding clusters are shown. The CTCF priming nucleosome (CPN) is indicated by a blue arrow. The low CTCF ChIP-seq signal at the cell-specific CTCF binding sites is highlighted by a yellow arrow.

### Supplementary Figure S5

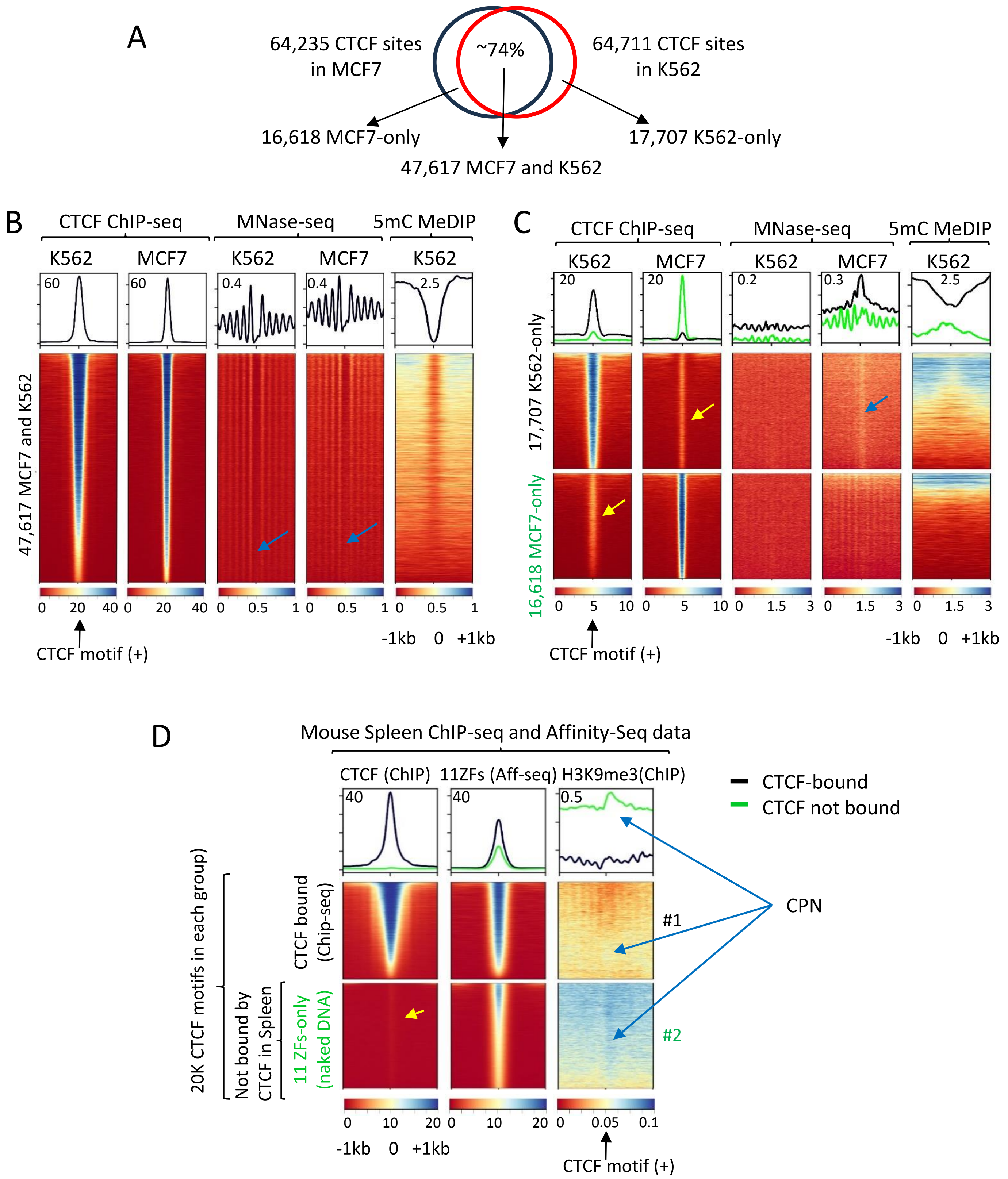

**Supplementary Figure S5. Cell-specific CTCF occupancy is regulated by the CTCF priming nucleosome**. **(A)** Venn diagram illustrating the overlap of CTCF ChIP-seq peaks mapped in K562 and MCF7 cells. **(B)** Heatmaps centered on CTCF plus motifs, displaying (from left to right): CTCF ChIP-seq data from K562 and MCF7 cells, MNase-seq data from K562 and MCF7 cells, 5mC MeDIP-seq data from K562 cells across CTCF sites, bound in both K562 and MCF7 cells. **(C)** Heatmaps showing the same datasets as panel **(B)** but aligned across two groups of cell-specific CTCF binding sites, identified in panel **(A)**: CTCF motifs bound only in K562 cells (black), CTCF motifs bound only in MCF7 cells (green). **(D)** Heatmaps centered on the plus-oriented CTCF motif, displaying (from left to right): CTCF ChIP-seq, CTCF Affinity-seq, H3K9me3 ChIP-seq. All datasets were obtained from mouse spleen cells. These data are aligned across two groups of CTCF binding sites: #1) CTCF-bound sites in mouse spleen cells (black), #2) CTCF motifs recovered by Affinity-seq but not identified as significant CTCF ChIP-seq peaks in spleen cells (green). For group #2, 20,000 CTCF sites were randomly selected to match the number of sites in group #1. **(B–D)** The CTCF priming nucleosome (CPN) is indicated by a blue arrow. The low CTCF ChIP-seq signal at cell-specific CTCF binding sites is highlighted by yellow arrows.

### Supplementary Figure S6

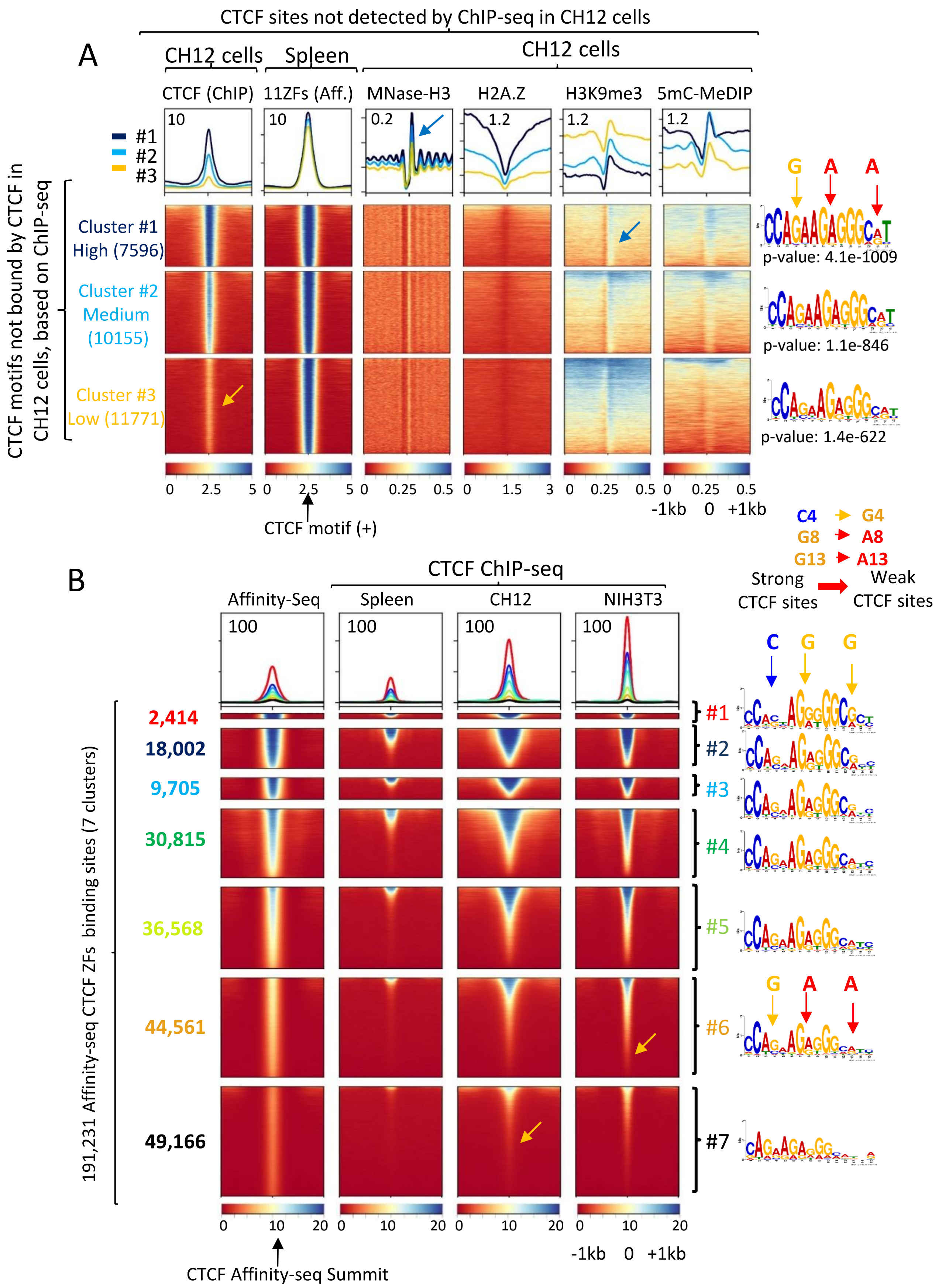

**Supplementary Figure S6. Motif and epigenetic analysis of CTCF sites recovered by Affinity-seq. (A)** Heatmap centered on the plus-oriented CTCF motif, showing CTCF binding sites recovered by Affinity-seq in spleen but not detected by ChIP-seq in CH12 cells. The following datasets are displayed (from left to right): CTCF ChIP-seq (CH12 cells); CTCF Affinity-seq (spleen); MNase-H3 nucleosome positioning (CH12); H3K9me3 ChIP-seq (CH12); 5mC MeDIP-seq data (CH12 cells). These CTCF binding sites, not detected by ChIP-seq in CH12 cells, are classified into three clusters based on CTCF ChIP-seq tag enrichment in CH12 cells: relatively high occupancy (black); medium occupancy (blue); lowest occupancy (yellow). The CTCF priming nucleosome (CPN) is indicated by a blue arrow. **(B)** Heatmap centered on the summit of 191,231 Affinity-seq mapped CTCF binding sites, classified into seven clusters based on Affinity-seq tag enrichment. The following datasets are displayed (from left to right): Affinity-seq (spleen); CTCF ChIP-seq in spleen, CH12, and NIH3T3 cells. The color of the clusters corresponds to the average plot color at the top of the heatmap. **(A,B)** On the right side of the heatmaps, CTCF motifs extracted from the corresponding clusters are shown. Yellow, blue, and red arrows indicate nucleotides at positions 4, 8, and 13 within the 14-bp CTCF consensus motif, where sequence variations correlate with differences in CTCF binding strength (related to Supplementary Figure S3B). The low CTCF ChIP-seq signal in the 11 zinc fingers-only group (#2) is highlighted by a yellow arrow.

### Supplementary Figure S7

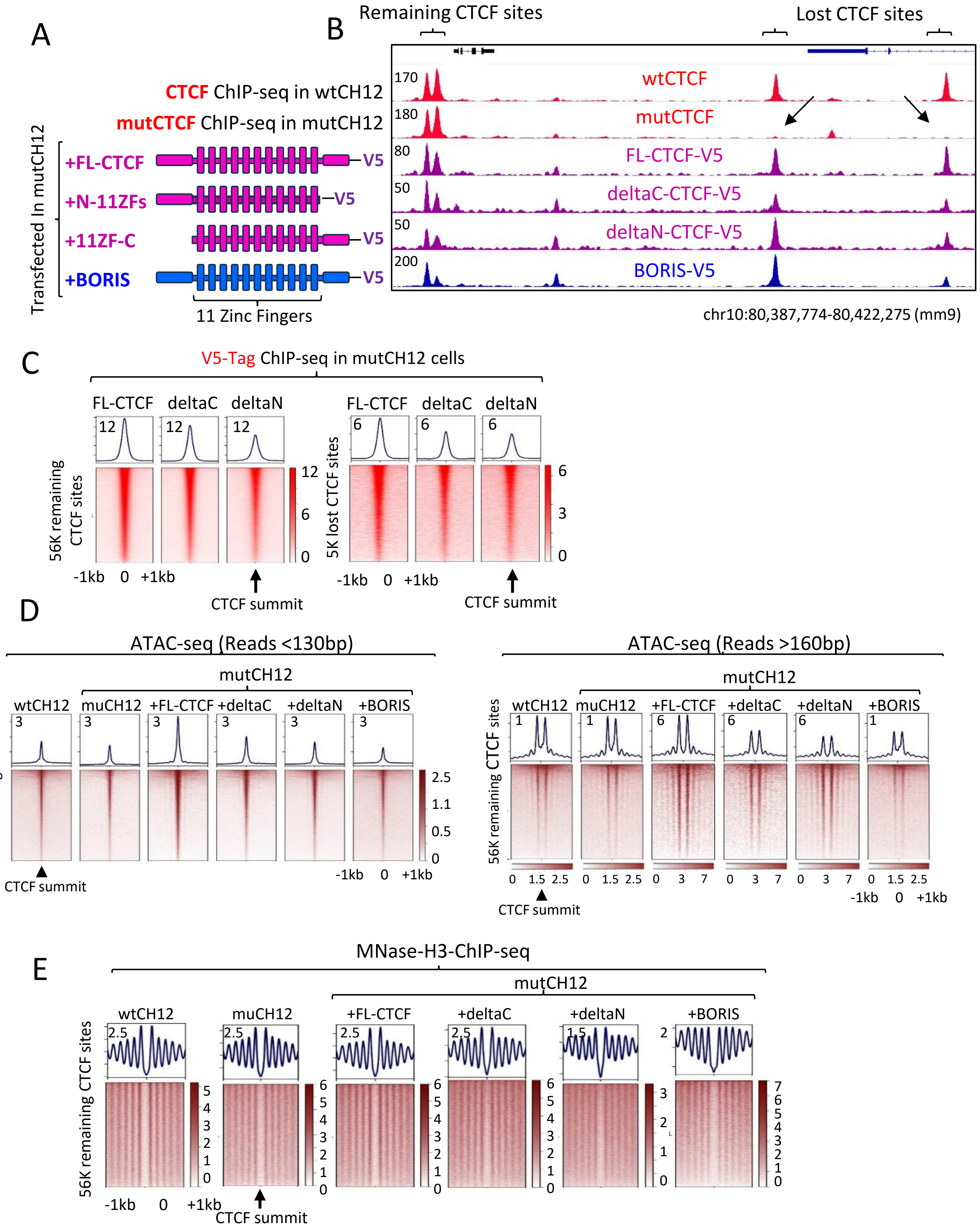

**Supplementary Figure S8. The use of wtCH12 and mutCH12 cells to study CTCF interaction with chromatin: Control data for Main Figures 3 and 4. (A)** Schematic representation of CTCF variants and BORIS ectopically expressed in mutCH12 cells. All ectopically expressed proteins were tagged with a V5-tag to allow mapping of their binding sites by ChIP-seq independent from endogenous CTCF. **(B)** Genome browser view showing examples of the 5K CTCF binding sites lost in mutCH12 cells, which were restored by ectopic expression of different CTCF variants and BORIS, as depicted in panel **(A).** The lost CTCF sites in mutCH12 cells are shown by black arrows**. (C)** Heatmap displaying V5-tag density for ectopically expressed CTCF vectors in mutCH12 cells. Left and right panels show a tag density at the remaining and the lost CTCF sites in mutCH12 cells, respectively. **(D)** Heatmaps depicting ATAC-seq density at remaining CTCF sites in wtCH12, mutCH12 cells, and mutCH12 cells ectopically expressing different CTCF vectors and BORIS (as indicated at the top of the heatmaps). Paired-end ATAC-seq reads were classified into short fragments (<130 bp), representing the CTCF footprint (Left panel), and nucleosome-sized fragments (>160 bp), reflecting nucleosome positioning (Right panel). **(E)** Heatmap showing MNase-seq-assisted H3 ChIP sequencing (MNase-H3) at the remaining CTCF sites in wtCH12 and mutCH12 cells. The heatmaps are labeled as in panel D.

### Supplementary Figure S8

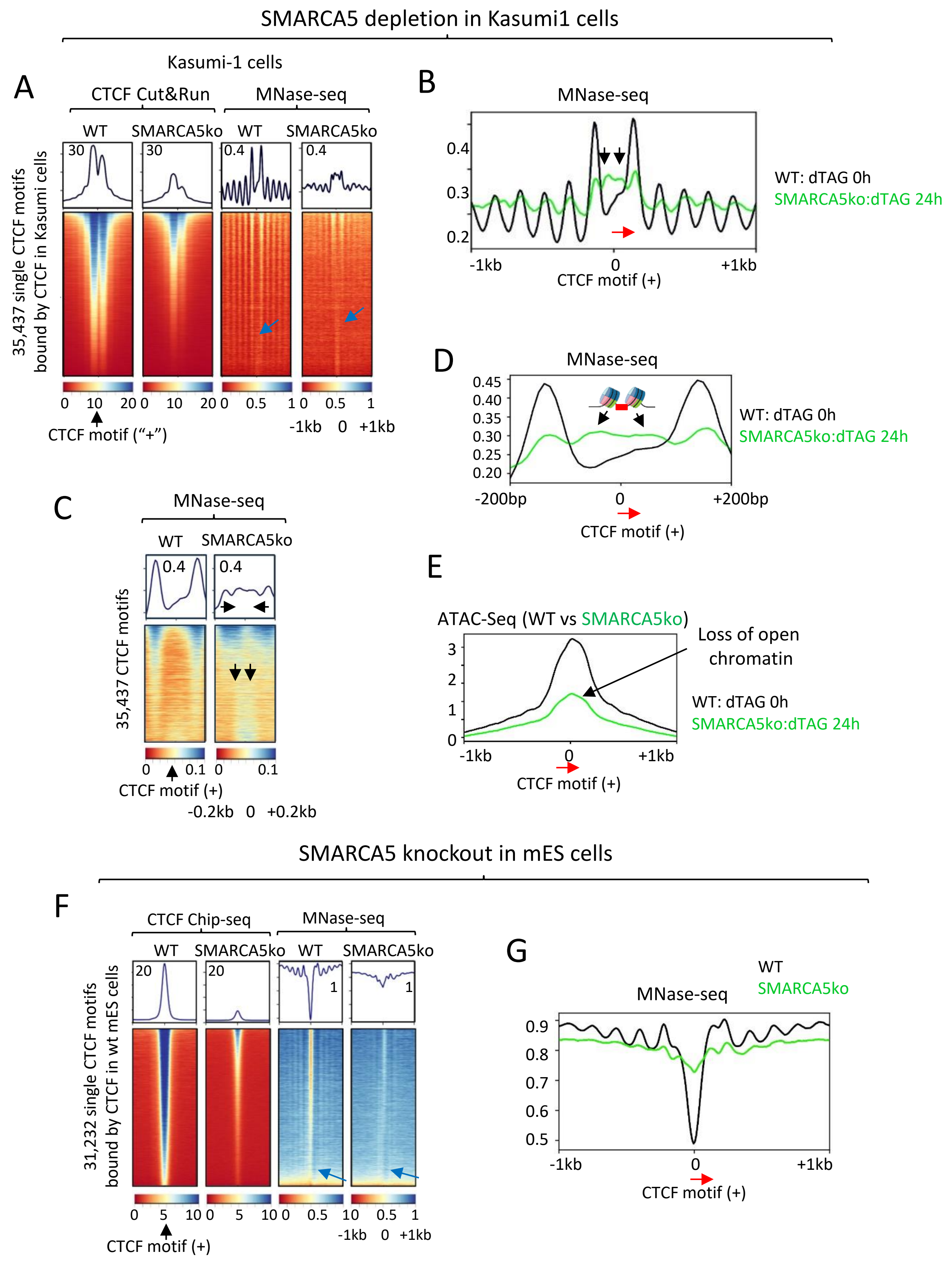

**Supplementary Figure S8. SMARCA5 degradation in Kasumi-1 and mES cells results in a loss of CTCF occupancy, accompanied by disrupted nucleosome positioning and decreased chromatin accessibility at CTCF binding sites. (A–E)** CTCF occupancy and chromatin status in Kasumi-1 cells following dTAG-47-mediated degradation of endogenously modified SMARCA5 protein using a small molecule proteolysis-targeting chimera (PROTAC). Kasumi-1 cells were either not treated with dTAG-47 (0h) or treated for 24 hours (24h, SMARCA5 knockout). **(A)** Heatmaps displaying CTCF occupancy (CUT&RUN) and nucleosome positioning (MNase-seq), centered on CTCF-bound motifs (in plus orientation) mapped by ChIP-seq in Kasumi-1 cells. The CTCF priming nucleosome (CPN) is indicated by blue arrows. **(B, C, D)** MNase-seq profiles showing nucleosome positioning at CTCF-bound motifs. Profiles compare untreated cells (0h, black) with cells treated with dTAG-47 for 24 hours (green, SMARCA5 knockout) within windows of 2,000 bp **(B)** or 400 bp **(C, D).** Flanking CTCF nucleosomes are indicated by black arrows. (**E)** ATAC-seq tag density centered at CTCF motifs in the plus orientation, comparing chromatin accessibility between untreated cells (0h, black) and cells treated with dTAG-47 for 24 hours (green, SMARCA5 knockout). **(F–G)** CTCF occupancy and chromatin status in mouse embryonic stem (mES) cells with SMARCA5 knockout. **(F)** Heatmaps showing CTCF occupancy and nucleosome positioning (MNase-seq), centered on CTCF-bound motifs mapped by ChIP-seq in mES cells. The CTCF priming nucleosome (CPN) is indicated by blue arrows. **(G)** MNase-seq profiles of nucleosome positioning at CTCF-bound motifs in the plus orientation in mES cells, comparing wild-type (WT, black) and SMARCA5 knockout (green) cells.

### Supplementary Figure S9

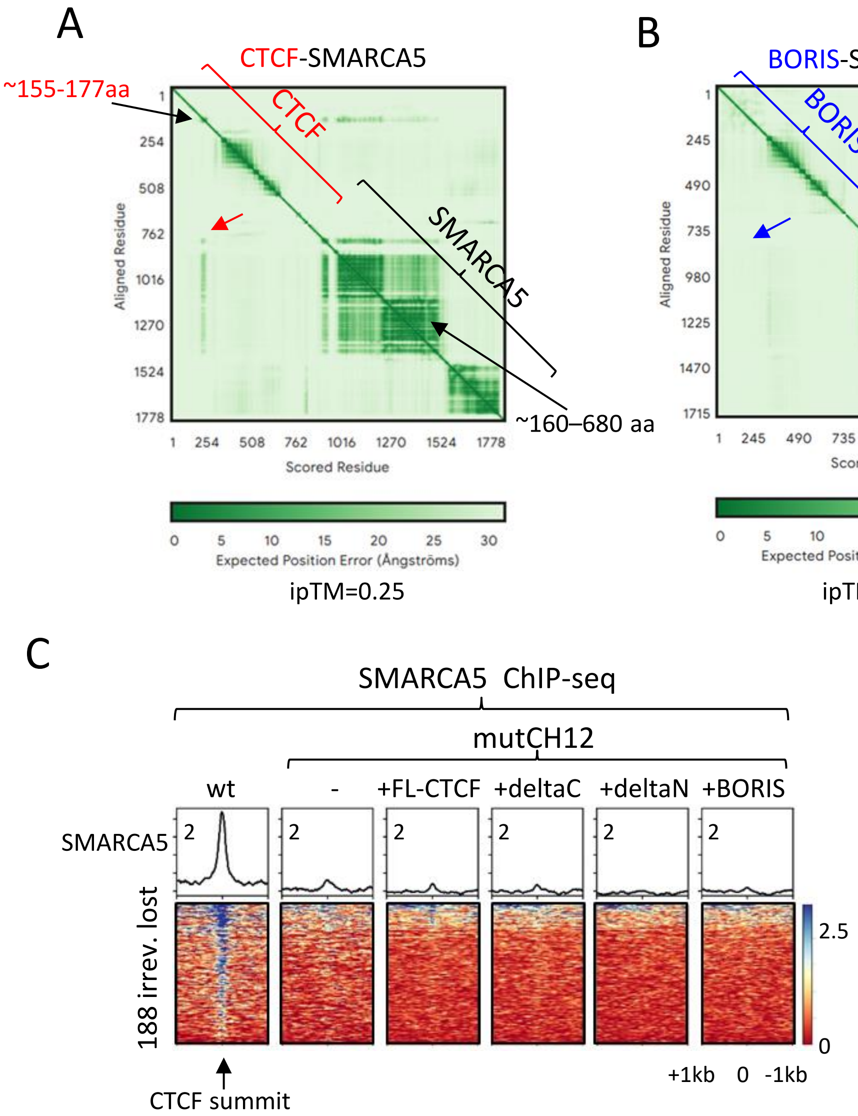


**Supplementary Figure S9. CTCF interaction with SMARCA5. (A, B)** Interactive PAE (Predicted Aligned Error) viewer plots highlighting the predicted interactions between CTCF **(A)** or BORIS **(B)** and SMARCA5 by interface predicted template modelling (ipTM). **(A)** PAE heatmap showing predicted interactions between CTCF and SMARCA5. Red arrow indicates potential interactions between the N-terminal domain of CTCF (~155–177 aa) and the P-loop-containing nucleoside triphosphate hydrolase domain of SMARCA5 (~160–680 aa). **(B)** PAE heatmap of BORIS and SMARCA5 as a control for CTCF. The blue arrow indicates the absence of predicted interaction between BORIS and SMARCA5. **(C)** Heatmaps of SMARCA5 ChIP-seq data across 199 irreversibly lost CTCF binding sites.

### Supplementary Figure S10

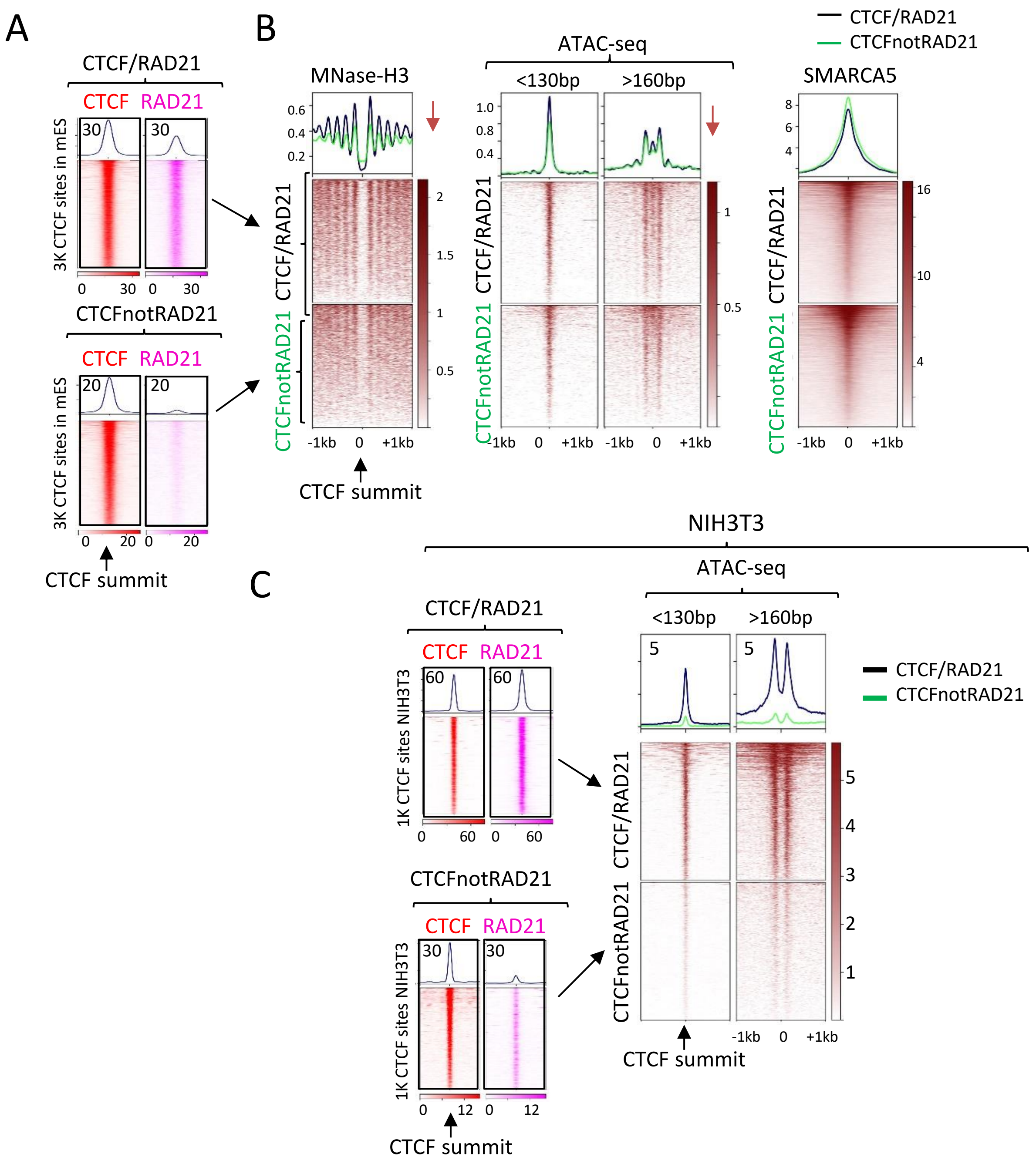


**Supplementary Figure S10. The comparison of chromatin accessibility at CTCF sites enriched with or depleted of cohesin in mES and NIH3T3 cells. (A)** Heatmap of CTCF (red) and RAD21 (pink) ChIP-seq occupancy at 3,000 (3K) CTCF/RAD21 and 3K CTCFnotRAD21 binding sites in mES cells, centered at the summit of CTCF peaks. **(B)** Heatmap showing MNase-H3 nucleosome positioning, ATAC-seq data (with <130 bp reads representing the CTCF footprint and >160 bp reads for nucleosome positioning), SMARCA5 ChIP-seq at CTCF/RAD21 (black) and CTCFnotRAD21 (green) binding sites in mES cells. The data in panel (B) correspond to the heatmap presented in panel (A). **(C)** Left: Heatmap of CTCF (red) and RAD21 (pink) ChIP-seq occupancy at 1,000 (1K) CTCF/RAD21 and 1K CTCFnotRAD21 binding sites in NIH3T3 cells, centered at the summit of CTCF peaks. Right: Heatmap showing ATAC-seq data at CTCF/RAD21 (black) and CTCFnotRAD21 (green) binding sites in NIH3T3 cells.

### Supplementary Figure S11

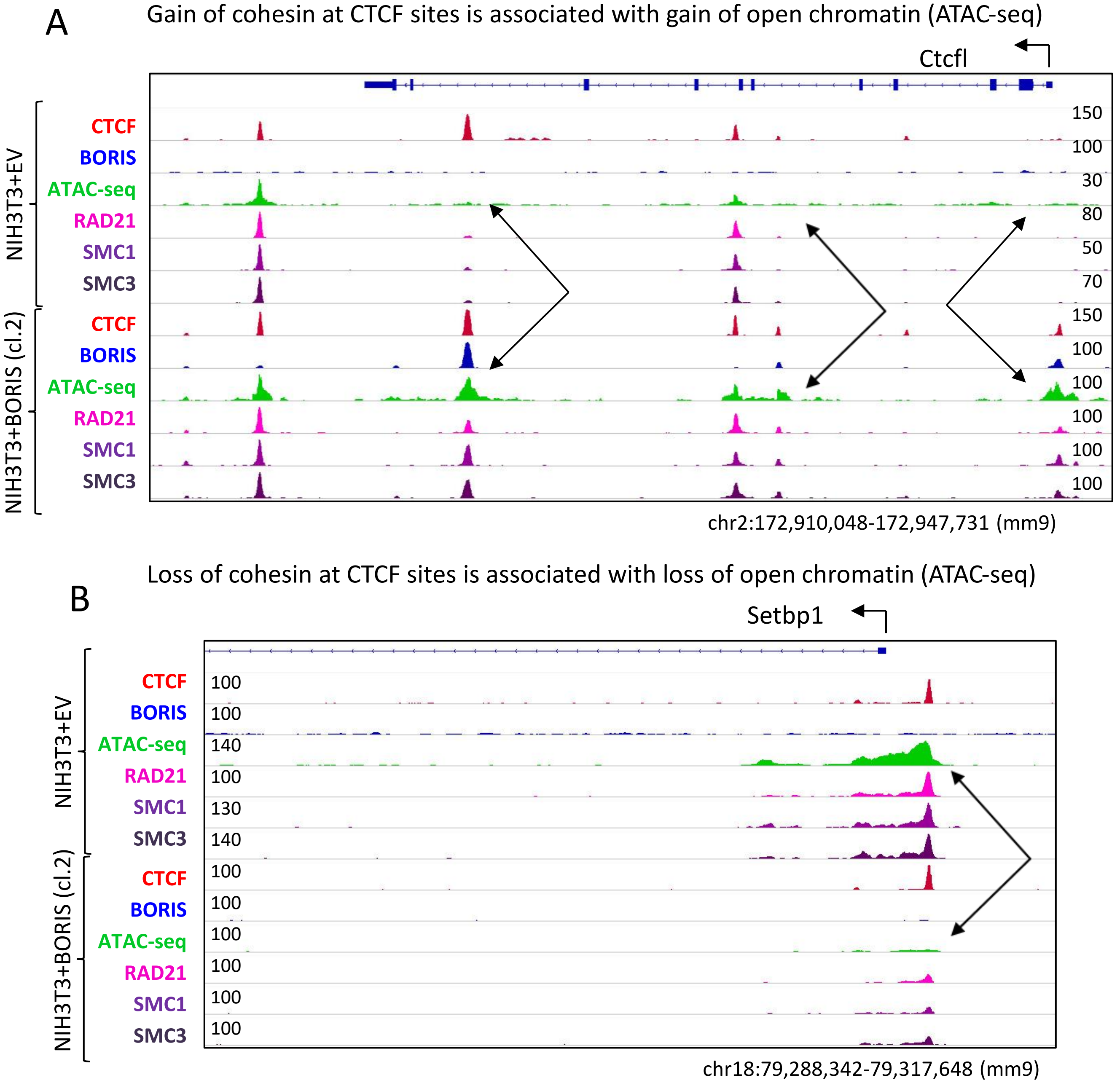


**Supplementary Figure S11. Gain or loss of cohesin occupancy at CTCF sites is accompanied by corresponding gain or loss of chromatin accessibility around CTCF sites, respectively. (A, B)** Genome browser views of CTCF (red), BORIS (blue), chromatin accessibility (ATAC-seq, green), and cohesin subunits (RAD21, SMC1, and SMC3) in NIH3T3 cells stably expressing either Empty Vector (EV, NIH3T3+EV) or BORIS (stable single-cell clone #2, NIH3T3+BORIS) as per PMID: 38297316. **(A)** Examples of CTCFnotRAD21 (cohesin-depleted) sites converted into CTCF/RAD21 (cohesin-enriched) sites in NIH3T3+BORIS cells compared to NIH3T3+EV cells (indicated by black arrows). The gain of cohesin at CTCF sites is accompanied by an increase in chromatin accessibility around these sites. **(B)** Example of a CTCF/RAD21 site converted into a CTCFnotRAD21 site in NIH3T3+BORIS cells compared to NIH3T3+EV cells (indicated by black arrows). The loss of cohesin at CTCF sites is accompanied by a reduction in chromatin accessibility around these sites.

### Supplementary Figure S12

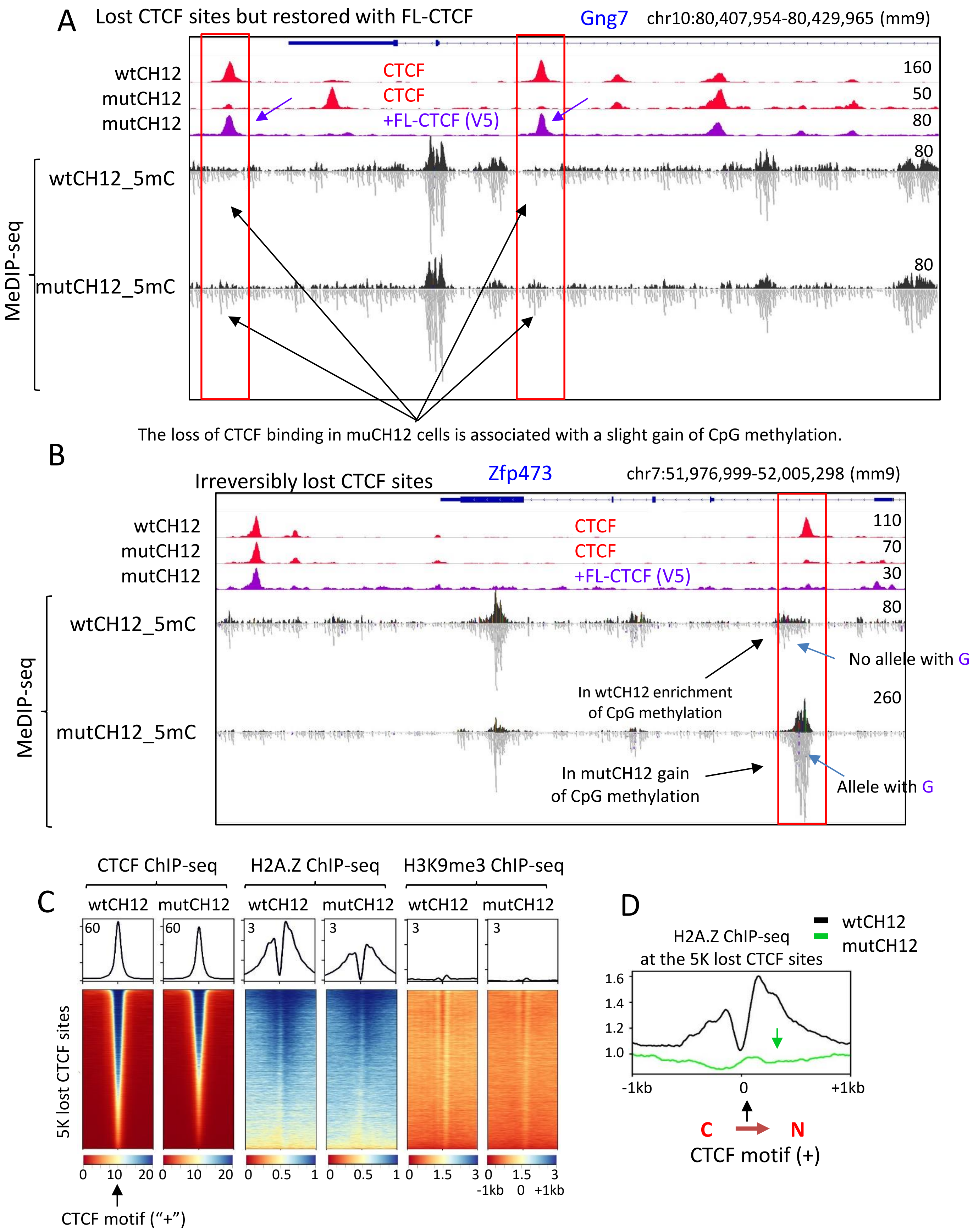

**Supplementary Figure S12**. **Loss of CTCF binding in mutCH12 cells results in a gain of CpG methylation at some CTCF sites, but not all. (A, B)** Genome browser view displaying CTCF (red) and ectopically-expressed FL-CTCF-V5 (purple) ChIP-seq data, along with 5mC MeDIP-seq data in wtCH12 cells and mutCH12 cells. The status of CTCF occupancy restoration via ectopic expression of FL-CTCF (V5-Tag antibodies) in mutCH12 cells is marked by purple arrows. Black arrows highlight changes in CpG methylation at lost CTCF sites in wtCH12 versus mutCH12 cells. **(A)** Example of CTCF sites that were lost and then restored with the ectopic expression of FL-CTCF in mutCH12 cells. These sites exhibit a slight gain of CpG methylation, consistent with the findings presented in main Fig. 6B (middle panel). **(B)** Example of a CTCF site that was lost and not restored with ectopic expression of FL-CTCF in mutCH12 cells. Hemimethylation at this CTCF site is indicated by light blue arrows: a significant gain of CpG methylation is observed in mutCH12 cells, evidenced by the presence of MeDIP-seq reads containing an additional "G" in one allele (indicated by small purple dots in the reads). This extra "G" is absent in MeDIP-seq reads mapped in wtCH12 cells, suggesting that only one allele carries partial CpG methylation at this CTCF site in wtCH12 cells, and both alleles are methylated in mutCH12 cells. **(C)** Heatmaps of CTCF, H2A.Z, and H3K9me3 ChIP-seq data in wtCH12 versus mutCH12 cells at the 56K remaining CTCF sites. **(D)** Profile of H2A.Z ChIP-seq data at the 5K lost CTCF sites in wtCH12 cells (black) versus mutCH12 cells (green), centered at CTCF motifs. The green arrow highlights a loss of H2A.Z in mutCH12 cells compared to wtCH12 cells.
